# The genome of the early diverged amphioxus, *Asymmetron lucayanum*, illuminates the evolution of genome architecture and gene repertoires in cephalochordates

**DOI:** 10.1101/2025.07.21.665930

**Authors:** Yifan Ren, Zepu Miao, Che-Yi Lin, Ludong Yang, Huihui Li, Linda Z. Holland, Sung-Jin Cho, Jr-Kai Yu, Jia-Xing Yue

**Affiliations:** State Key Laboratory of Oncology in South China, Guangdong Key Laboratory of Nasopharyngeal Carcinoma Diagnosis and Therapy, Guangdong Provincial Clinical Research Center for Cancer, Sun Yat-sen University Cancer Center, Guangzhou, China; Institute of Cellular and Organismic Biology, Academia Sinica, Taipei, Taiwan; Marine Biology Research Division, Scripps Institution of Oceanography, University of California, San Diego, USA; Department of Biological Sciences and Biotechnology, Chungbuk National University, Cheongju, Republic of Korea; Marine Research Station, Institute of Cellular and Organismic Biology, Academia Sinica, Yilan, Taiwan

## Abstract

Cephalochordates (amphioxus or lancelet) are considered as living proxies for ancestral chordates due to their key phylogenetic position and slow evolutionary rate. The genomes of living amphioxus thus can help to reveal the genetic basis shaping the evolutionary transition from invertebrates to vertebrates. To gain a comprehensive understanding of the genome architecture in amphioxus, here we generate a chromosome-anchored genome assembly for *Asymmetron lucaynum*, representing the earliest diverging cephalochordate genus. Our results show that *Asymmetron* has an enlarged genome compared to those of the other four cephalochordate genomes decoded so far (all in the genus *Branchiostoma*), causing by pervasive expansions of inter-genic transposable elements (TEs). Nevertheless, both macrosynteny and microsynteny remain highly conserved between *Asymmetron* and *Branchiostoma*, enabling reconstruction of the ancestral genomic architecture of the Cephalochordate lineage for tracing genome evolutionary process during Deuterostome and Chordate diversification. By coupling developmental transcriptomic analyses, we further show that purifying selection and constraints on co-transcriptional regulation may have contributed to the maintenance of the conserved microsynteny blocks among cephalochordate species. We also examine the evolutionary history of *Hox* cluster in cephalochordates and vertebrates, and identify species-specific inversions and TE invasions at this important locus in both *Asymmetron* and *Branchiostoma*. Finally, we survey key gene families involved in both innate and adaptive immunity (e.g., *TLR*, *NLR*, *MHC*, and *RAG*) and uncover their plausible ancestry and evolutionary dynamics in chordates. Taken together, our findings illuminate the genome and gene evolution of cephalochordates and provide valuable resources for understanding the early evolution of chordates and the origin of vertebrates.

## Introduction

The chordate subphylum Cephalochordata (amphioxus or lancelets) diverged from other chordates (vertebrates and urochordates) more than 550 million years ago (MYA)^1^. As “living fossils”, their body plans strongly resemble that of Cambrian fossils such as *Pikaia* and, except for lack of paired eyes, are highly similar to chordate fossils as *Haikouella*^2–4^. Compared to modern vertebrates, they also share many common characteristics such as a dorsal, hollow nerve cord, a notochord, pharyngeal gill slits, and post-anal tails^5^. There are three known genera of cephalochordates: *Asymmetron*, *Epigonichthys* and *Branchiostoma*, with *Asymmetron* being the earliest diverged lineage with the two others radiated more recently^6,7^. To date, the genomes of four *Branchiostoma* species (i.e. *B. floridae* [Bfl]*, B. belcheri* [Bbe], *B. lanceolatum* [Bla], and *B. japonicum* [Bja]) have been sequenced and assembled. Their genome sizes range from ∼383 Mb (*B. japonicum*) to 490 Mb (*B. floridae*), and their chromosome numbers vary between 18 (*B. japonicum*) and 20 (*B. belcheri*) ^8–13^.

Compared to vertebrates, cephalochordates have both simpler morphology and genomes and retain many features likely present in the ancestral chordate. For example, cephalochordates have homologs of the vertebrate forebrain, midbrain and hindbrain, hinting at a deep chordate origin of the vertebrate head^5,14–16^. Although cephalochordates lack the migrating neural crest characteristic of vertebrates, their ectodermal cells abutting the neural plate migrate over it as a sheet and express many of the genes expressed in vertebrate neural crest, suggesting a common evolutionary origin^17^. Moreover, since cephalochordates have not experienced the two-rounds of whole-genome duplications (2R-WGDs) that occurred during early vertebrate evolution, their genome architectures are more streamlined with much less redundancy. One prominent example is the *Hox* cluster, a set of colinearly arranged homologous genes specifying the anterior-posterior segment identity during early embryo development^18^. They were firstly discovered in the fruit fly^19–22^ and were subsequently found in many invertebrates and vertebrates (including human) with similar colinear genomic arrangement and expression patterns^23^. Like the *Drosophila* genome, those of cephalochordates possess a single *Hox* cluster^24^, in contrast to the four cluster configuration found in most vertebrates due to 2R-WGDs. Initial comparisons of the *B. floridae* genome and those of several vertebrates revealed strong macrosynteny conservation between cephalochordates and vertebrates and suggested that the last common ancestor of chordates likely possessed 17 ancestral linkage groups (ALGs)^8,11^. Subsequent studies including additional *Branchiostoma* amphioxus genomes as well as several deuterostome outgroups further suggested the ancestral chordate likely had 23 of 24 putative bilaterian ALGs^13,25–27^. In addition, while cephalochordates, like other invertebrates, only have an innate immune system but lack the adaptive immunity, they do have some important molecular building blocks of adaptive immunity, such as proto-MHC genes^28,29^ and proto-RAG transposons^30–32^. Together, these unique features make the cephalochordates the best living proxy for ancestral chordates before the appearance of vertebrates^33,34^.

Previous studies of cephalochordate genomes mostly focused on *Branchiostoma* species, with only a very few studies considering the genomes of *Asymmetron* and *Epigonichthys*. Morphologically, *Asymmetron* and *Epigonichthys* closely resemble *Branchiostoma* except that *Branchiostoma* has two rows of gonads on both sides of their bodies, whereas the other two each possess only a single row of gonads on the right side. Surprisingly, *Asymmetron* and *Branchiostoma* can hybridize, although only a few of the hybrids with *Branchiostoma* mothers and *Asymmetron* fathers can complete metamorphosis^35^. Although nothing is known of genes and development in *Epigonichthys,* some progress has recently been made on *Asymmetron lucayanum* since an accessible population was found in Bimini, Bahamas where it was originally described (in 1893)^36,37^. Our comparative transcriptomic analyses of *A lucayanum* has revealed a very slow rate of molecular evolution of its protein-coding genes, suggesting *Asymmetron* as optimal for studying the early evolution of chordates^7^. These transcriptome data, combined with whole-genome shotgun sequencing data, also shed light on *Asymmetron’s* ultra-conserved regions (UCRs)^38^, primordial germ cell (PGC) formation, and genes encoding for fluorescent proteins^38–41^.

For a comprehensive understanding of genome evolution in chordates, we used PacBio long-read sequencing and Hi-C technologies to generate a chromosome-level genome assembly for *A. lucayanum.* Comparisons to the genomes of the four *Branchiostoma* species revealed the intra-subphylum diversity of genome evolution in cephalochordates. We then compared chromosomal synteny, genomic loci, and gene families within cephalochordates, as well as to those of several representative invertebrate and vertebrate genomes. Our study showed that *Asymmetron* has a much larger genome than those of *Branchiostoma* species due to more transposable elements. Even so, evolution of its protein-coding regions is as slow as in other cephalochordates. Macro- and microsynteny is highly conserved between genomes of *Asymmetron* and *Branchiostoma*. Despite lineage-specific inter-chromosomal fusions in both genera, we were able to infer that the common ancestor of all cephalochordates likely had 20 ALGs, with the previously defined R bilaterian ALGs lost before the divergence of *Asymmetron* and *Branchiostoma*. We also found coordinated gene expression for genes enclosed in microsynteny blocks, reflecting possible influence of natural selection on the evolution of genomic organization. Finally, using cephalochordates as the anchor point, we examined the evolution of some key gene families and loci underlying metazoan development and immunity before and after the 2R-WGDs. For this, we focused on both the widely conserved *Hox* transcription factor genes and the more dynamically evolved genes functioning in innate and adaptive immunity. We provided a high-resolution view on the evolutionary history of the *Hox* genes in cephalochordates and vertebrates while unrevealing species-specific inversions and transposable element invasions at their residing locus (the *Hox* cluster) in both *Asymmetron* and *Branchiostoma*. Meanwhile, our genome-wide survey of some key immune genes (e.g., *TLR*, *NLR*) and their prototypes (e.g., *proto-MHC* and *proto-RAG*) illuminated their evolutionary dynamics and history during the transition from invertebrates to vertebrates.

## Results

### Chromosome-level genome assembly of *A. lucayanum*

Using PacBio long-read and chromatin conformation capture sequencing (both *in vitro*^42^ and *in vivo*^43^), we generated a chromosome-level reference assembly for *A. lucayanum* (Alu) with haplotype reconciliation. The chromosome-anchored assembly size is 677.4 Mb (distributed across 18 chromosomes), with a weighted median scaffold length (N50) of 36.0 Mb (Figure 1a; Table S1–S3). This agrees with the size of our initial diploid assembly (∼1.3–1.4 Gb) (Table S2) and with the genome size estimate based on read *k-*mers (Figure 1b). This substantially larger genome size of *Asymmetron* compared to that of *Branchiostoma* is in part due to its having 2–3 times more repetitive sequences (REs), (i.e., 292.0 Mb in *A. lucayanum* vs 108.0–157.0 Mb in *Branchiostoma* species). These are mainly transposable elements (TEs) in both intronic and intergenic regions (Figure 1c, d; Table S4). Among different classes of TEs, inverted terminal repeat TEs (TIR-TEs) are abundant in *Asymmetron* (Figure S1). While their expansion time is tricky to estimate, we found the expansion of long-terminal-repeat TEs (LTR-TEs) occurred relatively recently (<1 MYA) in *A. lucayanum* (Figure S2). By integrating Iso-Seq and RNA-Seq data, we annotated 35,203 protein-coding genes, 94.3% (33,199) of which are located on assembled chromosomes (Figure S3; Table S5–S7). The size of the coding region is comparable between the two genera, i.e., 46.9 Mb in *Asymmetron* and 40.1–47.3 Mb in *Branchiostoma*. A BUSCO score-based analysis of the assembly and annotation of the *A. lucayanum* genome indicated that both are of high quality (98.1% and 94.4% respectively)—on a par with the existing chromosome-level assemblies of the four *Branchiostoma species*.

**Figure 1.**
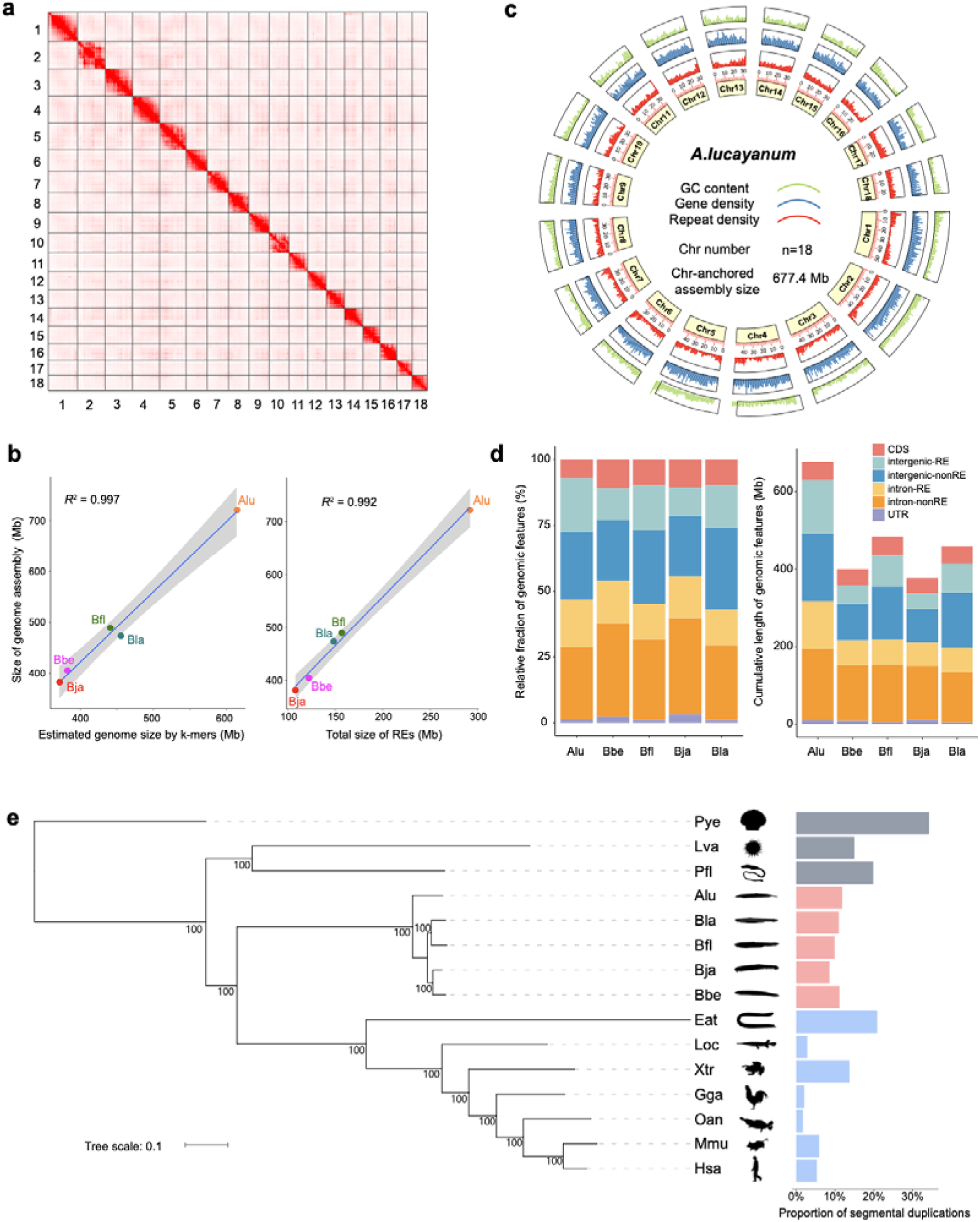
Chromosome-level genome assembly and annotation of *A. lucayanum*. **a**, Hi-C contact heatmap of the *A. lucyanum* reference haplotype assembly. b, Correlations among the assembled genome size (y-axis), the *k*-mer-estimated genome size, and the assembled repetitive sequence size. c, Circos plot of the genome-wide distribution of different genomic features (e.g., GC%, genes, and repeats) along the *A. lucyanum* genome. d, Relative and absolute abundance of different genomic features (e.g., CDSs, intron, and intergenic regions) across the 5 cephalochordate genomes. e, The maximal likelihood phylogeny of cephalochordates in the context of chordate evolution, with the fraction of their genomes involved in segmental duplications further displayed in parallel. Alu: *Asymmetron lucayanum*; Bbe: *Branchiostoma belcheri*; Bfl: *Branchiostoma floridae*; Bja: *Branchiostoma japonicum*; Bla: *Branchiostoma lanceolatum*; Eat: *Eptatretus atami*; Gga: *Gallus gallus*; Hsa: *Homo sapiens*; Loc: *Lepisosteus oculatus*; Lva: *Lytechinus variegatus*; Mmu: *Mus musculus*; Oan: *Ornithorhynchus anatinus*; Pfl: *Ptychodera flava*; Pye: *Patinopecten yessoensis*; Xtr: *Xenopus tropicalis*.

### Universally slow evolution of protein coding genes in cephalochordates

To further explore genome evolution in cephalochordates, we collected all available chromosome-level assemblies of the four *Branchiostoma* species (*B. belcheri, B. floridae, B. japonicum*, *B. lanceolatum*) and their transcriptomes (Table S3, S5) and reannotated their genomes using the same genomics pipeline that we used for *A. lucayanum.* In addition, we further incorporated three more invertebrates (scallop, sea urchin, acorn worm) and seven vertebrates (human, mouse, platypus, chicken, frog, spotted gar, brown hagfish). We built the phylogeny of all included species based on 947 single-copy orthologous genes shared amongst all of them. Relative to the common ancestor of chordates, we observed universally shorter branch lengths for the five cephalochordate species compared with their vertebrate counterparts, reinforcing the idea of slow evolution of protein coding genes in cephalochordates^7,8^. In the same vein, *A. lucayanum* and *B. japonicum* showed the shortest branch lengths, reflecting an even slower molecular evolution of their protein coding genes. In contrast, both *Asymmetron* and *Branchiostoma* cephalochordates showed a higher level of segmental duplications compared with vertebrates (Figure 1e). This agrees with previous reports for *Branchiostoma* species^12,13^ and suggests segmental duplication as a shared driving force shaping the evolution of cephalochordate genomes. By coupling this phylogeny with fossil-calibrated molecular dating, we then estimated a divergence time between *Asymmetron* and *Branchiostoma* as about 88.0 MYA, with the speciation of the four *Branchiostoma* species occurring more recently ∼32.0–49.4 MYA, in contrast to the deep pre-Cambrian divergence between cephalochordates and vertebrates (Figure S4; Table S9). Based on these divergence times and intra-cephalochordate synonymous substitution rates summarized from single-copy orthologs (Figure S5), we estimated a cephalochordate-specific mutation rate of 4.65×10^-9^ substitution/site/year, highly concordant with a recent estimate (4.41×10^-9^) by pedigree sequencing in *B. floridae*^44^.

### Karyotype evolution of cephalochordates

The chromosome-level genome assemblies and annotations of both *Asymmetron* and *Branchiostoma* species revealed a general one-to-one chromosomal correspondence between the two genera despite their 88-MYA divergence. The main difference is two additional *Asymmetron*–specific inter-chromosomal fusions in chromosomes 1 and 3 (Figure 2a). By matching with the 24 bilaterian ancestral linkage groups (ALGs) as well as the 29 Bilaterian, Cnidarian and Sponge (BCnS) ALGs previously defined^25^, we reconstructed the history of karyotype evolution in cephalochordates. These results extended the previously proposed 20 ALGs for the ancestral *Branchiostoma* to the common cephalochordate ancestor. These 20 ALGs correspond to 23 of 24 bilaterian ALGs and 28 of 29 BCnS ALGs^13^. Starting from the 24 bilaterian ALGs, our analysis indicates that the loss of the R ALG occurred during the early evolution of chordates, predating the split between *Asymmetron* and *Branchiostoma*. In addition, we identified three ancestral inter-chromosomal fusion events (A1⊗A2, C1●J2 and I●O1) that occurred before the *Asymmetron*-*Branchiostoma* divergence, among which the J2/C1 ALG fusion showed distinct patterns in *Asymmetron* and *Branchiostoma* (Figure 2b). Specifically, *Asymmetron* may have undergone additional inversion events resulting in a “J2-C1…C1-J2” pattern, or alternatively *Branchiostoma* might have undergone J2●C1 (J2 insert into C1) while *Asymmetron* could have experienced C1●J2 (C1 insert into J2). Conservation of synteny around the J2/C1 fusion site compared to the pre-fusion condition in the hemichordate *Ptychodera flava* revealed additional reshuffling between the J2 and C1 ALGs in *A. lucayanum* compared to *Branchiostoma* species (Figure S6). Except for *B*. *belcheri*, which lacks inter-chromosomal fusions, the other cephalochordate species each had 1–2 such fusions (Table S10). A sliding-window-based genomic scan for these fusion sites showed peaks of segmental duplications at all seven fusion sites that we examined (Figure S7), suggesting a possible intrinsic link between these two types of chromosomal rearrangements. In comparison, the local distribution of repetitive sequences seems more evenly across the fusion sites. At the global scale, we observed a modest tendency for more segmental duplications and repetitive sequences (but a lower proportion of CDSs) in fusion sites spanning over larger genomic regions (Figure S8), hinting at the possible evolutionary process and consequences of those fusions (see Discussion).

**Figure 2.**
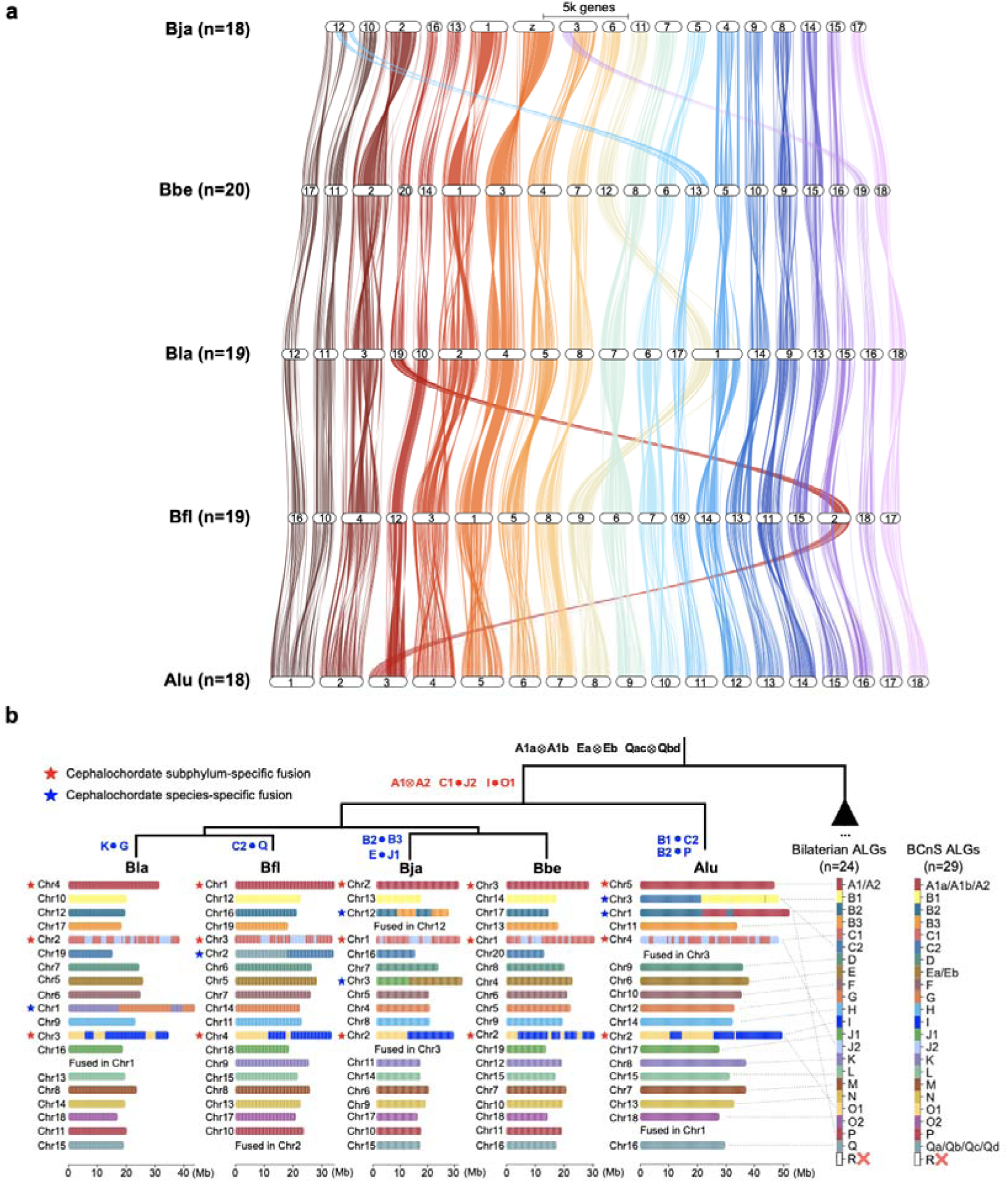
Karyotypes evolution of the five amphioxus species. **a**, Conserved macrosynteny across five cephalochordate species. The chromosomal karyotype of each species is shown horizontally, with colored lines (color-coded based on *A. lucayanum* chromosomes) linking orthologous genes shared among them. b, The reconstructed history of cephalochordate karyotype evolution. Each colored block represents one or more (when they still stay together on the same chromosome in cephalochordates) previously defined Bilaterian (n=24) and BCnS (n=29) ALGs. Two types of chromosomal fusion events (●: fusion without mixing, ⊗: fusion-with-mixing) are indicated on top of the phylogenetic tree.

### Signatures of macro- and microsynteny conservation in chordate evolution

We detected a high degree of macrosynteny (i.e., chromosomal gene colocalization) between *Asymmetron* and several vertebrates and invertebrates (Figure S9). Combined with similar findings made with *Branchiostoma*^8,11,13^, this suggests universally slow evolution of macrosynteny in cephalochordates. Regarding microsynteny (i.e., the conservation of local gene orders), we detected 1,278 pan-cephalochordate microsynteny blocks, each of 3–25 genes, encompassing a total of 6,704 orthologous gene groups (Figure 3a; Table S11). The distribution of block sizes fits best to a log-normal distribution, indicating the existence of multiplicative effects such as selection on top of exponential decay (Figure S10-S11). Consistent with this, genes within these microsynteny blocks show slower rates of molecular evolution (dN and dS) and stronger purifying selection (dN/dS) compared with genes outside the blocks (Figure S12). The in-block correlation of gene-specific measurements of both selection (dN/dS) and expression (TPM; transcripts per million) were also significantly higher than calculations based on reshuffled blocks (Figure 3b, c), suggesting that genes contained in the same microsynteny blocks are inherently more alike in regard to these measurements. By further requiring strict in-block gene collinearity with the human genome, we detected 52 such highly conserved cephalochordate microsynteny blocks (each containing 3–5 genes), indicating strong selection constraints on local gene orders over chordate evolution (Figure S13, Table S12). These 52 cephalochordate-human microsynteny blocks recaptured some developmentally important gene clusters such as the *Hox* genes^21^ and genes patterning the pharynx^45^, suggesting a role of developmentally regulated spatiotemporal transcription in preserving the local gene order within these microsynteny blocks.

**Figure 3.**
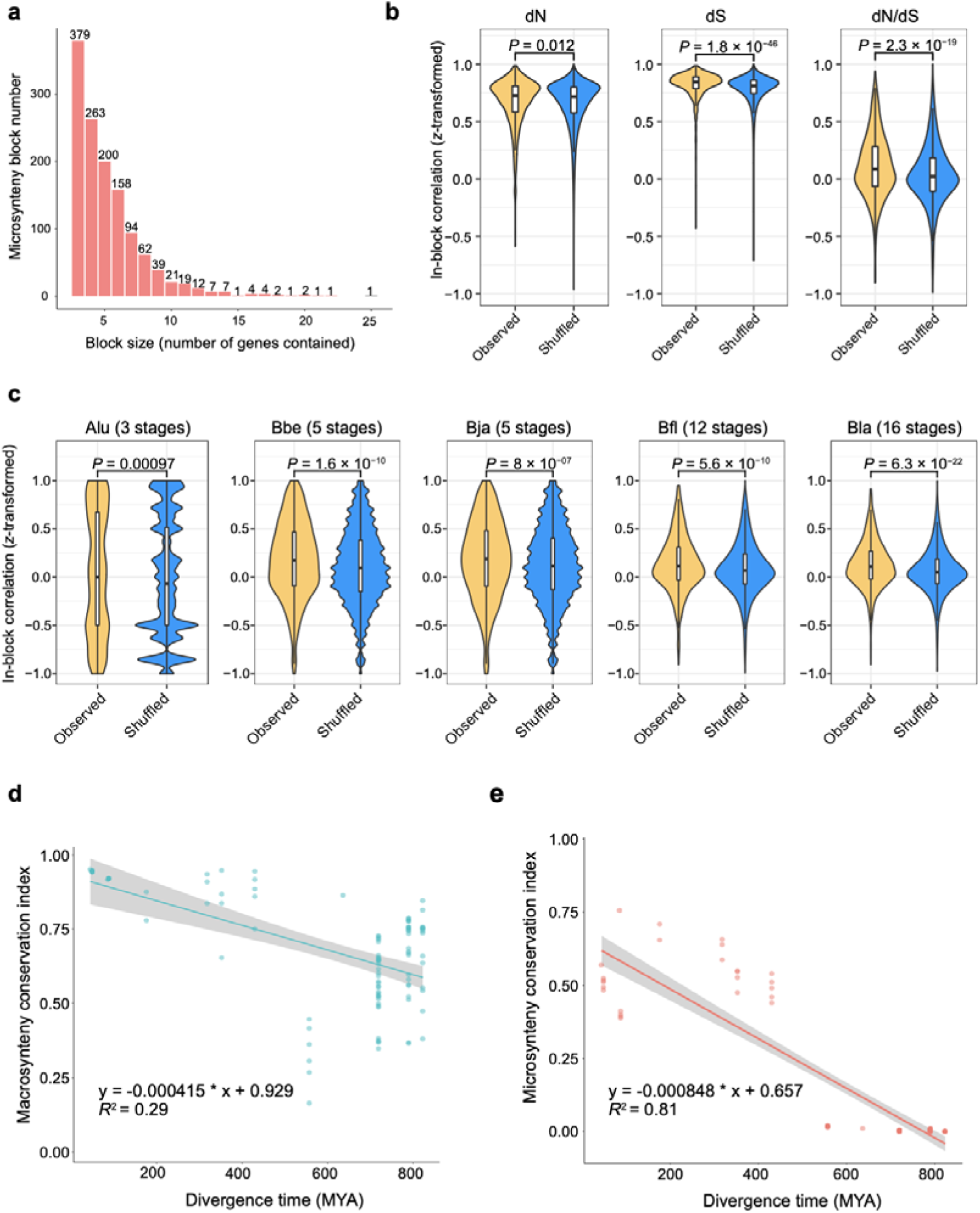
Macrosynteny and microsynteny conservation. **a**, Size-dependent distribution of pan-cephalochordate microsynteny blocks. b, Within-block correlation of molecular evolution metrics (dN, dS, and dN/dS) of genes from the observed cephalochordate microsynteny blocks against those of genes sampled from randomized blocks. The molecular evolution metrics were calculated for all pairwise comparison of the five cephalochordate species. c, Within-block correlation of gene expression metrics (TPM) of genes from the observed cephalochordate microsynteny blocks against those of genes sampled from randomized blocks. This analysis was conducted for the five cephalochordate species separately based on their respective expression data across multiple developmental stages (indicated on top). d-e, Negative correlation between synteny conservation (d: macro-, e: micro-) and divergence time. TPM: Transcripts Per Million. Two-sided Wilcoxon rank-sum test was used for the comparison in panels b-c. Pearson linear regression was performed for panels d-e.

To quantify the degree of conservation of macro- and microsynteny between different lineages, we performed all possible pairwise comparisons among the 7 vertebrates and 8 invertebrates (Table S13, S14). In general, for both vertebrates and invertebrates, macrosynteny exceeds microsynteny while both macro- and microsynteny are greater within each group than between the two groups. Not surprisingly, conservation of both macro- and microsynteny decreases with increasing divergence time between species, with a more rapid decay of the latter (Figure 3d, e). Among the vertebrates, the hagfish (*Eptatretus atami*) genome showed substantially less conservation in both macro- and microsynteny compared with cephalochordates and other vertebrates (Table S13, S14). This is likely due to a cyclostome-specific whole genome duplication (2R_CY_), followed by reshuffling of genes during re-diploidization^46,47^. Likewise, a generally faster decay of macrosynteny relative to other vertebrate lineages was detected for the mouse within vertebrates (Table S13), reflecting its accelerated chromosome evolution.

### Evolutionary history of *Hox* genes in chordates

Like *Branchiostoma*^5,13^*, Asymmetron* has a single *Hox* cluster of 15 genes, suggesting that this is the ancestral cephalochordate *Hox* configuration. Designating hemichordates as the outgroup^48^, we analyzed the orthologous correspondence of *Hox* genes among both *Asymmetron* and *Branchiostoma* cephalochordates and several representative vertebrates. This analysis confirmed the recently proposed one-to-one orthology between the cephalochordate and vertebrate *Hox1–Hox5* genes and *Hox9* genes^13^, while identifying an additional case for their *Hox6* genes (Figure 4a, b; Table S15). Chordate *Hox7* and *Hox8* genes cluster together as immediate phylogenetic neighbors, suggesting their common ancestry from a tandem duplication in the ancestral chordate. Cephalochordate *Hox 10–12* genes collectively correspond to the vertebrate *Hox10*, suggesting that they are derived from cephalochordate-specific duplication events. Cephalochordate *Hox13–15* genes are closely related to the vertebrate *Hox11–14* genes, but their precise correspondence is obscured, likely a reflection of so-called “posterior flexibility”^49^. Specifically, cephalochordate *Hox13* and *Hox14* genes originated from a cephalochordate-specific gene duplication and together correspond to the vertebrate *Hox11* and *Hox12* genes, which are derived from a vertebrate-specific duplication. Cephalochordate *Hox15* is orthologous to vertebrate *Hox13*. We detected no cephalochordate ortholog for vertebrate *Hox14*, which was only found in early-diverged vertebrate lineages such as lamprey, hagfish, shark, and coelacanth^49–53^. Therefore, vertebrate *Hox14* is likely a lineage-specific duplication of vertebrate *Hox13*.

**Figure 4.**
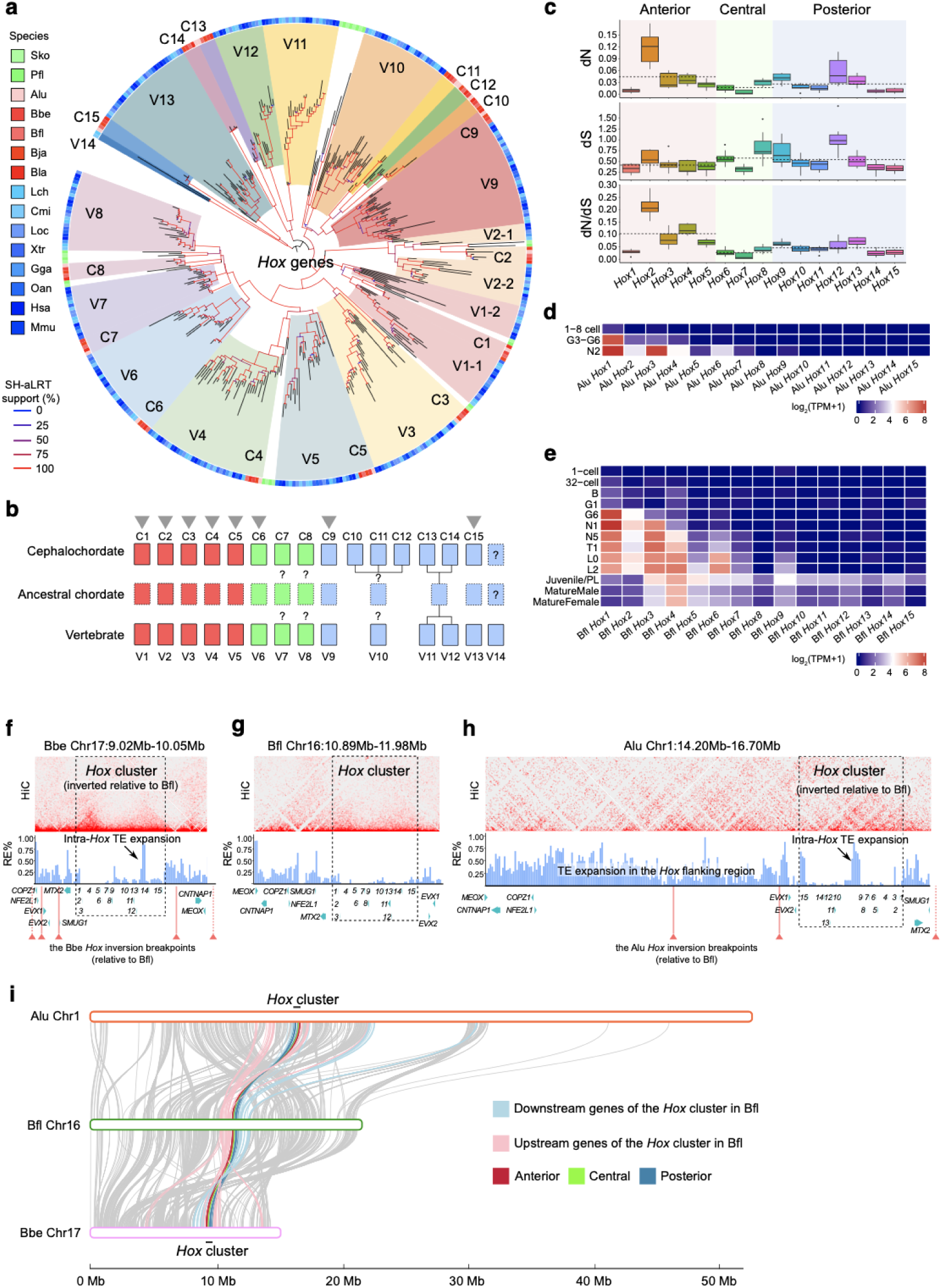
The *Hox* cluster evolution in cephalochordates and vertebrates. **a**, The maximal likelihood tree of *Hox* genes from representative cephalochordates, vertebrates, and hemichordates. b, Proposed model of *Hox* gene evolution in cephalochordates, vertebrates and their putative common ancestor. *Hox* genes with clear one-to-one orthologous relationship are indicated by grey triangles. Dashed boxes indicate putative gene loss events, while question marks denote uncertainty of orthologous relationship. c, The molecular evolution metrics (dN, dS, and dN/dS) of the *Hox* genes in cephalochordates. Dashed line indicated the algorithmic mean value for the corresponding segment (i.e., anterior, central, posterior). d-e, the stage-(d) and tissue-specific (e) patterns of *Hox* gene expression in cephalochordates (Bla). f-h, The local landscapes of 3D genome interaction (HiC) and repetitive elements (RE%, 10-kb window) of the *Hox* cluster in *B. belcheri* (f), *B. floridae* (g), and *A. lucayanum* (h), with the *B. belcheri* and *A. lucayanum Hox* inversion breakpoints (relative to *B. floridae*) further indicated using solid (when the actual breakpoints are in the depicted region) or dashed arrows (when the actual breakpoints go beyond the depicted region). The species-specific RE expansions in *B. belcheri* and *A. lucayanum* are indicated by arrows. i, Distinct *Hox*-flanking inversions of *A. lucayanum* and *B. belcheri* relative to *B. floridae*.

In contrast to the posterior and central *Hox* genes of cephalochordates, the coding sequences of the anterior ones evidently evolved under more relaxed constraints, as shown by their substantially higher dN/dS values (Figure 4c). The same trend holds for the human-chimpanzee *Hox* comparison, suggesting that this may be a more general trend (Figure S14). Such faster molecular evolution of anterior *Hox* genes is potentially due to a relaxation of purifying selection in directing their developmental stage- and tissue-specific expression (Figure 4d, e; Figure S15). We also found that the *A. lucayanum Hox* cluster like those of *B. belcheri*^13^ is inverted relative to those in *B. floridae, B. japonicum*, and *B. lanceolatum*, but has distinct breakpoints (Figure 4f–i). Interestingly, the inferred breakpoints suggest that there were at least two different inversion events in both the *A. lucayanum* and *B. belcheri Hox* clusters, highlighting the surprising structural dynamics of the *Hox* cluster in relation to its flanking genomic regions. In addition, although *Hox* clusters typically have relatively few repetitive elements^54,55^, we found substantial intra-*Hox* TE expansions between *Hox9* and *Hox10* in *A. lucayanum* (Mutator TE) as well as between *Hox13* and *Hox14* in *B. belcheri* (Helitron TE) (Figure 5f–h; Figure S16). Such co-occurrence of *Hox* cluster inversion and intra-Hox TE invasion in both *A. lucayanum* and *B. belcheri* may be non-random. The functional impact, if any, of such co-occurrence remains to be determined in future.

**Figure 5.**
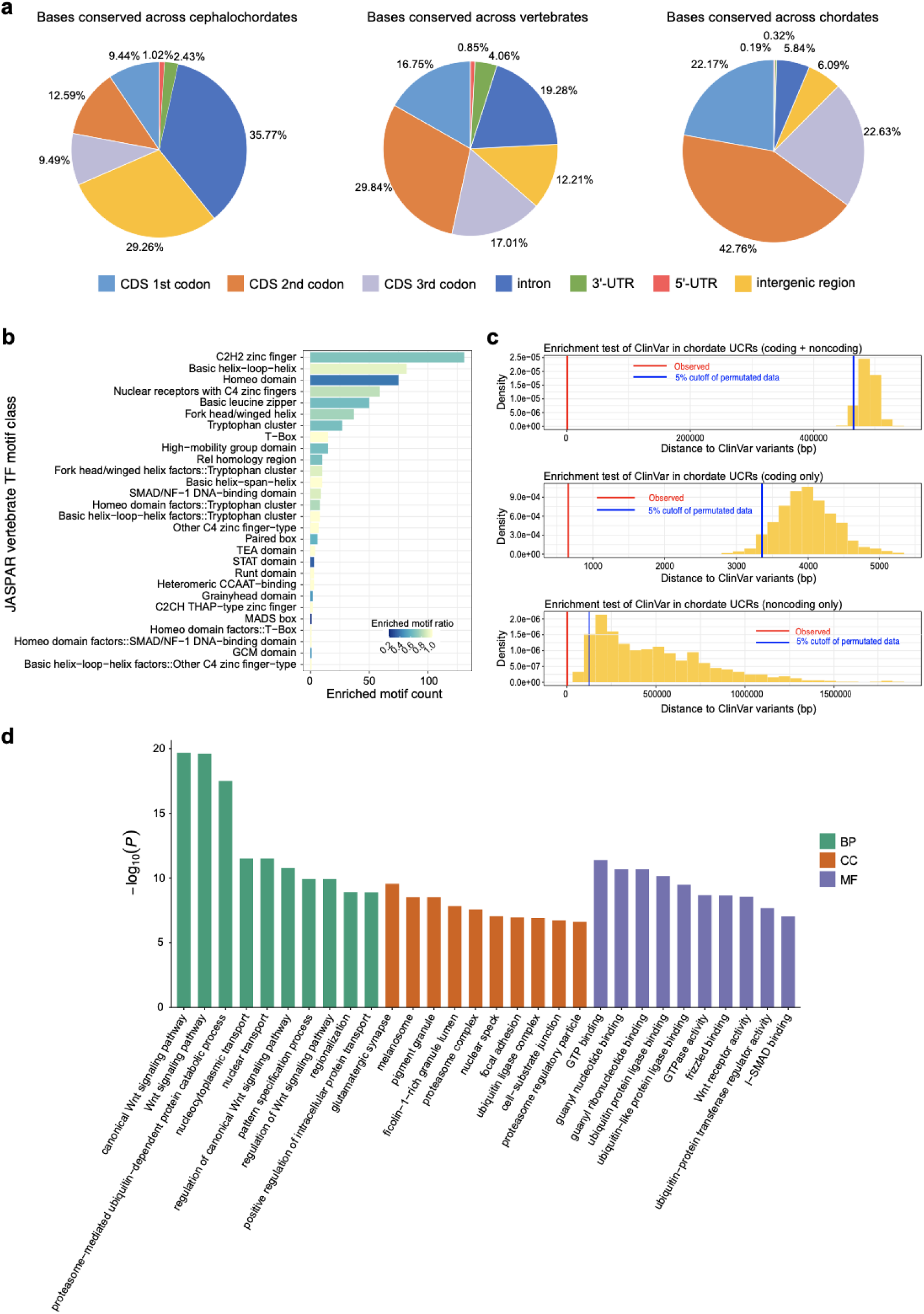
Genomic regions under strong selective constraints. (a). Breakdown pie charts of evolutionarily conserved genomic regions in cephalochordates (left), vertebrates (center), and chordates (right). (b) Enriched vertebrate transcription factor binding motifs in conserved noncoding regions identified for cephalochordates compared with randomized regions. (c) Enrichment of human disease-associated variants in the identified chordate UCRs compared with randomized regions. (d) Gene ontology enrichment of human genes with substantial overlaps (≥50% coding regions) with the identified chordate UCRs. Only top10 results are shown in the figure. BP: biological process, MF: molecular function, CC: cellular component.

### Ultra-conserved regions shared between cephalochordates and vertebrates

In addition to conserved microsynteny, conserved genomic regions in evolutionarily diverged lineages also reflect strong purifying selection. Whole genome alignments revealed 49,178,410 bases conserved between *A. lucayanum* and all four *Branchiostoma* species and 9,553,475 bases conserved among six vertebrates (spotted gar, frog, chicken, platypus, mouse, human) as well as 2,047,162 bases conserved among all eleven chordates (Figure 5a). Throughout these comparisons, we found the second codon position of the CDSs consistently overrepresented compared to the first and third positions among the conserved bases, possibly driven by its crucial impact on the physicochemical properties of codons^56,57^. In addition, there were far more conserved bases in the 3’untranslated regions (UTRs) than in the 5’ UTRs, indicating strong evolutionary constraints on 3’UTRs evidently due to their roles in regulating the localization, stability and translation of mRNAs^58^.

Of the conserved bases among the 5 cephalochordates, 31.52% correspond to CDSs, covering at least 50% of the CDS length of ∼70% of the *A. lucayanum* genes. The remaining 68.48% of conserved bases correspond to noncoding regions such as introns, UTRs, and intergenic regions, suggesting their conserved regulatory functions. To test this hypothesis, we merged these conserved bases into 808,417 pan-cephalochordate conserved regions (see Methods), covering 62.6%–70.4% of open chromatin peaks previously identified in *B. lanceolatum* across multiple developmental stages^10^ and 70.2–83.6% micro-RNAs curated for *B. belcheri* and *B. floridae* in the miRBase database^59^. Of 52 *cis-*regulatory elements previously determined for *Branchiostoma* by transgenic reporter assays^5,10,38,60–79^, 47 (90%) matched the pan-cephalochordate conserved noncoding regions we found (Table S16). Therefore, most of these noncoding regions conserved among both *Asymmetron* and *Branchiostoma* likely have highly similar regulatory functions. Additionally, there were 579 matches to transcription factor binding motifs of vertebrate genes in the JASPAR database^80^. Many of these correspond to C2H2 zinc finger genes, basic helix-loop-helix (bHLH) genes, homeobox genes, and nuclear receptor genes with Z4-fingers (Figure 5b).

Not surprisingly, there are fewer regions conserved between cephalochordates and vertebrates, likely due to their >500 MYA divergence and to rewiring of the gene regulatory networks during and after the vertebrate 2R-WGDs^81^. We used additional synteny-based filters (see Methods) to merge these conserved regions into 18,584 human-genome-projected ultra-conserved regions (UCRs) that are shared amongst all eleven chordates. For the 33 chordate UCRs matched with human noncoding regions, half (16/33) of them are strictly noncoding in all 11 chordate species, with three of these regions harboring known micro-RNA genes (Table S17). These 16 noncoding UCRs included known bilaterian/chordate ultra-conserved noncoding elements associated with *MSX1*, *EBF3*, *HOX4* and *ZNF503* (Bicore2)^38,62^. Our analysis filtered out one known bilaterian ultra-conserved noncoding element associated with *IDI* (Biocore1)^62^ as it was lost in the frog (*Xenopus tropicalis*) genome. In addition, we also identified other noncoding UCRs shared between cephalochordates and vertebrates (Table S17). These included four that are closely spaced along the 3’-UTRs of cephalochordate *MEX3* and vertebrate *MEX3B*. Two of them (UCR10 and UCR11) are directly connected in cephalochordates but are separated in vertebrates (Figure S17). Notably, among the four paralogs of *MEX3* in vertebrates (i.e., *MEX3A, MEX3B, MEX3C, MEX3D*), only *MEX3B* exhibits such striking conservation of the 3’-UTR, suggesting its functional importance. In contrast, the ancestral chordate copy of the noncoding UCR of the vertebrate *PTPRN/PTPRN2* was retained in both post-WGD paralogs, likely constrained by the highly conserved microRNA gene *miR153* that it harbors. Other noncoding UCRs that we identified are associated with genes including *SMAD6*, *MIB1*, *SIX1*, *TNPO1*, and *TPM1* (Table S17).

Although the specific functions of most of these UCRs have yet to be determined, given their strong sequence conservation for >500 million years, there is a high probability that mutations in these UCRs may result in genetic diseases. Indeed, we found that variants associated with human diseases (curated in the ClinVar^82^ database) are considerably enriched in our identified chordate UCRs (Figure 5c). In addition, many of the chordate UCRs that closely match human coding regions (i.e., covering ≥50% CDS region for a given gene), are associated with genes involved in Wnt signaling and protein ubiquitination. (Figure 5d; Table S18). Finally, there was a significant overlap with previously defined clinically important genes. Taken together, the wide conservation of these UCRs likely indicates targets of deep functional constraints in evolution of coding genes in chordates.

### Pan-cephalochordate survey of the repertoire and genomic organization of immune genes

Innate immune systems serve as front-line defenses against pathogens in both plants and animals^83,84^. Their chief components are pattern recognition receptors, most notably the trans-membrane Toll-like receptors (TLRs) and the intracellular NOD-like receptors (NLRs)^85^. In addition to innate immunity, vertebrates also have adaptive immunity, providing a second layer of pathogen defense^86^. While cephalochordates lack adaptive immunity, they do possess homologs of genes mediating adaptive immunity in vertebrates, including the major histocompatibility complex (MHC) and the recombination activating genes (RAGs)^5,31,87,88^.To better understand evolution of these important immune genes in the context of chordate evolution, we performed a comprehensive genome-wide survey of diverse metazoan phyla including both *Asymmetron* and *Branchiostoma* genomes.

### Toll-like receptors (TLRs)

In each of the five cephalochordates, we identified 26–39 *TLR* genes, more than the *TLRs* in all examined vertebrate genomes (e.g., 10 in human and 9 in chicken), although fewer than in some other invertebrates (e.g., 44 in the mollusk *Patinopecten yessoensis*, 57 in the hemichordate *Ptychodera flava*) (Table S19). About one-quarter (e.g., 24.2% in *B. japonicum*) to one-third (e.g., 34.3% in *B. lanceolatum*) of these cephalochordate *TLRs* evidently resulted from tandem duplications (Figure 6b; Figure S18). Notably, cephalochordate *TLRs* fall into two major groups at opposite ends of the *TLR* gene tree (Figure 6a). The upper group (including four *TLR* clades shared by both *Asymmetron* and *Branchiostoma* and five *Branchiostoma*-only *TLRs* clades) is closely clustered with the protostome and non-vertebrate deuterostome *TLR* genes, suggesting that they arose before the protostome-deuterostome divergence. All four *TLRs* that we identified from two urochordate species are present in close adjacency to this upper group (Figure 6a; Figure S19). The lower group (including eight *TLR* clades shared by both *Asymmetron* and *Branchiostoma*, seven *Branchiostoma*-only and one *Asymmetron*-only *TLRs* clade) is more closely related to hemichordate and vertebrate *TLRs*, representing evolutionarily more derived *TLR* genes (Figure S20). Of the previously characterized *TLR* genes, *TLR3*, which senses double-stranded RNAs (dsRNAs), represents the most early diverging *TLR* in the lower group. It is conserved as a single copy in all examined cephalochordates and vertebrates (Figure 6a; Figure S21). Related *TLRs* from hemichordates and echinoderms lie close together but have variable copy numbers, suggesting that *TLR3* is a chordate-specific innovation. There is also an expanded clade of cephalochordate *TLR* genes that is closely related to the bacterial-lipopolysaccharide-sensing *TLR4* (only found in amniotes) and bacterial-flagellin-sensing *TLR5* in vertebrates, hinting that these cephalochordate *TLRs* might well recognize bacterial pathogens (Figure 6a; Figure S21). We also found a clade of cephalochordate *TLRs* closely related to the vertebrate *TLR7*/*8*/*9*. The divergence among *TLR7*, *TLR8* and *TLR9* occurred after the rise of jawed vertebrates, with their pre-triplicated ortholog (*TLR7/8/9*) maintained as a single copy in jawless vertebrates (e.g., lamprey and hagfish) (Figure S22). The vertebrate-specific *TLRs* that were lost in human (*TLR11/1213/21/22*) are related to *TLR7/8/9*. There is no closely related cephalochordate homolog for the vertebrate *TLR1/2/6/10/15*, which predominantly sense lipopeptides.

**Figure 6.**
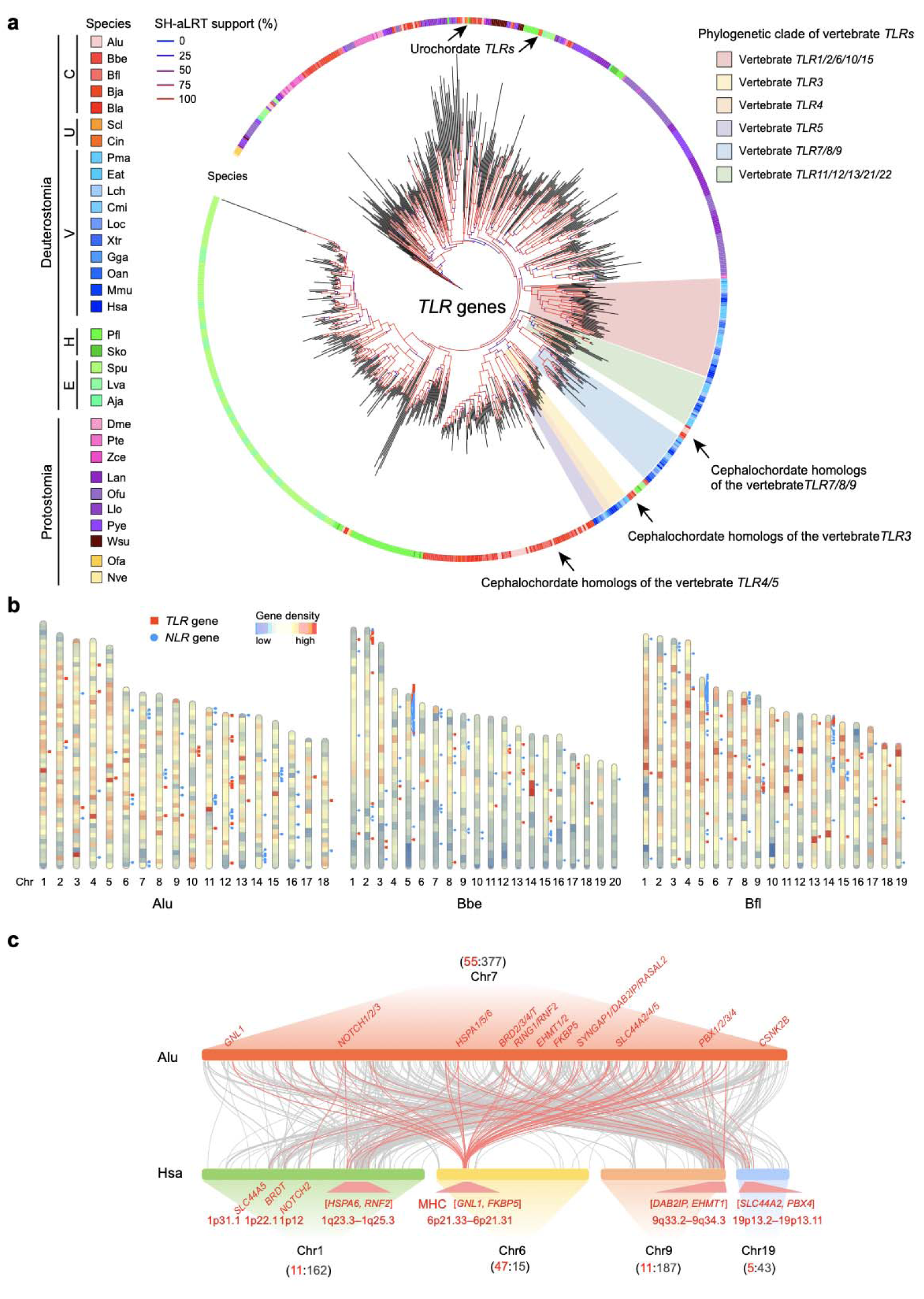
Evolution of immune-related gene and gene families in cephalochordates. (a) Phylogenetic tree of metazoan *TLR* genes. C: Cephalochordata, U: Urochordata, V: Vertebrata, H: Hemichordata, E: Echinodermata. (b) Genome-wide distribution of cephalochordate *TLR* and *NLR* genes in relation to their chromosomes. (c) Macrosynteny conservation between the proto-MHC-carrying *Asymmetron* chromosome 7 and the human chromosomes 1, 6, 9, and 19 with MHC paralogs. The *Asymmetron*-human gene orthology correspondence are indicated by red and grey lines, with those matching with human MHC genes colored in red and the rest colored in grey. Gene count summaries of genes falling into these two categories (red vs. grey) are provided for the *Asymmetron* chromosome 7 and its four homologous chromosomes in human. For each of the four human MHC-homologous blocks (reflected by highly colocalized MHC paralogs), their leftmost and rightmost anchor genes as well as the corresponding cytoband information is further denoted.

### NOD-like receptors (NLRs)

*NLRs* are intracellular receptor genes that sense diverse pathogenic triggers and activate different innate immune responses, including inflammasome formation, PANoptosis induction, and MHC gene expression^89^. NLR proteins are characterized by their tripartite protein domain architectures, including a highly conserved central NACHT domain fused with variable N-terminal (e.g., CARD, Death, DED, HEPN_DZIP3, etc.) and C-terminal domains (e.g., LRR, TPR, etc.). Cephalochordates each have 55–76 NACHT-encoding genes — far more than in most vertebrates (e.g. 25 in human, 8 in chicken) (Figure S23; Table S19). Like for *TLRs*, tandem duplication also accounts for 25.5%–43.4% of cephalochordate *NLRs* species wide (Figure 6b; Figure S18). Some notable examples include clusters of *NLRs* on chromosome 5 in both *B. belcheri* (containing 19 *NLR* genes within a 452.14-kb region) and *B. floridae* (containing 18 NLR genes within a 307.52-kb region). Our phylogenetic analysis of *NLRs* from diverse metazoan lineages suggests a strong pattern of lineage-specific expansion, with *NLRs* from species of the same (sub-)phylum predominantly clustered together (Figure S23). Notably, echinoderms and cnidarians have extensively expanded *NLR* genes (Table S19). In some protostomes and all cephalochordates, additional diversity of *NLR* genes was generated by fusions between some N-terminal domains (e.g., Death, DED, HEPN_DZIP3) and the central NACHT domain. The absence of such fusions in urochordates and vertebrates suggests lineage-specific loss of these *NLR* genes in both groups after their split from cephalochordates (Table S19, S20). In contrast, the domain combinations of BIR-NACHT, PYRIN-NACHT, and FISNA-NACHT only occur in vertebrates, suggesting that they are vertebrate-specific innovations (Table S19, S20).

### The proto-MHC genes

The vertebrate-specific MHC locus encodes a wide variety of cell surface molecules that mediate antigen recognition and initiate the adaptive immune response^90^. Its prototypical form, the proto-MHC, has an ancient metazoan origin^91^. Previous studies characterized some MHC-specific anchor genes in *B. floridae* based on fragmented genomic information and proposed a proto-MHC locus homologous to four human MHC paralogous blocks (1p31-1q31, 6p21.3, 9q32-34.3, and 19p13)^28,87^. Here we revisited the proto-MHC evolution by examining MHC-homologous genes in all five chromosome-level cephalochordate genome assemblies. A cephalochordate-human genome comparison revealed strong gene orthology correspondence between a single cephalochordate chromosome (e.g., the *A. lucayanum* chromosome 7) and four human chromosomes (chromosome 1, 6, 9, 19), with hits on human chromosome 6 corresponding to the human *MHC* locus (Figure S24–S28). In our *A. lucayanum*-human comparison, we found a total of 432 genes on *A. lucayanum* chromosome 7 displaying orthologous correspondence with genes on human chromosomes 1, 6, 9, and 19. Among them, 55 genes are orthologous to human *MHC* genes on human chromosome 6 (n=47) or to their close paralogs on human chromosomes 1 (n=11), 9 (n=11) and 19 (n=5) (Figure 6c). These 55 genes are scattered across the entire chromosome 7 in *A. lucayanum*, in sharp contrast to the much more concentrated distribution of their orthologs on human chromosomes: 1q23.3–q25.3 [*HSPA6*, *RNF2*], 6p21.33–6p21.31 [*GNL1*, *FKBP5*], 9q33.2–9q34.3 [*DAB2IP*, *EHMT1*], and 19p13.2–p13.11 [*SLC44A2*, *PBX4*]. Also, rather than being strongly co-localized within a narrow genomic locus like vertebrate MHCs, the cephalochordate orthologs of the 55 human *MHC* genes underwent substantial species-specific duplication and order reshuffling (e.g., *SKIC2*-like genes in *A. lucayanum* and *PSMB7/10*-like genes in *B. floridae*) (Figure S29). No orthologs of the human core *MHC* genes such as *HLA-A/B/C/DR/DP/DQ*, or local gene linkages corresponding to human *MHC* ClassI-III-II partitions could be detected in the cephalochordate chromosomes containing *proto-MHC* genes.

### Proto-RAGs

The extreme diversity of immunoglobulins and T cell–antigen receptors involved in adaptive immunity of jawed vertebrates is achieved by the RAG1 and RAG2 proteins via V(D)J somatic recombinations^92^. After the split of jawless and jawed vertebrates, RAG1 and RAG2 apparently evolved from an ancestral transposable element containing both sequences^93^. The recent discovery of intact proto-RAG transposable elements in *B. belcheri* has suggested cephalochordate *RAG1-like* (*RAG1L*) and *RAG2-like* (*RAG2L*) genes as the closest common ancestors for vertebrate *RAG1s* and *RAG2s*^31^; however, there is phylogenetic evidence indicates that hemichordate *RAG1L* is more closely related to vertebrate *RAG1* ^94,95^. To settle this debate, we made a genome-wide scan of all five cephalochordate species and found intact proto-RAGs (*RAG1L*+*RAG2L*) in *B. lanceolatum* (2 copies) as well as two additional *RAG1Ls* in *B. belcheri* and *B. japonicum* respectively (each with a single copy) plus one additional *RAG2L* in *B. lanceolatum* (Table S21). Note that we did not find intact proto-RAG in the *B. belcheri* genome assembly used in this study despite its better quality compared to the original short-read-based assembly where the *B. belcheri* proto-RAG was discovered^31^. This suggests that proto-RAG is still under dynamic evolution (therefore not fixed) even within *B. belcheri* populations. No *RAG1L* or *RAG2L* was found in *A. lucayanum* or *B. floridae*. We also extended our survey to several representative species of vertebrates, tunicates, hemichordates, and echinoderms. As expected, we found unambiguous copies of *RAG1* and *RAG2* in all surveyed jawed vertebrates (human, chicken, spotted gar and coelacanth, and elephant shark) but none in jawless vertebrates (lamprey and hagfish). In hemichordates and echinoderms, more *RAG1L* and *RAG2L* were found but with substantially varied species-specific distributions (e.g., 4 in the hemichordate *Ptychodera flava* but none in the hemichordate *Saccoglossus kowalevskii* for *RAG1L*). By combining all *RAG1(L)* and *RAG2(L)* genes identified in our survey and a recent study^95^, we re-evaluated their evolutionary history in the context of deuterostome evolution (Figure S30). Orthologs of both *RAG1(L)* and *RAG2(L)* in cephalochordates are tightly clustered together and clearly related to previously defined *RAGL-B* family members of hemichordates and echinoderms^94,95^. The vertebrate RAGs are most closely related to hemichordate and echinoderm *RAG1/2Ls* from the *RAGL-A* family as previously noted^94,95^. The *RAGL-A* family members from the hemichordate *P. flava* appear as the most closely related invertebrate ortholog to vertebrate *RAGs*. Taken together, our study with an expanded set of cephalochordate *RAGLs* provides additional support for a likely hemichordate origin of the *RAGL-A* gene through horizontal gene transfer in the ancient common ancestor of jawed vertebrates^95^.

## Discussion

In the present study, we used PacBio long-read sequencing and Hi-C technologies to construct a chromosome-level assembly of the genome of *Asymmetron lucayanum.* Compared to previously published *Branchiostoma* genomes, the *Asymmetron* genome is much larger and contains more inter-genic repeats and transposable elements. Nevertheless, both macrosynteny and microsynteny remain highly conserved between *Asymmetron* and *Branchiostoma*, enabling us to reconstruct the ancestral genomic architecture of the cephalochordate lineage, as well as to retrace its evolution during deuterostome diversification. Furthermore, by combining comparative genomics and developmental transcriptomic analyses, we show that purifying selection and constraints on co-transcriptional regulation may have contributed to the maintenance of the conserved microsynteny blocks among cephalochordate species. Overall, our findings collectively lead to a better understanding of the genome biology and evolution of chordates and help to illuminate the deep evolutionary history of many of the functionally important coding and regulatory loci of vertebrates.

Our results show that *Asymmetron* has a larger genome (chromosomally anchored size: 677 Mb) compared to *Branchiostoma* species (383 Mb to 490 Mb). This genome expansion appears to have resulted from the accumulation of intergenic repeats and TEs -- consistent with previous findings of genome expansions caused mainly by accumulation of intergenic repeats and TEs in animals with massively expanded genomes, such as the Mexican axolotl, Antarctic krill, and lungfishes^96–99^. However, each animal lineage appears to accumulate different kinds of repeat elements. For example, long terminal repeat (LTR) elements are the most abundant class of TEs in the axolotl, while LINEs are more prevalent in lungfishes. These two kinds of TEs belong to the same class of copy-and-paste retrotransposons, although they propagate via different mechanisms^100^. Interestingly, in the *Asymmetron* genome we observed a more substantial expansion of the TIR type of DNA transposons (Table S4), which belong to the “cut-and-paste” class of TEs. Such TIR-TE expansion is reflected by multiple families of TIR-TEs (e.g., CACTA, Mutator, PIF_Harbinger, etc.), underscoring their dominant roles in causing genome expansion. Overall, our observed substantial TE expansion in the *Asymmetron* genome provides another empirical example of how TE evolution can contribute to genome size variation among closely related species.

Despite the unique genome expansion by pervasive TE invasion, which presumably can trigger more chromosome rearrangements, the *Asymmetron* genome still retains strong macrosynteny and microsynteny conservation compared to those of *Branchiostoma* species. Moreover, sequence conservation is widespread across all five cephalochordates species in not only CDSs and UTRs but also in many intronic and intergenic regions. Aside from functional constraints imposed by selection, this observation also suggests a universally slow rate of genome evolution in cephalochordates. The substantial genome conservation between cephalochordates and vertebrates further reflects a deep constraint on genome architecture evolution in these two chordate subphyla. In contrast, genomes of tunicates (a.k.a., urochordates, representing the third chordate subphylum Urochordata) have rapidly evolving genomes^101,102^, which is why they were omitted from our analysis. One obvious question is what caused such accelerated genome evolution in tunicates? The answer may lie in the developmental mode. Cephalochordates and vertebrates have regulative development, which is thought to be ancestral in the Bilateria^103^, showing concomitant radial cleavage with cell fates determined gradually during embryogenesis. Thus, their cell fates depend to a considerable degree on cell-cell communication. The first two blastomeres, if separated, can each develop into a complete organism resulting in twins^104^. In vertebrates, the twins are typically identical, while in amphioxus, the twins are not quite identical as germ-cell determinants, which are concentrated into a small area of cytoplasm, are segregated into only one of the first two blastomeres^105^. Correspondingly, mutations in early-expressed developmental genes in cephalochordates and vertebrates are often lethal^106,107^. In contrast, tunicates evolved mosaic development with very early determination of cell fates^108^. If the first two blastomeres of an ascidian embryo are separated, each develops into what it would normally have formed in the intact embryo^109^. Thus gene knockouts done in very early ascidian embryos are often non-lethal^110^. Correlated with such early cell fate commitment, tunicate genomes have undergone loss of many genes and duplications of others. For example, while all cephalochordates have a single cluster of 15 *Hox* genes, different tunicates have 6–9 *Hox* genes^111^. In other words, the tunicates’ early decision of cell fates potentially lowered the evolutionary constraints of genes expressed later in development, resulting in a highly derived genome content and architecture.

The conserved macrosynteny between *Asymmetron* and *Branchiostoma* enabled us to reconstruct the ancestral Cephalochordate genomic architecture as 20 ALGs, similar to the chromosome composition of the extant *B. belcheri* genome^13^. From this ancestral condition, we infer that inter-chromosomal fusion events are the major factor shaping the current chromosomal structure of different cephalochordate species. Furthermore, our analysis shows that fusion events occurred in regions with segmental duplications, and are often coupled with higher abundance of repetitive sequences. Notably, a similar pattern was also observed for the ancestral chromosome 2 fusion (2q13–2q14.1) in the human genome^112,113^, suggesting a common mechanism in shaping the local genomic landscape of chromosome fusion sites. The identification of a large number of microsynteny blocks across cephalochordate species raises the question of the possible evolutionary constraints in maintaining such conservation of gene order. By combining comparative genomics and developmental transcriptomic analyses, we showed that purifying selection and constraints on co-transcriptional regulation likely contributed to these conserved patterns. This is in line with what has occurred in other metazoan lineages^114,115^, suggesting constraints on *cis*-regulatory elements as a powerful factor in shaping and maintaining local genome architectures.

Our comparative genomic analysis of chordate *Hox* genes suggests that the 15-gene configuration of the *Hox* cluster is ancestral for cephalochordates. Interestingly, the *Hox* clusters of both *A. lucayanum* and *B. belcheri* are inverted compared to other *Branchiostoma* species. These two species also have extensive expansion of TEs in the *Hox* flanking regions, as well as in particular genomic areas within the *Hox* clusters, suggesting a potential role for TEs in causing the inversions. From the inversion breakpoints and the types of TEs in these two species, it seems likely that the *Hox* inversions in *A. lucayanum* and *B. belcheri* are independent inversion events mediated by distinct TEs. Although the current Hi-C data are not sufficient to show clear local 3D genome interactions around the *Hox* cluster in *A. lucayanum*, Hi-C data from *B. belcheri* show evident overlap of the inversion break points just outside the boundaries of the two topologically associating domains (TADs) in the *Hox* cluster (see Figure 4f). In vertebrates (e.g., mouse), the TADs around the *Hox* gene clusters function in segregating regulatory elements and ensure proper gene expression patterns during embryonic development^116–118^. Thus, the observed TADs around the cephalochordate *Hox* cluster may also contribute to the regulation of the conserved temporal co-expression patterns of *Hox* genes in different species. This may impose strong evolutionary constrains on maintaining the “en-bloc” configuration of the *Hox* cluster in spite of TE invasion.

Interestingly, we also found intergenic invasion of different types of TEs in the *Hox* clusters of *A. lucayanum* (Mutator TE between *Hox9* and *Hox10*) and *B. belcheri* (Helitron TE between *Hox13* and *Hox14*). Invasion of TEs into *Hox* clusters has been associated with species radiation and morphological diversification in teleost fish and reptiles^119–121^. It is tempting to speculate that the observed TE invasion in *A. lucayanum* and *B. belcheri* may have caused some subtle changes in the expression pattern of nearby *Hox* genes, in turn leading to the morphological variation in cephalochordates, such as the myotome number variation among different *Branchiostoma* species and the distinct shapes of the caudal fin between *Asymmetron* and *Branchiostoma* genera^122^. While the association between *Hox* genes and major evolutionary transitions in body plans have been well documented^123–125^, studies on the genetic and functional variation of *Hox* genes among closely related species have been relatively limited. Fine-scale comparative studies coupled with genome editing perturbations may reveal novel insights regarding this issue.

Finally, by comprehensive genome-wide scans, we traced the evolutionary history of important immune genes (i.e., *TLR*, *NLR*, *MHC*, and *RAGs*). As cephalochordates fall in the key transition point between invertebrates and vertebrates, they have been important for understanding the evolution of chordate immune system before the appearance of adaptive immunity in vertebrates. Similar to previous findings in *Branchiostoma* species.^5,126^, we found lineage-specific expansion of innate immune receptor genes like *TLR* and *NLR* in *A. lucayanum*. Such expanded diversity in the innate immune system is important to cephalochordates as they live in shallow seas near the shore where the rich microorganisms are likely to constantly challenge their immune systems. While cephalochordates lack a functional adaptive immune system, prototypical forms of key adaptive immune genes such as *MHC* and *RAGs* are present in cephalochordate genomes, hinting at their deep evolutionary roots. Most interestingly, in contrast to the highly regionalized and compact *MHC* locus in vertebrates, cephalochordate *proto-MHC* genes are widely distributed across the *MHC*-corresponding chromosome, therefore preserving macrosynteny but not microsynteny. This is different from a previous suggestion based on a few anchor genes of a *Hox*-like “en bloc” configuration of proto-MHC in cephalochordates^28,87^. By this logic, the *MHC* regionalization and compaction should have occurred after the separation between cephalochordates and vertebrates. The recent description of a highly conserved and compact *MHC* locus in cartilaginous fish (including sharks, rays, and chimeras)^127^, sets a lower bound regarding the evolutionary time window for the *MHC* regionalization and compaction before the diversification of jawed vertebrates. The exact mutational and/or selection scheme shaped such a highly specialized genomic organization remains to be discovered.

In conclusion, our study assembled and characterized the genome of an early diverged cephalochordate species *A. lucayanum*, with which we carried out comparative genomic and transcriptomic studies with genomes of late diverging cephalochordates. This gave a comprehensive picture of the genome evolution of cephalochordates as a whole. By further extending our analysis to include more representative metazoan species (both invertebrates and vertebrates), we further illuminated the evolutionary history of important developmental (e.g., *Hox*) and immune genes (e.g., *TLR*, *NLR*, *MHC*, and *RAGs*) in the context of chordate evolution. Given the pivotal role of cephalochordates as an important model system for understanding early chordate evolution, we anticipate our work to serve as a foundational resource not only for future studies on the cephalochordates per se, but for a better understanding of early chordate evolution and the origin of vertebrates in general.

## Methods and Materials

### Sample collection and sequencing

Specimens of *A. lucayanum* were collected in Bimini, Bahamas as previously described^37^. After transportation to the lab, they were fed twice a day on phytoplankton (*Isochrysis sp.*, *Tisochrysis 463*, *Pavlova*) and placed on an artificial moonlight regime that mimics that in Bimini together with a 14-hr light/10-hr dark daylight regime. As in the field, the animals spawned a few days before each new moon. A single ripe male was isolated in a 6 cm petri dish containing about 10 ml of sea water. The male spawned about 30 min after the artificial sunset and the sperm were pelleted by centrifugation.

### DNA extraction

High molecular weight DNA was extracted from the sperm pellet with the Qiagen DNeasy kit (Cat: 69504, Qiagen, Germantown, MD, USA). Proteinase K digestion at 56°C was extended to 24 hrs. The column was eluted with three 100-µl aliquots of pH 9.0 water. DNA was stored at 4°C with DNAStable plus (Cat: 53091-016, Biomatrica, San Diego, CA, USA).

### PacBio long-read sequencing

Using the g-tube (Covaris, Woburn, MA, USA), we generated 20-kb fragments by shearing genomic DNA according to the manufacturer’s recommended protocol. The AMpureXP bead purification system (Beckman Coulter) was used to remove small fragments. A total of 5 μg for each sample was used for library preparation. The SMRTbell library was constructed with the SMRTbell Template Prep Kit 1.0 (Cat: PN100-259-100). The BluePippin Size selection system (Saga Science, Beverly, MA, USA) was used to remove the small DNA fragments from the sequencing library. After annealing a sequencing primer to the SMRTbell template, DNA polymerase was bound to the complex (Sequel Binding Kit 2.0). Purification was performed with SMRTbell Clean Up Columns v2 Kit-Mag (Cat: PN01-303-600). The purification step comes after polymerase binding to remove excess unbound polymerases and polymerase molecules which are bound to small DNA inserts. The HMW MagBead Kit (Cat: D6060, Zymo Research, Irvine, CA, USA) was used to bind the library complex to MagBeads before sequencing. MagBead bound complexes provide for more reads per SMRT Cell. This polymerase-SMRTbell-adaptor complex is then loaded into zero-mode waveguides (ZMWs). The SMRTbell library was sequenced using SMRT cells (Pacific Biosciences, Sequel™ SMRT® Cell 1M v2) and the Sequel Sequencing Kit 2.0. Light-pulse movies (1L×L600 min) were captured for each SMRT cell using the Sequel (Pacific Biosciences, USA) sequencing platform with the P6-C4 chemistry.

### Initial diploid genome assembly and polishing

De novo genome assembly was performed under the framework of LRSDAY^128^ (v1.5.0). Given the high allelic heterozygosity of cephalochordates, we adopted two assembler protocols that were tailored for assembling such genomes. First, we used Canu^129,130^ (v1.8) with the customized settings as suggested by its developers for highly heterozygous genome assemblies: correctedErrorRate=0.105 corOutCoverage=200 batOptions=-dg 3 -db 3 -dr 1 -ca 500 -cp 50. In addition, we also used Falcon-Unzip^131^ (shipped with LRSDAY via pb-assembly [v0.0.2]) with the following configuration options: pa_DBsplit_option= -x500 -s200 ovlp_DBsplit_option= -x500 -s200 pa_REPmask_code= 0,300;0,300;0,300 pa_HPCdaligner_option= -v -B128 -M24 pa_daligner_option= -k18 -e.75 -l1200 -h256 -w8-s100 falcon_sense_option= --output-multi --min-idt 0.70 --min-cov 3 --max-n-read 200 ovlp_daligner_option= -k24 -e.96 -l1800 -h600 -w6 -s100 ovlp_HPCdaligner_option= -v-B128 -M24 overlap_filtering_setting= --max-diff 100 --max-cov 100 --min-cov 2 length_cutoff_pr=1000. For both assemblies, our previously estimated *A. lucayanum* genome size (644 Mb)^38^ was used to set the genome size prior. Although FALCON-Unzip has the design feature of doing haplotype phasing, it didn’t perform well for our *A. lucayanum* assembly, with a small fraction of an alternative assembly separated from the primary assembly. We therefore re-merged its separated primary and alternative assembly files and treated them as an unphased diploid assembly. The resulting assembly size from both assemblers tallied around 1.3 Gb, therefore indicating a diploid genome assembly. For both assemblies, we used PacBio’s pbalign-quiver/arrow pipeline (shipped with LRSDAY via pb-assembly v0.0.2) for three consecutive rounds of PacBio-read-based polishing. BUSCO^132^ (v5.3.2; with the metazoa_odb10 database) was used to evaluate the completeness of both Canu and FALCON-Unzip diploid assemblies.

### Diploid assembly decoupling and haplotype assembly separation

Haplomerger^133,134^ (v20180603) was used to decouple the Canu and FALCON-Unzip diploid assemblies. The resulting haplotype-phased Canu assembly consists of a 691.6-Mb primary assembly, a 635.3-Mb alternative assembly, and 29.7-Mb patch sequences. The resulting haplotype-phased FALCON-Unzip assembly consists of a 681.4-Mb primary assembly, a 632.1-Mb alternative assembly, and 44.0-Mb patch sequences. The patch sequences are sequences from the input diploid assembly that cannot be decoupled. We evaluated the completeness of these two phased assemblies with BUSCO and selected the haplotype-phased Canu assembly for further analysis given its better continuity and BUSCO completeness. Noticing that the patch sequences still matched with ∼20-40 BUSCO genes, we added back the patch sequences to the associated primary and alternative assemblies to ensure maximal completeness.

### RNA extraction and transcriptome sequencing

For Iso-Seq, total RNA of *A. lucayanum* was isolated from several developmental stages (blastula, G6, G7, N1, N2, L1, and adult). These samples were pooled together for the PacBio SMRTbell library preparation. Sequencing was carried out on a PacBio Sequel system with two SMRT Cells. For bulk RNA-seq, total RNA of *A. lucayanum* from the blastula, 1-8 cell, G3-G6, and N2 stages was isolated separately for library preparation using the TruSeq kit (Illumina, USA) separately. The sequencing was carried out using the Illumina HiSeq2500 system at the NGS Genomics core facility of the Biodiversity Research Center affiliated with the Academia Sinica. Likewise, multi-stage total RNA samples were collected for *B. belcheri* (stage: 1-cell, B, G6, N4, L0) and *B. japonicum* (stage: 1-cell, B, G6, N2, T1) respectively for RNA-seq experiment following the same library preparation and sequencing protocols. The developmental stage notation used here was based on an updated developmental staging system recently proposed^135^.

### Quantification of expression profiles for different cephalochordate species

The gene expression profiles of different cephalochordate species were analyzed using both the bulk-RNA-seq datasets generated by this study (*A. lucayanum*: 3 developmental stages; *B. belcheri*: 5 developmental stages; *B. japonicum*: 5 developmental stages) as well as published studies (*B. floridae*: 12 developmental stages; *B. lanceolatum*: 16 developmental stages and 9 tissue types)^10,136^ (Table S1; Table S5). For each RNA-seq dataset, library preparation adaptors and low-quality bases from the raw reads were removed by Trimmomatic^137^ (v0.36). The trimmed reads were then mapped to the corresponding genome assemblies using STAR^138^ (v2.7.10a) with default parameters. Raw count tables for each sample were generated using featureCounts^139^ (v2.0.1) based on the gene annotations generated by this study. Expression levels were quantified as transcripts per million (TPM) and visualized as log_2_(TPM+1) using ComplexHeatmap^140^ (v2.22.0). Since different developing staging systems were used across these datasets, we remapped their respective staging system to the unified staging system^135^ (Supplementary Dataset 2).

### 3D-DNA-based chromosome-level scaffolding

We used the HMW DNA prepared as described above but from a different *A. lucayanum* individual for the CHiCAGO and Hi-C library construction (with the DpnII restriction enzyme) by Dovetail Genomics (Scotts Valley, CA, USA). Both libraries were initially quality-checked by the MiSeq platform (Illumina, CA, USA) with ∼1–2 million paired-end (PE) 75-bp Illumina reads and then formally sequenced by the HiSeqX platform (Illumina, CA, USA) with PE 150-bp reads. The haplotype-decoupled Canu primary and alternative assemblies (both with patch sequences added back) were scaffolded through the Dovetails HiRise software pipeline^42^. The resulting chromosome-level assemblies were manually evaluated and curated using the Juicebox^141^ (v1.9.8).

### *K*-mer-based genome size estimation

To derive the genome-wide *k*-mer distribution of the five cephalochordate species, we used Jellyfish^142^ (v2.2.10; option: count -C -m 27 -s 1000000000) to examine their *k*-mer characteristics with their whole-genome shotgun reads (Table S5). The outputs were further processed by GenomeScope^143^ (v2.0) for genome size estimation.

### Repeat element and gene annotation

For both the *Asymmetron* genome assembly as well as the published *Branchiostoma* assemblies (Table S3, S8), we employed EDTA^144^ (v2.0.1; options: --species others --step all --sensitive 1--anno 1) to construct their respective repeat libraries, which were further fed to RepeatMasker (v4.1.1; https://www.repeatmasker.org) for repeat-aware soft masking. Based on the masked genome assemblies, protein-coding gene and tRNA gene annotations were carried out using FunAnnotate (v1.8.15; https://github.com/nextgenusfs/funannotate; model training options:

--repeats2evm --busco_db metazoa --keep_no_stops --optimize_augustus --organism other

--max_intronlen 200000) with species-specific transcriptome evidence supports. For *A. lucayanum*, the Iso-Seq (mixed developmental stages) and bulk RNA-seq (from three developmental stages) data collected for this study were used (Table S1). For the four *Branchiostoma* species, public Iso-Seq and bulk RNA-seq data were used (Table S5). The completeness of the final protein-coding gene annotation was evaluated by BUSCO^132^ (v5.3.2) based on the metazoa_odb10 database. RNAmmer^145^ (v1.2) was used for rRNA gene annotation.

### Phylogenetic analysis and molecular dating for the species tree

We used OrthoFinder^146^ (v2.4.0) to identify orthologous groups shared by the five cephalochordates as well as three other invertebrates (acorn worm, sea urchin, scallop) and seven vertebrates (human, mouse, chicken, frog, spotted gar, hagfish) (Table S8). A total of 947 strictly conserved one-to-one ortholog groups was selected. The protein sequences of each gene were aligned by MUSCLE^147^ (v3.8) and their concatenated super matrix was constructed accordingly. After alignment trimming by ClipKIT^148^ (v1.4.1; option: -m smart-gap), IQTREE^149^ (v1.6.12) was employed for tree construction with the options: -safe -m MFP -nstop 150 -bb 1000 -bnni -alrt 1000. Based on the same protein alignment matrix and phylogenetic tree, MCMCTree (shipped with PAML v4.10.0) was further used for molecular dating analysis. PAML’s CODEML program was first executed with the pre-shipped LG substitution matrix to estimate the branch lengths, gradient, and Hessian, all of which are needed for MCMCTree. Fossil-based vertebrate-specific divergence time points of a few internal nodes were used for time calibration (Table S9) and an upper bound of 830 MYA was used to cap the divergence time between human and scallop. The MCMCTree run was executed with the following options: seqtype=2, clock=2, model=3, alpha=0.5, ncatG=5, cleandata=0, BDparas=1 1 0.1, rgene_gamma=2 20, sigma2_gamma=1 10, burnin=2000000, sampfreq=1000, nsample=20000. FigTree (v1.4.4; http://tree.bio.ed.ac.uk/software/figtree/) and iTOL^150^ (v6) were used for phylogenetic tree visualization.

### Macrosynteny and microsynteny

For macrosynteny, rbhxpress (v1.2.3; https://github.com/SamiLhll/rbhXpress) was used to identify orthologous gene pairs shared between the two compared species. MacrosyntR^151^ (v0.3.3) was then used to construct Oxford grid dotplots for comparing the chromosomal positions of these orthologous genes. Moreover, by assuming a null hypothesis that orthologous genes are randomly distributed across both genomes, macrosyntR also performed a significance test on macrosynteny conservation, based on which homologous chromosome pairs can be deduced. Additionally, for each pairwise comparison, a macrosynteny conservation index was defined by dividing the number of orthologous genes located in all identified homologous chromosomes by the total number of orthologous gene pairs. Finally, we used the odp pipeline^152^ (v0.3.0) to further evaluate the correspondence between cephalochordate genomes and the previously defined Bilaterian-Cnidarian-Sponge (BCnS) ancestral linkage groups. GENESPACE^153^ (v0.9.4) and chromoMap^154^ (v4.1.1) were used for macrosynteny visualization.

For microsynteny, we used MCscanX^155^ (Github commit version: a8443a9; options: -s 5 –e 1e-05 –m 25) to identify gene clusters with conserved orders shared between species. The size distribution of microsynteny blocks shared across the five cephalochordate species was summarized based on the number of one-to-one orthologous genes encompassed by the corresponding block. A minimal requirement of three such genes within a defined microsynteny block was used. The observed data distribution was further fitted by different statistical distributions such as log normal, gamma, Weibull, and exponential by the *R* package fdistplus^156^ (v1.1). In addition to macrosynteny, we also defined a microsynteny conservation index by the proportion of orthologous genes being colinearly organized.

### Molecular evolution analysis for genes within microsynteny blocks

For each one-to-one orthologous gene enclosed in our identified cephalochordate microsynteny blocks, we used KaKs_Calculator^157^ (v2.0; option: -m MA) to calculate different molecular evolution metrics (dN, dS, and dN/dS). The values correspond to all pairwise comparison combinations across the five cephalochordate species. For each molecular evolution metric, its pairwise Spearman correlation coefficients between different one-to-one orthologous genes within the same microsynteny block were further calculated with additional Fisher’s *z*-transformation (for converting the skewed distribution of Spearman correlations into approximate normal distribution as previously suggested^115^). In parallel, we also reshuffled these within-block genes by randomizing their gene-block correspondence and applied the same in-block dN, dS, and dN/dS correlation calculation to the permutated datasets. A total of 1000 reshuffled datasets were generated and analyzed in this way. A two-sided Wilcoxon rank-sum test was used for statistical comparison between observed and reshuffled datasets.

### Expression correlation analysis for genes within microsynteny blocks

For genes enclosed in the same cephalochordate microsynteny blocks, we calculated the pairwise Spearman correlation coefficients of their gene expression TPM values across multiple developmental stages with additional Fisher’s *z*-transformation as previously suggested^115^. In parallel, expression correlation coefficients were also calculated for the 1000 reshuffled datasets described above (See: molecular evolution analysis for genes within microsynteny blocks). A two-sided Wilcoxon rank-sum test was used for comparing the expression correlation coefficients between observed and reshuffled datasets.

### *Hox* cluster analysis

The *Hox* clusters of the five cephalochordate species were identified based on the flanking anchor genes such as *MEOX2*, *MTX2*, and *SMUG1*. For each individual *Hox* gene (e.g., *Hox1*, *Hox*2, etc.), we used MACSE^158^ (v2.04) to construct their CDS and protein alignments with the following options: -prog alignSequences -gc_def 1 and -prog exportAlignment -align Hox.cds.macse_NT.aln.fa -codonForFinalStop --- -codonForInternalStop NNN -codonForInternalFS NNN -codonForExternalFS --- -charForRemainingFS -. KaKs_Calculator^157^ (v2.0; option: -m MA) was then used to calculate dN, dS, and dN/dS values. In addition, we retrieved the *Hox* genes from human and chimpanzee from Genbank and constructed their CDS alignment and performed dN, dS, and dN/dS calculations in the same way. The phylogenetic tree of the *Hox* genes from 15 representative species was constructed with IQTREE^149^ (v1.6.12) from protein sequence alignments and further visualized with iTOL^150^ (v6). The genome synteny plot for the *Hox* cluster comparison among *A. lucayanum*, *B. floridae*, and *B. belcheri* was plotted by TBtools^159^ (v2.210).

### Estimation of mutation rates in cephalochordates

For each of the 947 strictly conserved one-to-one orthologs described above, we calculated its pairwise dS values among the five cephalochordate species using KaKs_Calculator^157^ (v2.0; option: -m MA). Based on this, a corresponding median dS value was calculated for each cephalochordate species pair. The median dS values were linearly correlated with the estimated divergence time in the corresponding species pairs. Based on the correlation equation of dS = µ · 2T, we estimated the cephalochordate-specific mutation rate (μ), which was further used for the estimation of LTR-TE insertion time.

### LTR-TE insertion time estimation

For annotation of LTR-TEs in the five cephalochordate species, we first estimated their insertion times by directly comparing the sequence divergence (with Jukes-Cantor correction^160^) of the two flanking LTR sequences of each intact LTR-TE and calculating their insertion times with our estimated cephalochordate-specific mutation rate (μ=4.65×10^-9^ substitutions/base/year). To further control possible underestimation introduced by gene conversion^161^, for intact LTR-TEs with unambiguous strand information (annotated by EDTA^144^), we further examined their potential gene conversion tracts (criteria: *P*-value <0.05, length >10 bp) using GENECONV^162^ (v1.81; options: /w123 /lp /f /eb /g1 –nolog) in all pairwise combinations following a previously proposed procedure. The detected gene conversion tracts were subsequently excluded in the estimation of sequence divergence and insertion time of LTR-TE.

### Multi-species whole-genome alignment and identification of constrained regions

We built a five-way cephalochordate whole-genome alignment (including all five amphioxus species examined in this study) and a six-way vertebrate whole genome alignment (including spotted gar, frog, chicken, platypus, mouse, and human) respectively with Cactus^163^ (v2.7.0). For each alignment, the cactus-hal2maf tool (shipped with Cactus) was used to convert Cactus’ output files (in HAL format) into the MAF format using the respective reference coordinate system (*A. lucayanum* for the cephalochordate alignment and human for the vertebrate alignment) with the options: --noAncestors --onlyOrthologs --maxRefGap 0. Based on these two multi-way whole-genome alignments, we employed phyloP (options: -msa-format MAF--wig-scores --method LRT --mode CONACC) from the PHAST^164^ package (v1.5) to score the sequence conservation levels. Conserved bases are defined as those with phyloP scores >0 and covered by all aligned species. Afterwards, an additional set of chordate conserved bases was further constructed by intersecting the cephalochordate and vertebrate conserved base sets based on coordinate correspondence inferred from the Alu-human whole genome alignment generated by LastZ^165^ (v1.04.22).

Cephalochordate conserved bases were merged into cephalochordate ultra-conserved regions (UCRs) shared between cephalochordates and vertebrates with the following criteria: gap size <5 bp, block size ≥20 bp, fraction of conserved bases ≥60%. The genomic coordinates of our identified cephalochordate UCRs were compared with those of ATAC-seq-based *B. lanceolatum* open chromatin peaks^10^ and known *B. belcheri* and *B. floridae* micro-RNA genes (curated in miRbase database^59^) under the same reference system using BEDtools^166^. The *B. lanceolatum* open chromatin peaks were called based on the genomic coordinate of an older version of the *B. lanceolatum* genome assembly (NCBI Genbank accession: GCA_900088365.1). We used several tools from the UCSC Utilities^167,168^ (namely faSplit, blat, liftUp, axtChain, chainMergeSort, faSize, chainNet, netChainSubset, LiftOver) to create the correspondence chain file between this old *B. lanceolatum* assembly and the one that we used (Table S8) and to perform the coordinate conversion accordingly. As for *B. belcheri* and *B. floridae* micro-RNA genes, we retrieved their curated sequences from miRBase database^59^ and used BLAST^169^ to locate their corresponding locations on the *B. belcheri* and *B. floridae* genome assemblies that we used (Table S8).

We further extracted the noncoding fraction of the cephalochordate UCRs (which tallies 805,481 regions) and used it as the query dataset. Meanwhile, we randomly sampled the noncoding regions of the *A. lucayanum* reference genome for 100 simulated datasets with the sampled region number and region size distribution matched with the query dataset. The query and simulated datasets were collectively used for a test of motif enrichment based on the JASPAR2024 CORE nonredundant database^80^. For each curated JASPAR motif, we employed the AME program^170^ (e-value cutoff: 0.05) from the MEME suite^171^ (v5.5.7) to run the analysis with our query dataset against each of the 100 simulated dataset. The corresponding *P*-value for each JASPAR motif was calculated as the number of nonsignificant tests divided by 100.

We also merged chordate conserved bases into chordate UCRs with the same merging criteria used for merging cephalochordate UCRs. Given the much larger evolutionary time-scale for chordate UCRs, additional checks were performed to ensure that these chordate UCRs are: (1) supported by the pairwise LastZ alignment of *A. lucayanum* against each of the six vertebrates; (2) within or adjacent to the same homologous genes that have maintained the same physical relationship to the corresponding gene annotations (e.g., both being intronic) in *Asymmetron* and human reference genomes. Based on the human gene associated with these chordate UCRs, the following functional enrichment analyses were examined: 1) Gene Ontology (GO) term, 2) human variants implicated in diseases (ClinVar^82^ v20250330), 3) a merged gene set (n=857) with high-priority clinical significance^172,173^. The GO enrichment test was assessed by the *R* package ClusterProfiler^174^ (v4.8.3). For the enrichment test for ClinVar variants, the ClinVar variants were retrieved from (https://ftp.ncbi.nlm.nih.gov/pub/clinvar/vcf_GRCh38/clinvar_20250330.vcf.gz) with overlapping variant regions merged by BEDTools^166^ (v2.29.2) and variants located on sex chromosomes were excluded. The permutation-based test (1000 permutations) was applied separately by regioneR^175^ (v1.22.0)’s permTest function (evaluate.function=meanDistance) for coding and noncoding chordate UCRs and combined with randomly sampled coding and noncoding regions (with matched region size) from the human reference genome (GRCh38). The enrichment test against the clinically relevant gene set was assessed by Fisher’s exact test based on gene set overlaps by the *R* function fisher.test().

### Genome-wide survey for immune-related genes and gene families

For *TLR*, *NLR*, and *RAG* genes, a comprehensive genome-wide survey was performed for a total of 32 species representing different metazoan lineages, including 5 cephalochordates, 10 vertebrates, 2 urochordates, 2 hemichordates, 3 echinoderms, 3 insects, 1 branchiopod, 1 annelid, 1 nemertean, 1 mollusk, 1 bryozoan, and 2 cnidarians. Proteomes of these representative species were retrieved (Table S8) and used for genome-wide scans performed by the Pfamscan script (v1.6) from the HMMER^176^ package (v3.3.2; option: -e_seq 0.01). When one gene corresponded to multiple transcripts (and therefore multiple protein sequences), only the longest one was considered to avoid double counting. Based on the Pfamscan results, two e-value cutoffs (0.01 and 0.0001) were used to identify genes that encode key characteristic protein domains (e.g., TIR, LRR, NACHT, RAG1, RAG2, etc.) related to the *TLR*, *NLR*, and *RAG* genes. While the numbers of identified genes under both e-value cutoffs were reported in the supplementary tables (Table S19, S20), those corresponding to the 0.01 cutoff were reported in the main text. For *NLRs*, there is a great diversity of other N-terminal and C-terminal domains co-existing with their central NACHT domains. Therefore, we extensively examined the diversity of domain combinations in not only the 32 above-mentioned species but also all NACHT-encoding protein sequences retrieved by the Taxonomy-based query function of the InterPro database^177^. The metazoan lineages surveyed include: Porifera, Placozoa, Anthozoa, Hydrozoa, Ecdysozoa, Spiralia, Echinozoa, Asterozoa, Tunicata, Cyclostomata, Gnathostomata (Table S20). The gene and domain combination statistics for Cephalochordata were calculated by combining the corresponding results of the five amphioxus species examined in this study.

### Gene tree analysis of the *Hox*, *TLR*, *NLR*, and *RAG* genes

For the *Hox* gene cluster, we retrieved the *Hox* gene members of all compared species based on BLAST-based homology searches and literature surveys. We retrieved the *TLR*, *NLR*, and *RAG* genes based on their encoded domains as described above. For each gene or gene family, we used MUSCLE^147^ (v3.8) to construct their full-length protein sequence alignment, based on which IQTREE^149^ (v1.6.12) was used for gene tree construction with the options: -safe -m MFP-nstop 150 -bb 1000 -bnni -alrt 1000. Regarding tree visualization, iTOL^150^ (v6) was used for the *TLR* and *NLR* tree, while ggtree^178^ (v2.4.0) was used for visualizing the *RAG1* and *RAG2* trees.

### Comparative analysis for proto-MHC genes

A list of the *MHC* and *proto-MHC* anchor genes was collected from the literature^28,29,127,179^ and correspondence of orthologs in both human and the five cephalochordate species was identified using OrthoFinder (v2.4.0). The chromosomal organization of these MHC and proto-MHC genes along the human and cephalochordate chromosomes was visualized using the *R* package geneviewer^180^ (v0.1.8; https://github.com/nvelden/geneviewer). The genome synteny correspondence between the *A. lucayanum* chromosome 7 and the four human chromosomes carrying *MHC*-related duplication blocks (i.e. human chromosomes 6, 1, 9, and 19) was visualized by TBtools^159^ (v2.210).

## Data availability

The raw reads of genome, 3D-genome, and transcriptome sequencing for *A. lucayanum* have been deposited in the Sequence Read Archive (SRA) database (https://www.ncbi.nlm.nih.gov/sra) of the National Center for Biotechnology Information (NCBI) under the BioProject accession number of PRJNA1130632 and sequencing run accession numbers of SRR29874002, SRR30049563, SRR29874001, SRR30169925, SRR30153493, SRR30153494, and SRR30153495. The *A. lucayanum* reference and alternative genome assemblies generated by this study have been deposited in Genome Warehouse of National Genomics Data Center (https://ngdc.cncb.ac.cn/gwh) under the accession numbers of GWHFWAS00000000.1 and GWHFWAV00000000.1 respectively. The gene annotation files for *A. lucayanum*, *B. belcheri*, *B. floridae*, *B. japonicum*, and *B. lanceolatum* generated by this study have been deposited in Zenodo (https://doi.org/10.5281/zenodo.15280774). The scripts used for various analyses performed in this project were deposited at GitHub (https://github.com/ArthurSilver/Asymmetron_Pipelines).

## Acknowledgements

We thank Dr. Nicholas D. Holland and the staff of the Bimini Biological Station, Bimini, Bahamas, for facilitating the collection of specimens of *Asymmetron lucayanum.* We also thank the staff at the core facility and the Marine Research Station of the Institute of Cellular and Organismic Biology, and NGS Genomics core facility of the Biodiversity Research Center, Academia Sinica for technical assistance.

## Author contributions

JXY, JKY, LZH, SJC conceived the study. YR, ZM, CYL, LY, HL, JXY performed the data analysis and results visualization. LZH provided the animal sample for genome and transcriptome sequencing. SJC fund the long-read-based genome sequencing and JKY fund the Iso-Seq and Dovetails’ CHiCAGO and HiC library construction and sequencing. All authors contributed to results interpretation and discussion. JXY, JKY, and LZH wrote the initial draft of the manuscript, and all authors reviewed and contributed to the final version.

