## Supplementary Figures for "The genome of the early diverged amphioxus, *Asymmetron lucayanum*, illuminates the evolution of genome architecture and gene repertoires in cephalochordates"

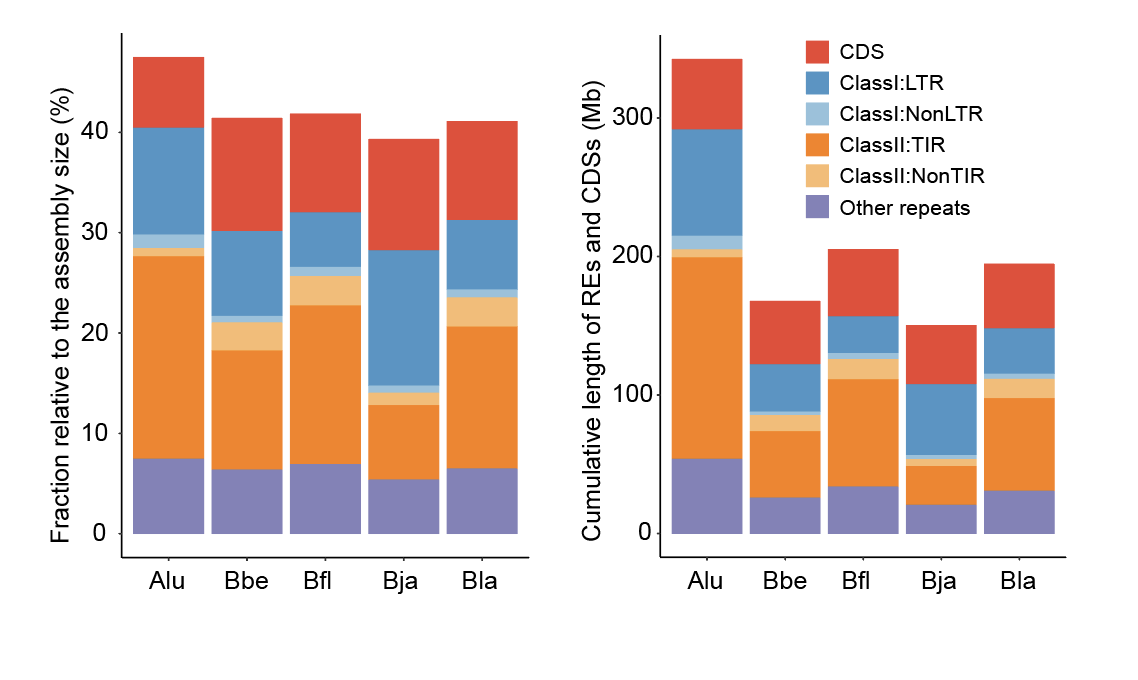

**Figure S1. Comparison of TE content across the five amphioxus genomes.**

Alu: *Asymmetron lucayanum*; Bbe: *Branchiostoma belcheri*; Bfl: *Branchiostoma floridae*; Bja: *Branchiostoma japonicum*; Bla: *Branchiostoma lanceolatum*.

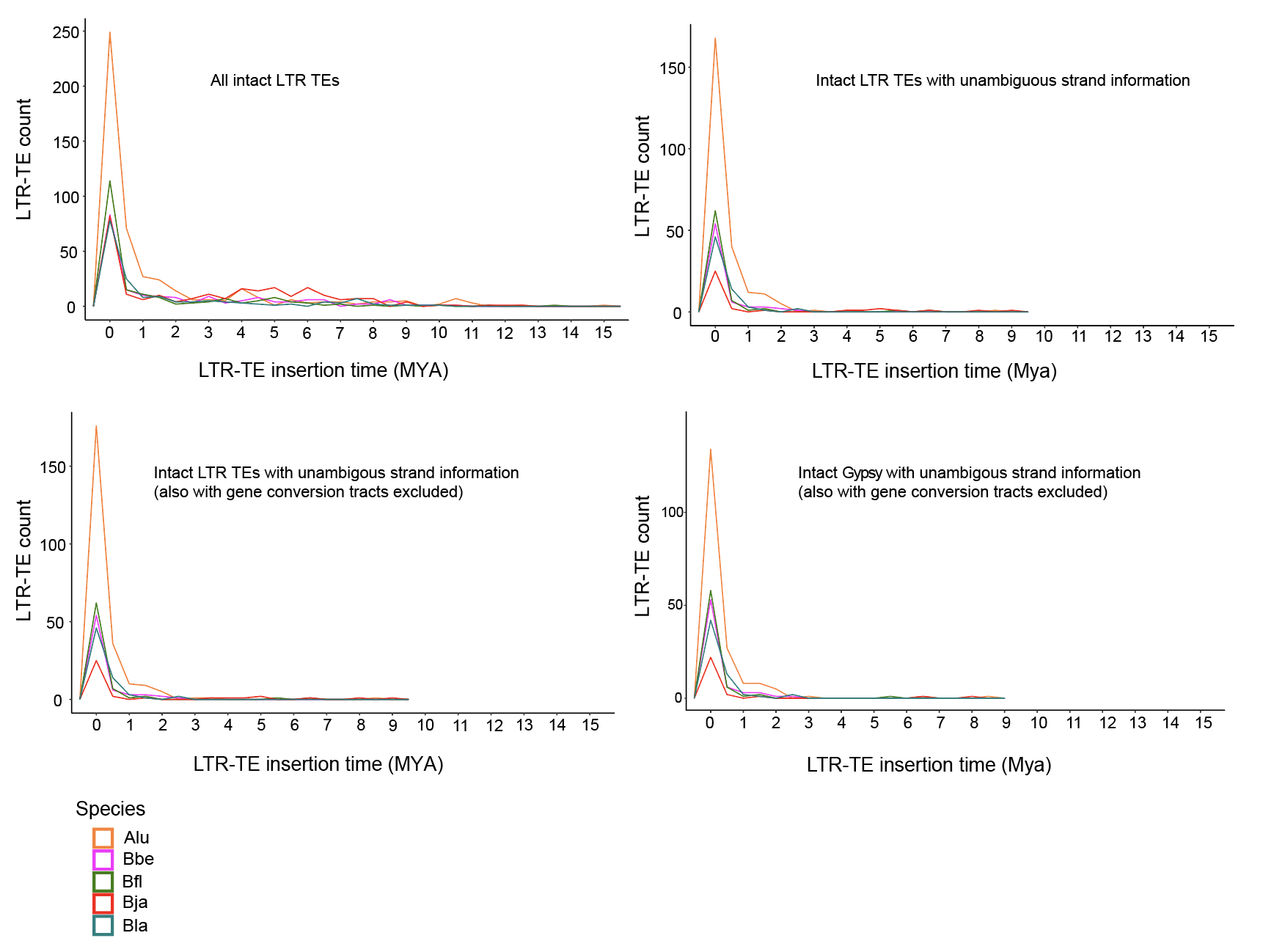

**Figure S2. Molecular dating of the LTR transposable elements (LTR-TEs) insertion time.**

The LTR-TE insertion time was estimated with the Juke-Cantor corrected sequence divergence between the 5’- and 3’-flanking LTR sequences of the same TE. Alu: *Asymmetron lucayanum*; Bbe: *Branchiostoma belcheri*; Bfl: *Branchiostoma floridae*; Bja: *Branchiostoma japonicum*; Bla: *Branchiostoma lanceolatum*. The LTR-TEs identified in this study predominantly either Gypsy TEs or unclassified LTR-TEs.

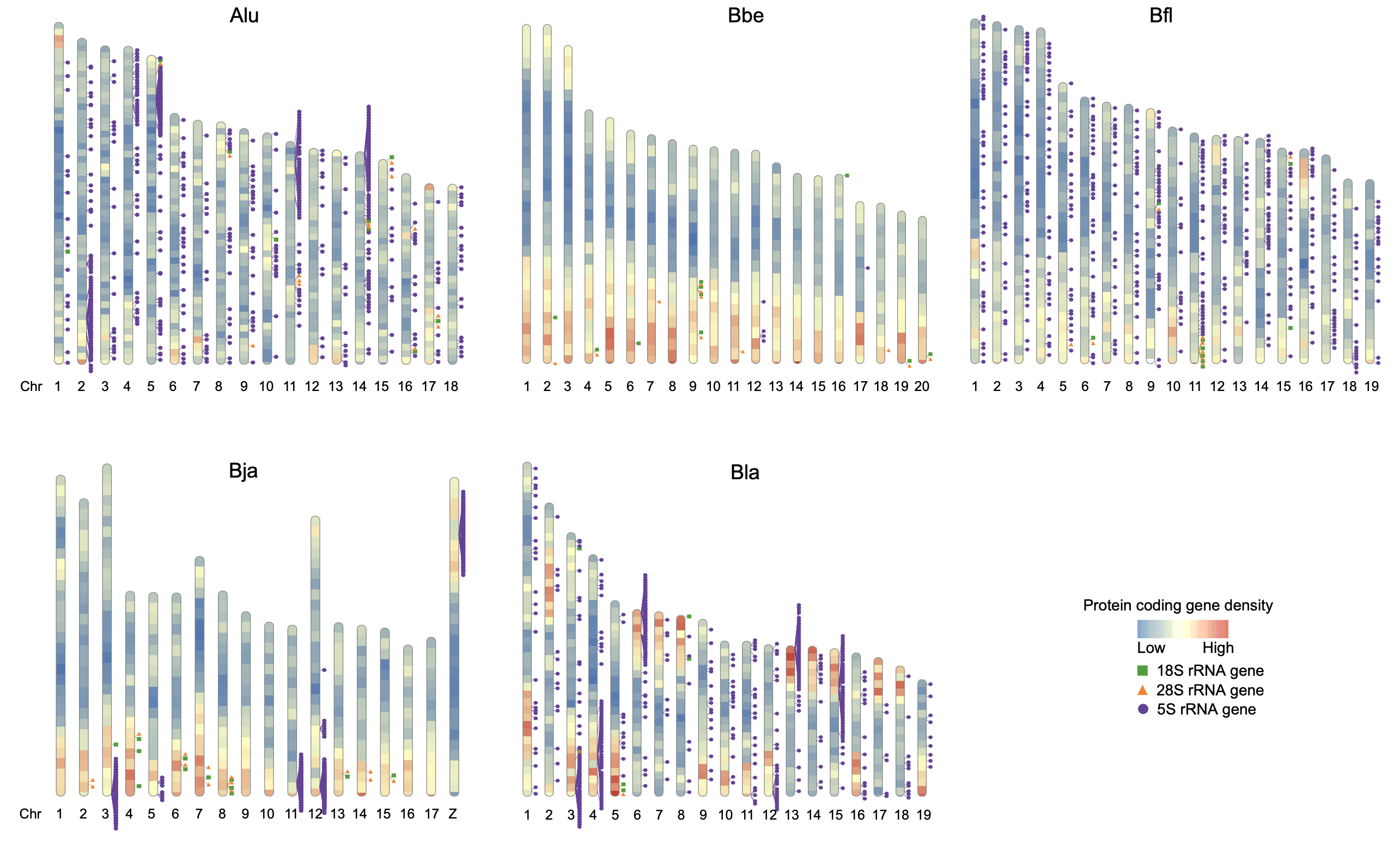

**Figure S3. Genomic distribution of rRNA genes in five cephalochordate species.**

Alu: *Asymmetron lucayanum*; Bbe: *Branchiostoma belcheri*; Bfl: *Branchiostoma floridae*; Bja: *Branchiostoma japonicum*; Bla: *Branchiostoma lanceolatum*.

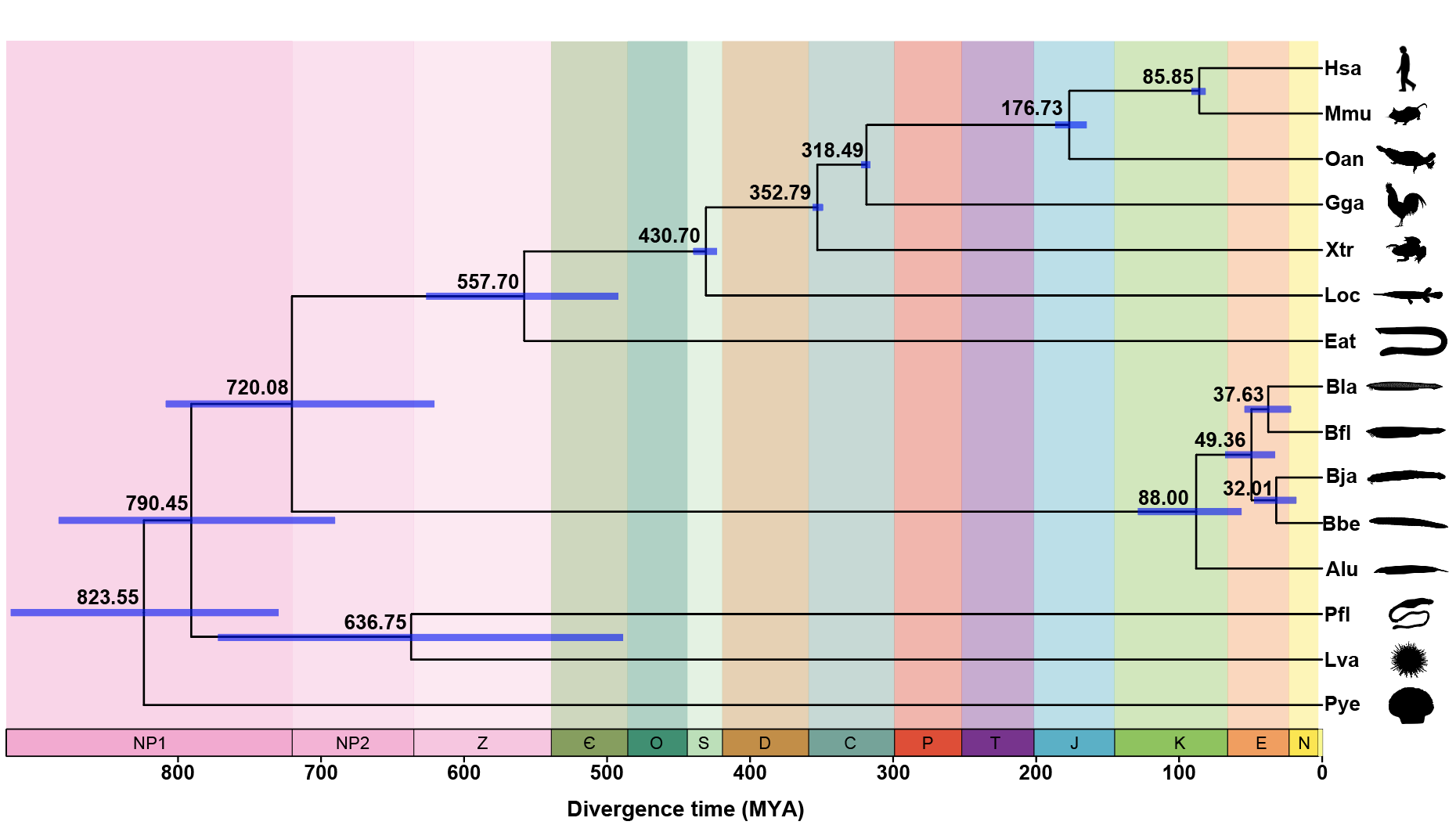

**Figure S4. Fossil-calibrated evolutionary time frame for cephalochordate and chordate evolution.**

Alu: *Asymmetron lucayanum*; Bbe: *Branchiostoma belcheri*; Bfl: *Branchiostoma floridae*; Bja: *Branchiostoma japonicum*; Bla: *Branchiostoma lanceolatum*; Eat: *Eptatretus atami*; Gga: *Gallus gallus*; Hsa: *Homo sapiens*; Loc: *Lepisosteus oculatus*; Lva: *Lytechinus variegatus*; Mmu: *Mus musculus*; Oan: *Ornithorhynchus anatinus*; Pfl: *Ptychodera flava*; Pye: *Patinopecten yessoensis*; Xtr: *Xenopus tropicalis*.

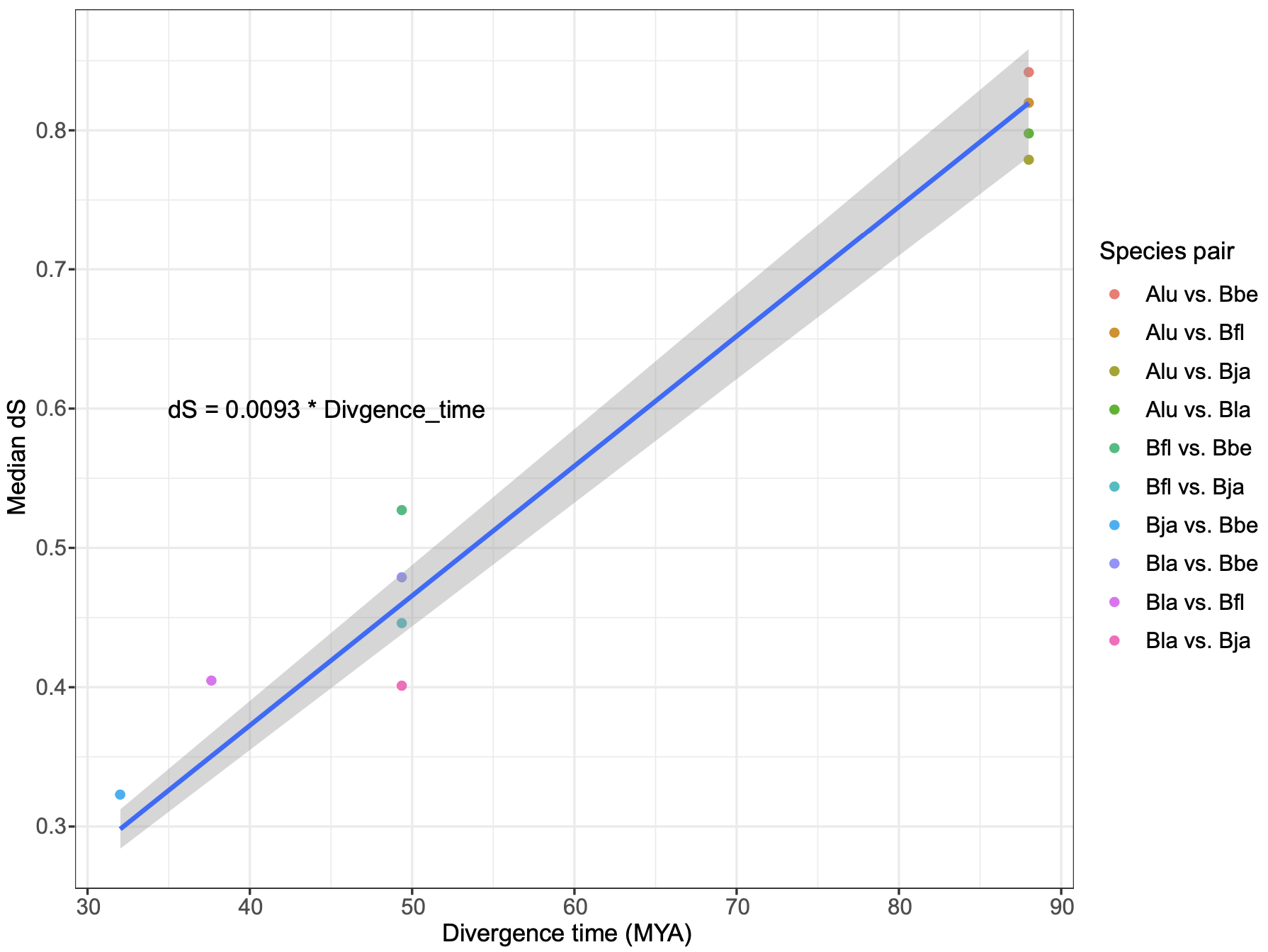

**Figure S5. The linear correlation between genome-wide median synonymous substitution rate (dS) and divergence time (measured in MYA) based on the pairwise comparison among different cephalochordates.**

The median dS calculation was based on 947 1-to-1 ortholog genes used for the phylogenomic analysis of this study. MYA: million years ago. Alu: *Asymmetron lucayanum*; Bbe: *Branchiostoma belcheri*; Bfl: *Branchiostoma floridae*; Bja: *Branchiostoma japonicum*; Bla: *Branchiostoma lanceolatum*.

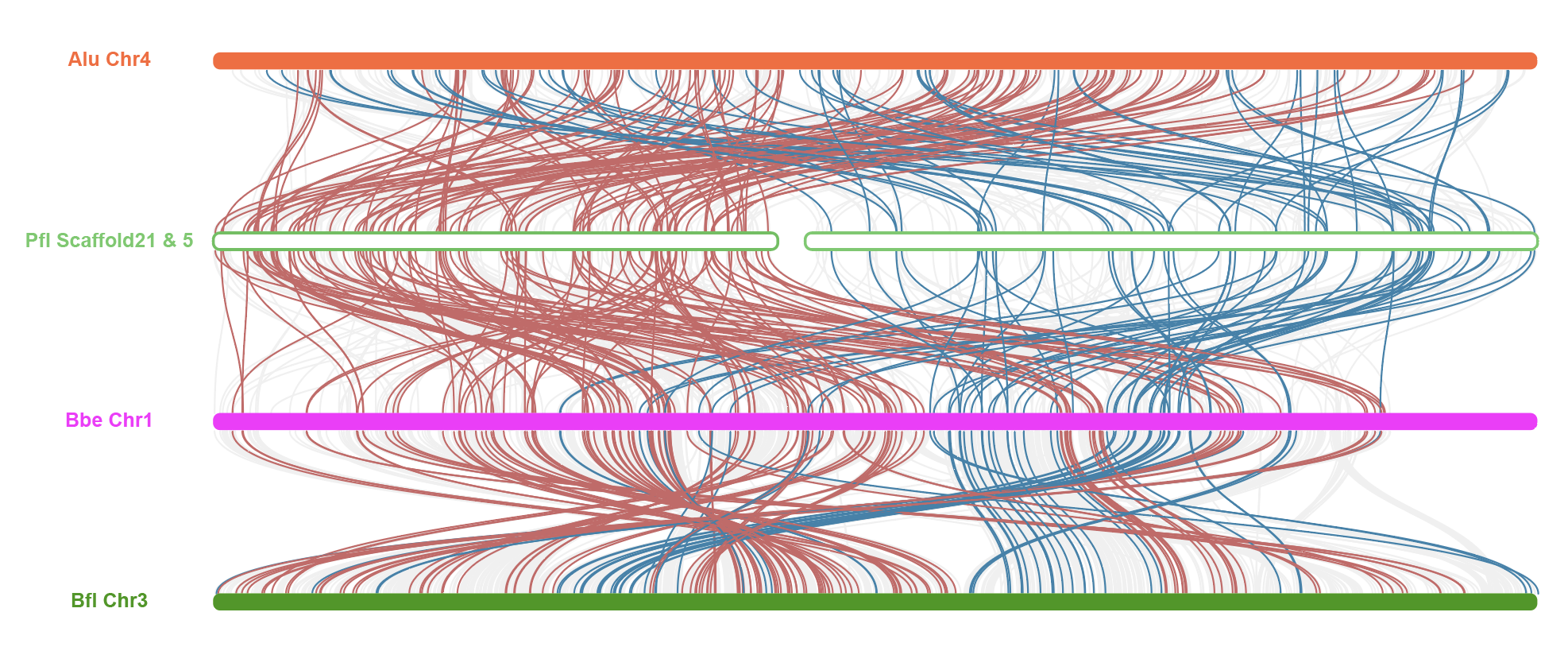

**Figure S6.** **The gene synteny comparison around the J2/C1 fusion in cephalochordates (Alu, Bbe, and Bfl) and hemichordates (Pfl).**

The hemichordate Pfl is used as a pre-fusion outgroup with the two scaffolds representing the J2 and C1 ancestral linkage groups. Alu: *Asymmetron lucayanum*, Bbe: *Branchiostoma belcheri*, Bfl: *Branchiostoma floridae*, Pfl: *Ptychodera flava*.

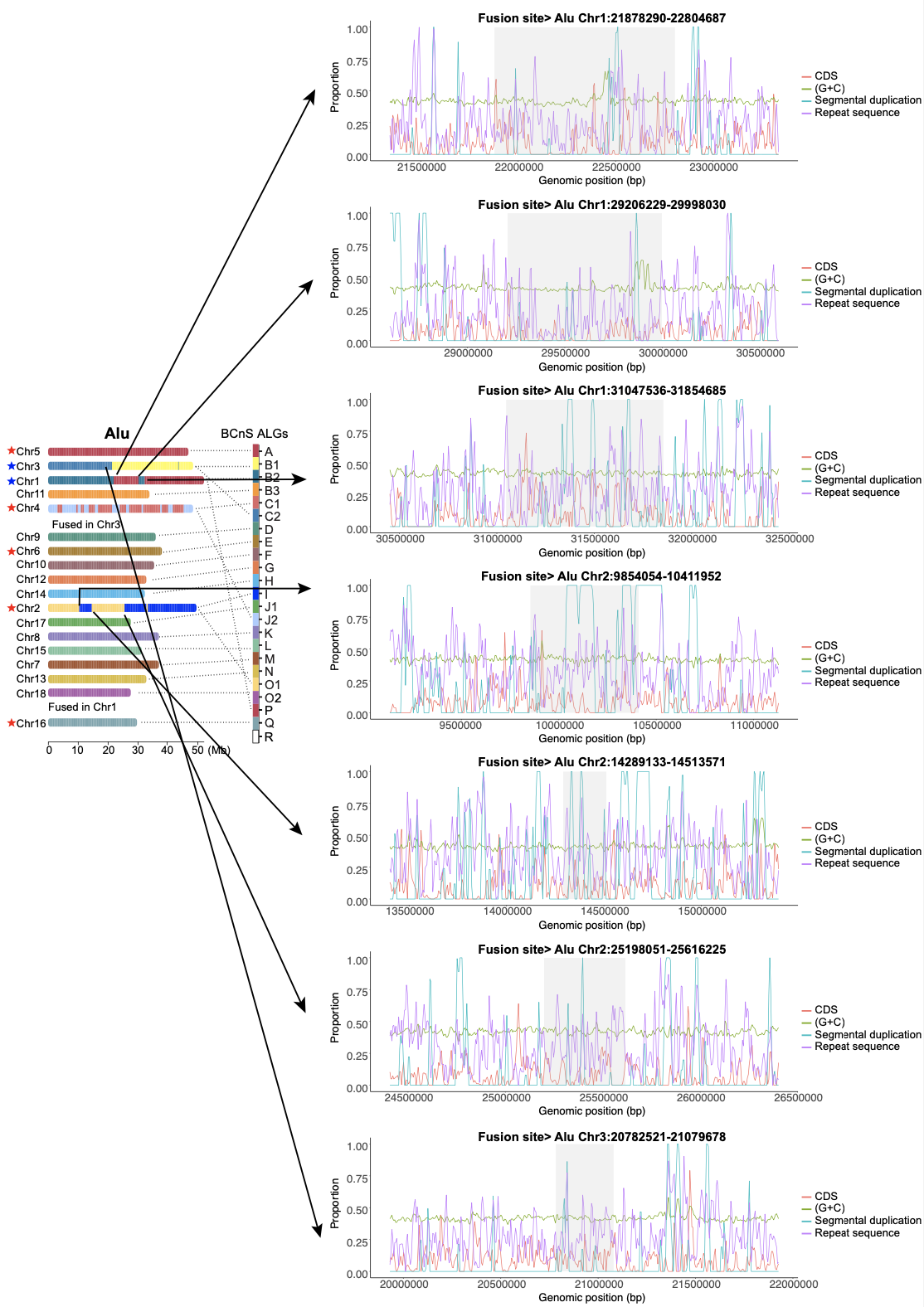

**Figure S7. Profiles of local genomic features at the cephalochordate fusion sites of BCnS ancestral linkage groups (ALGs) with Alu as an example.**

The examined genomic features include GC% as well as the window-based proportion of CDS, segmental duplications and repeat sequences. Alu: *Asymmetron lucayanum*.

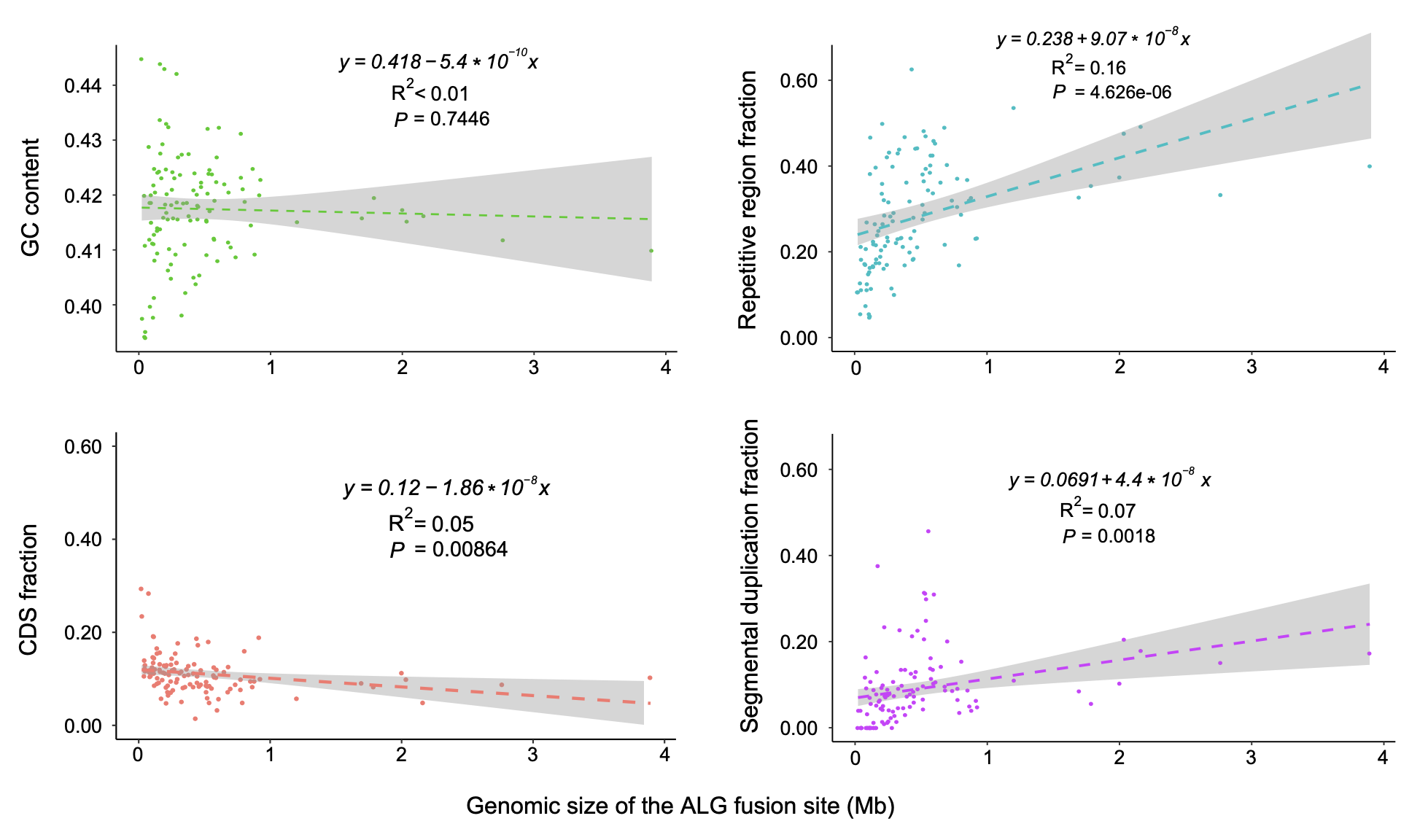

**Figure S8. Profiles of genomic features at the cephalochordate fusion sites of BCnS ancestral linkage groups (ALGs) with all five species combined.**

For genomic features such as GC bases, CDS regions, repetitive regions, and segmental duplication regions, scatterplots regarding their respective proportion at each fusion sites against the size of the corresponding fusion site were plotted. Pearson correlation was also used to evaluate such relationships.

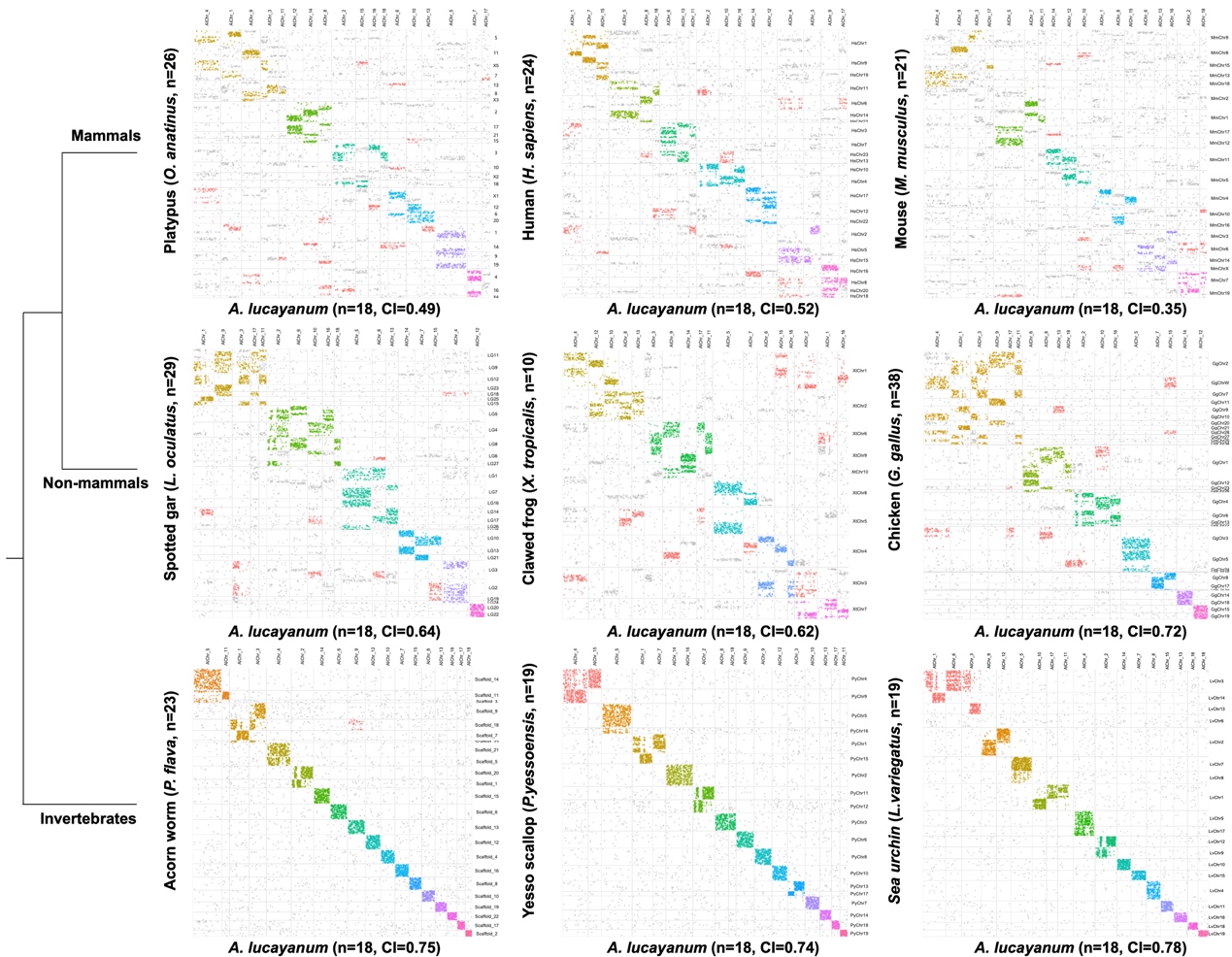

**Figure S9.** **Macrosynteny conservation between the cephalochordate *Asymmetron lucayanum* and representative vertebrate and invertebrate species.**

In each individual Oxford grid dotplots, orthologs inferred to the same linkage group (based on macrosyntR’s greedy clustering algorithm) was colored in the same color. The colors used in each individual Oxford grid dotplots are pairwise-comparison-specific and nontransferable between different comparison group. n: chromosome number; CI: macrosynteny conservation index.

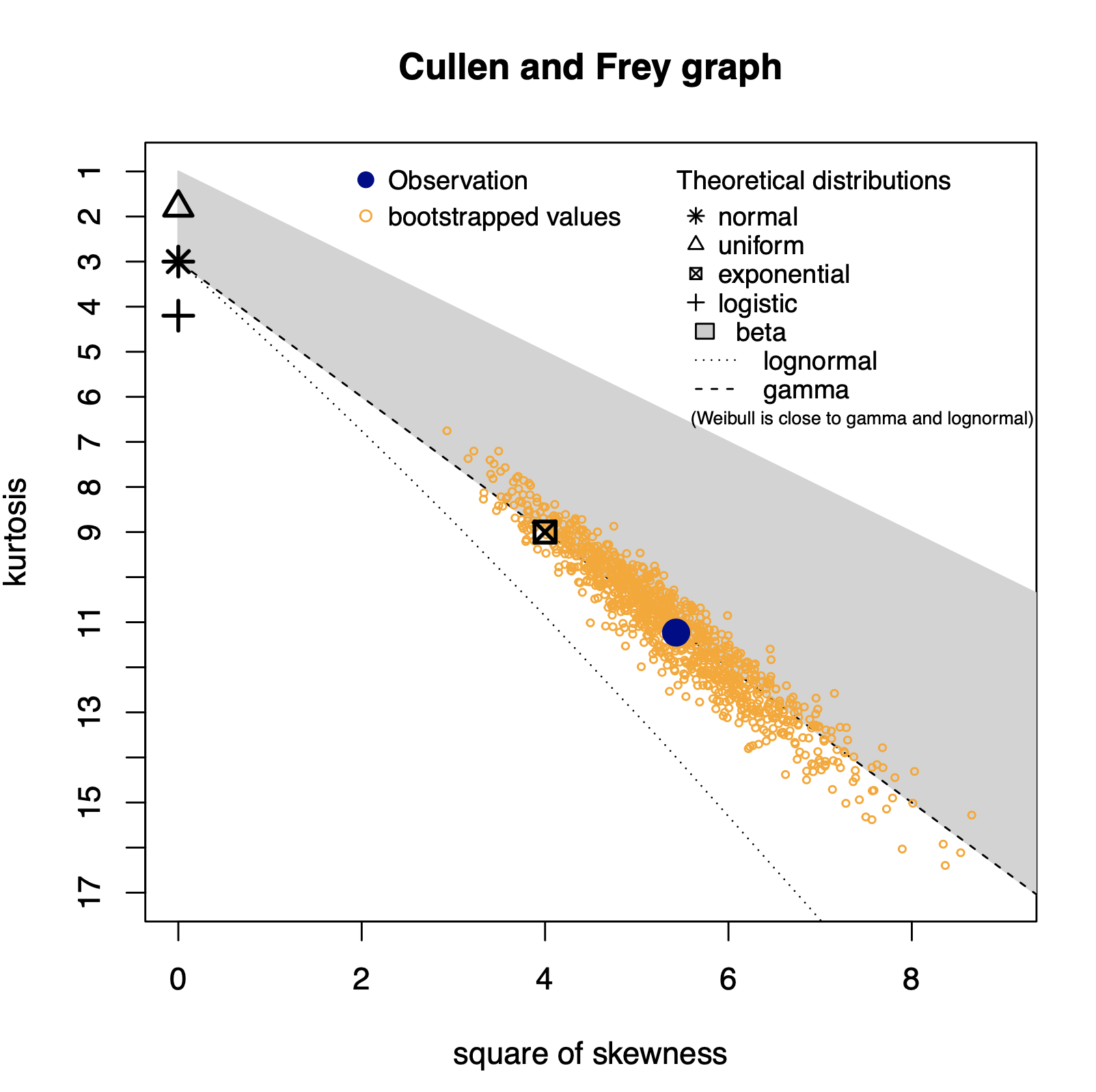

**Figure S10. The Cullen and Frey diagnostic graph to evaluate potential statistical distribution choice for fitting with the observed cephalochordate microsynteny block size distribution.**

A total of 1000 bootstrap permutations were used in this analysis. This plot suggests log normal, gamma, Weibull, and exponential distributions as potential candidates for fitting with the observed distribution.

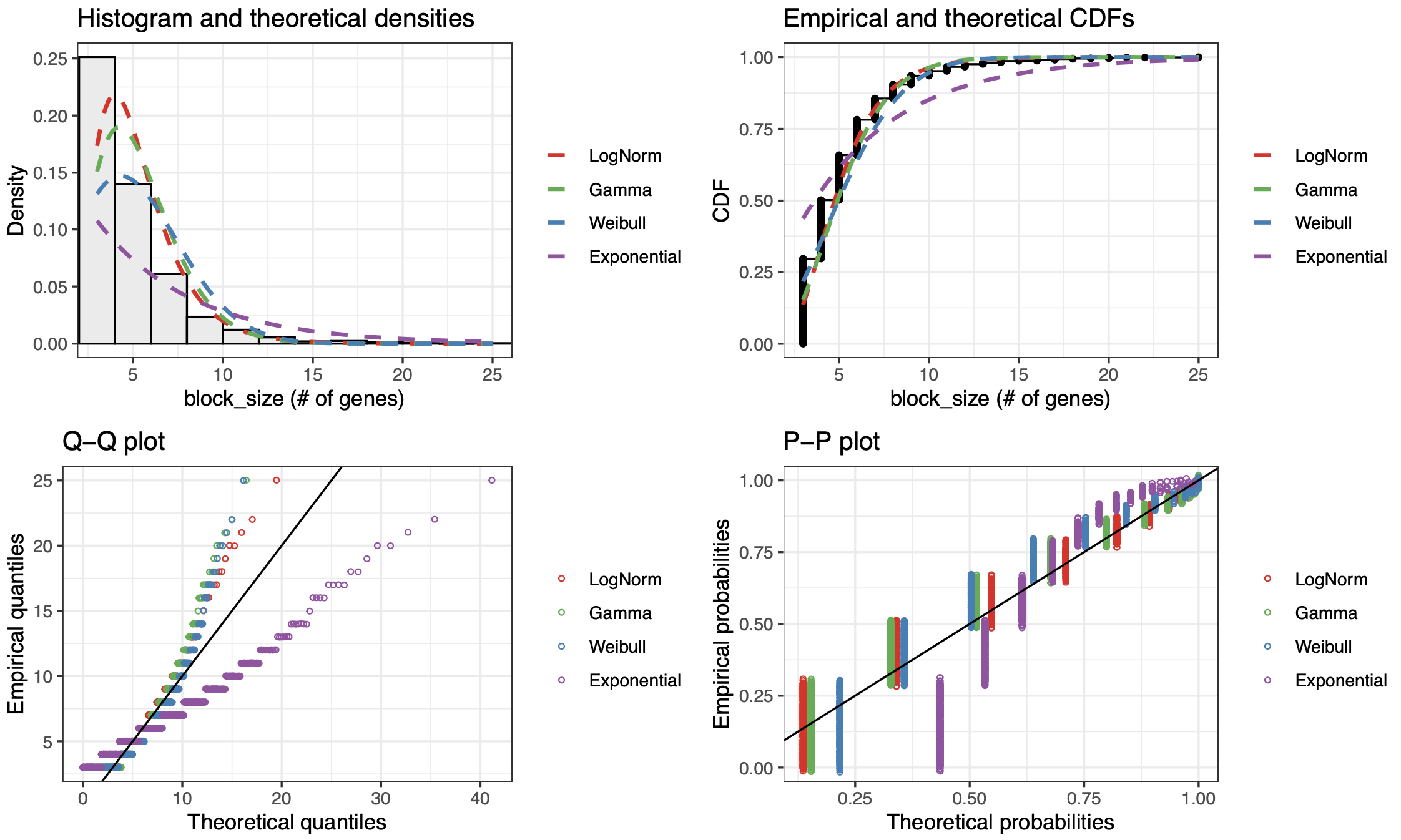

**Figure S11.** **Statistical distribution model fitting for the observed cephalochordate microsynteny block size distribution.**

The log normal, gamma, Weibull, and exponential distributions were evaluated, with the log normal distribution appears to be the one that is most close to the real data. CDF: cumulative distribution function.

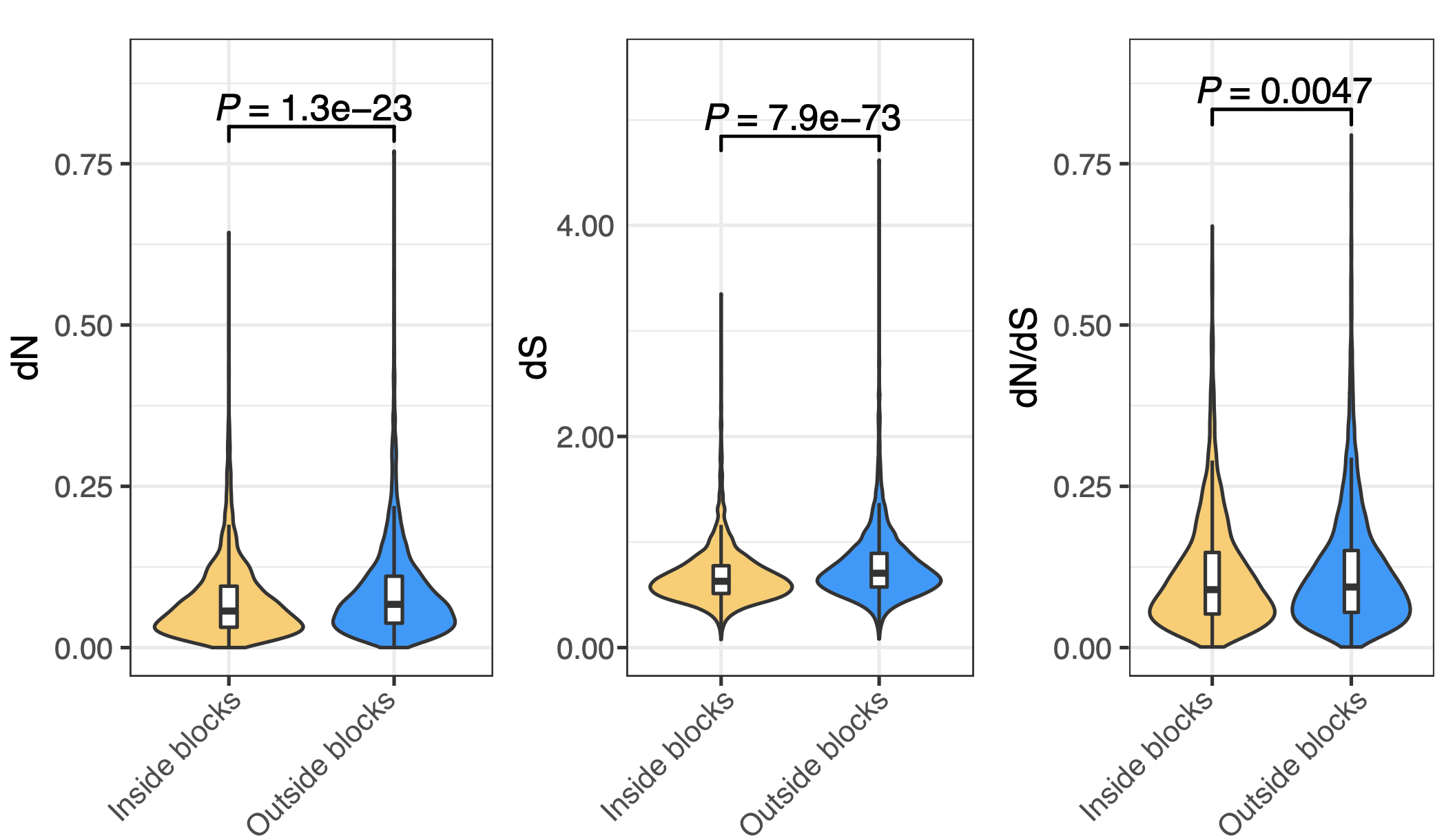

**Figure S12.** **Comparison of molecular evolution rates between genes within and outside the cephalochordate microsynteny blocks.**

Only one-to-one ortholog genes were used for this calculation. dN: nonsynonymous substitution rate. dS: synonymous substitution rate. dN/dS: nonsynonymous-to-synonymous substitution rate.

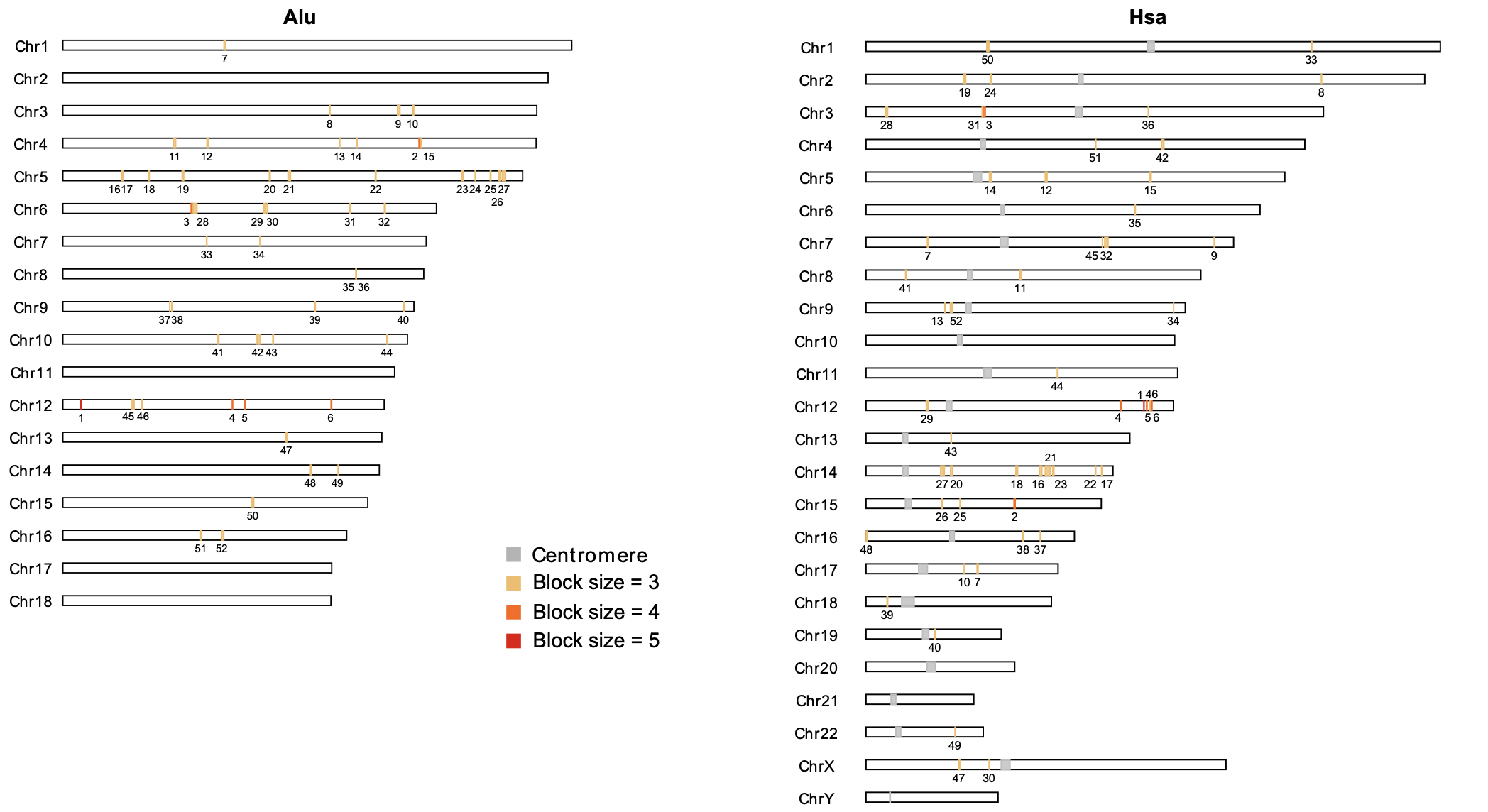

**Figure S13. Chromosomal distribution of microsynteny blocks shared between cephalochordates and human.**

The blocks are ordered by the block size (number of syntenic ortholog genes enclosed). Alu: *Asymmetron lucayanum*; Hsa: *Homo sapiens*.

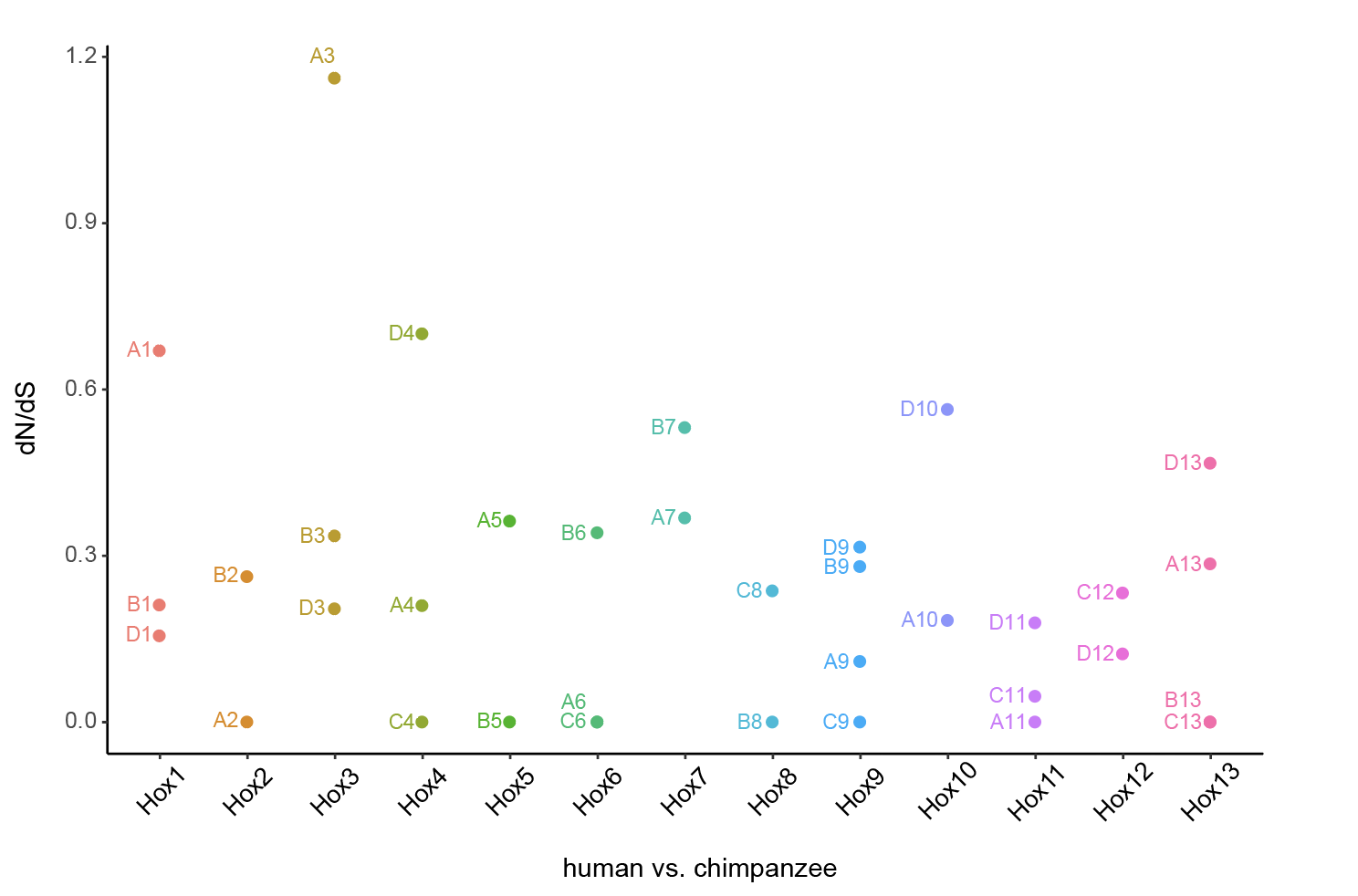

**Figure S14. The dN/dS values of *Hox* genes in the human-chimpanzee comparison.**

The A, B, C, D prefixes denote the four vertebrate *Hox* paralogons after 2R-WGDs.

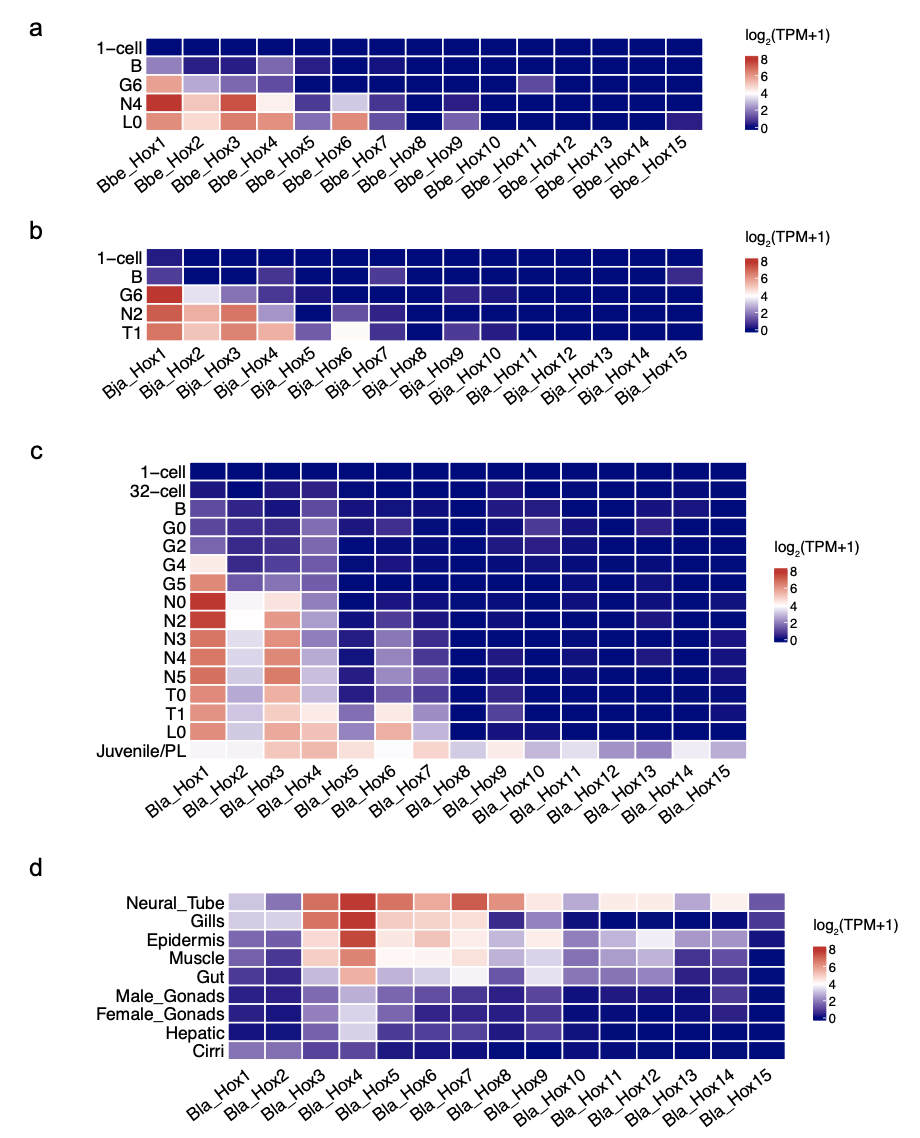

**Figure S15. The *Hox* gene expression in other *Branchiostoma* species.** a-c. The stage- *Hox* expression pattern for *B. belcheri* (Bbe), *B.* *japonicum* (Bja)*,* and *B. lanceolatum* (Bla) respectively. d. The tissue-specific *Hox* expression pattern for *B. lanceolatum* (Bla).

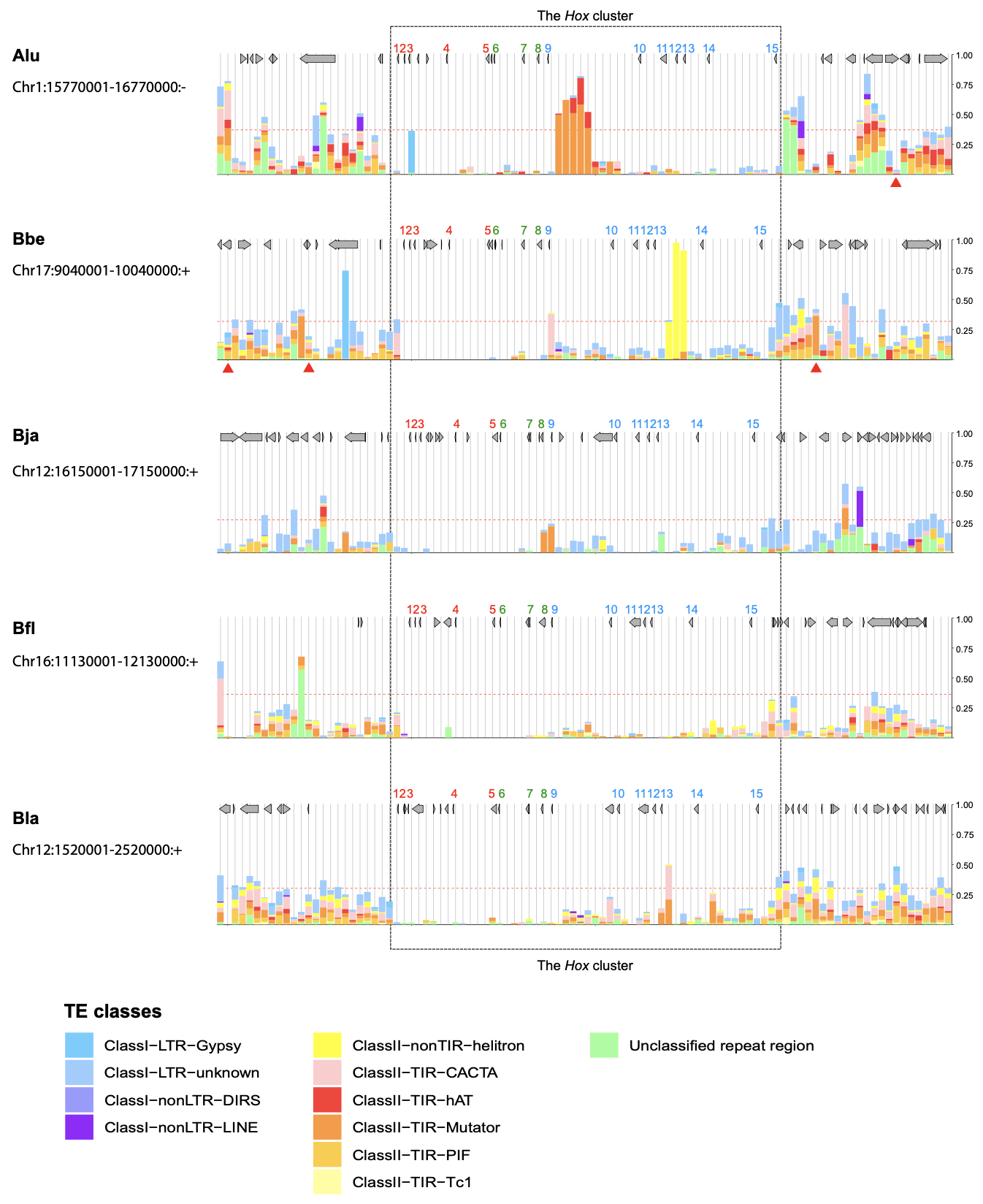

**Figure S16. The abundance of different classes of repetitive sequences in the *Hox* cluster of the five cephalochordate species.**

Alu: *Asymmetron lucayanum*; Bbe: *Branchiostoma belcheri*; Bfl: *Branchiostoma floridae*; Bja: *Branchiostoma japonicum*; Bla: *Branchiostoma lanceolatum*.

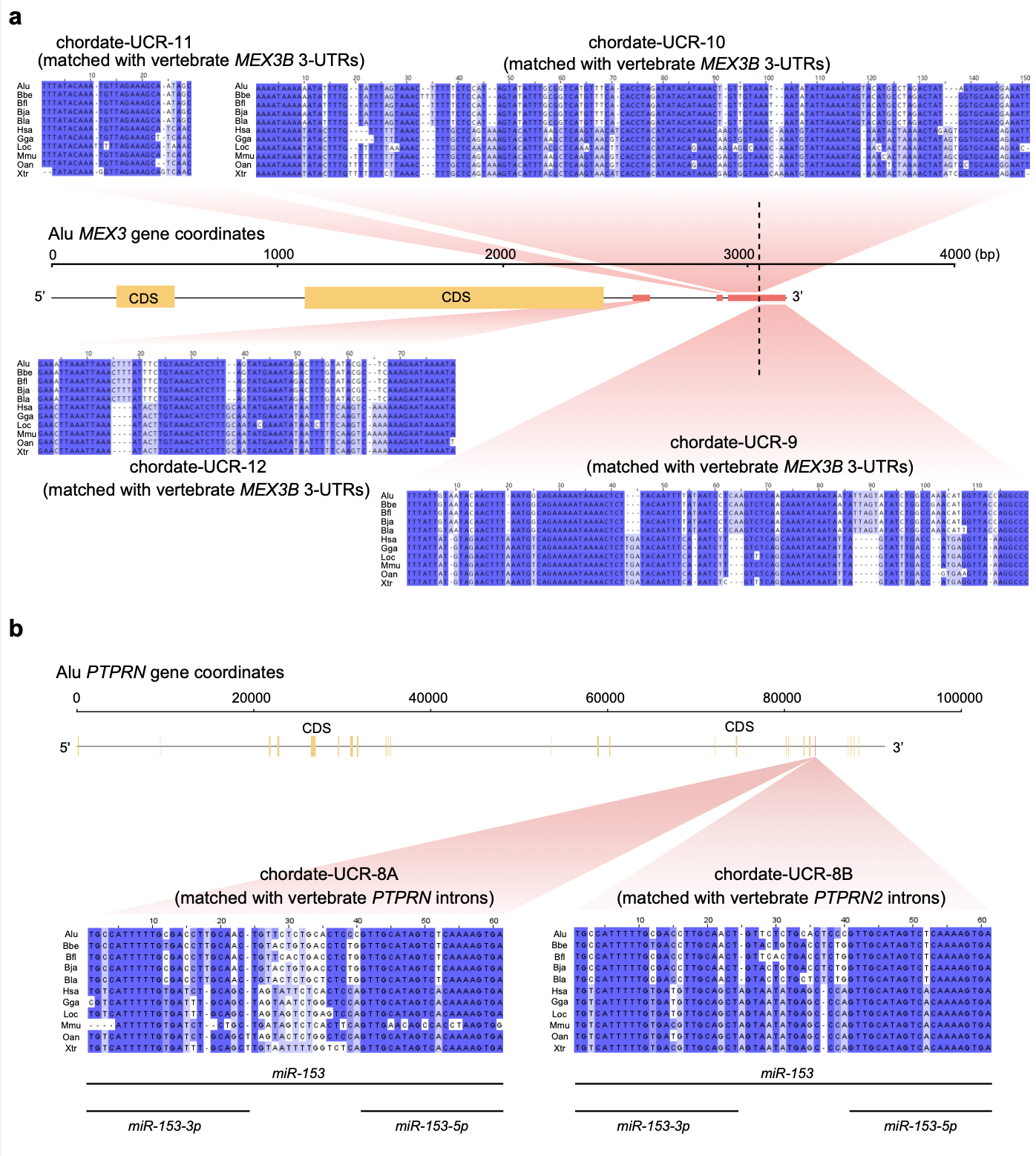

**Figure S17. Examples of chordate ultra-conserved regions (UCRs).**

(a) The chordate-UCRs associated with *MEX3*. (b) The chordate-UCRs associated with *PTPRN/PTPRN2*, which matched with the highly conserved noncoding RNA gene *miR-153*. Alu: *Asymmetron lucayanum*; Bbe: *Branchiostoma belcheri*; Bfl: *Branchiostoma floridae*; Bja: *Branchiostoma japonicum*; Bla: *Branchiostoma lanceolatum*; Gga: *Gallus gallus*; Hsa: *Homo sapiens*; Loc: *Lepisosteus oculatus*; Mmu: *Mus musculus*; Oan: *Ornithorhynchus anatinus*; Xtr: *Xenopus tropicalis*.

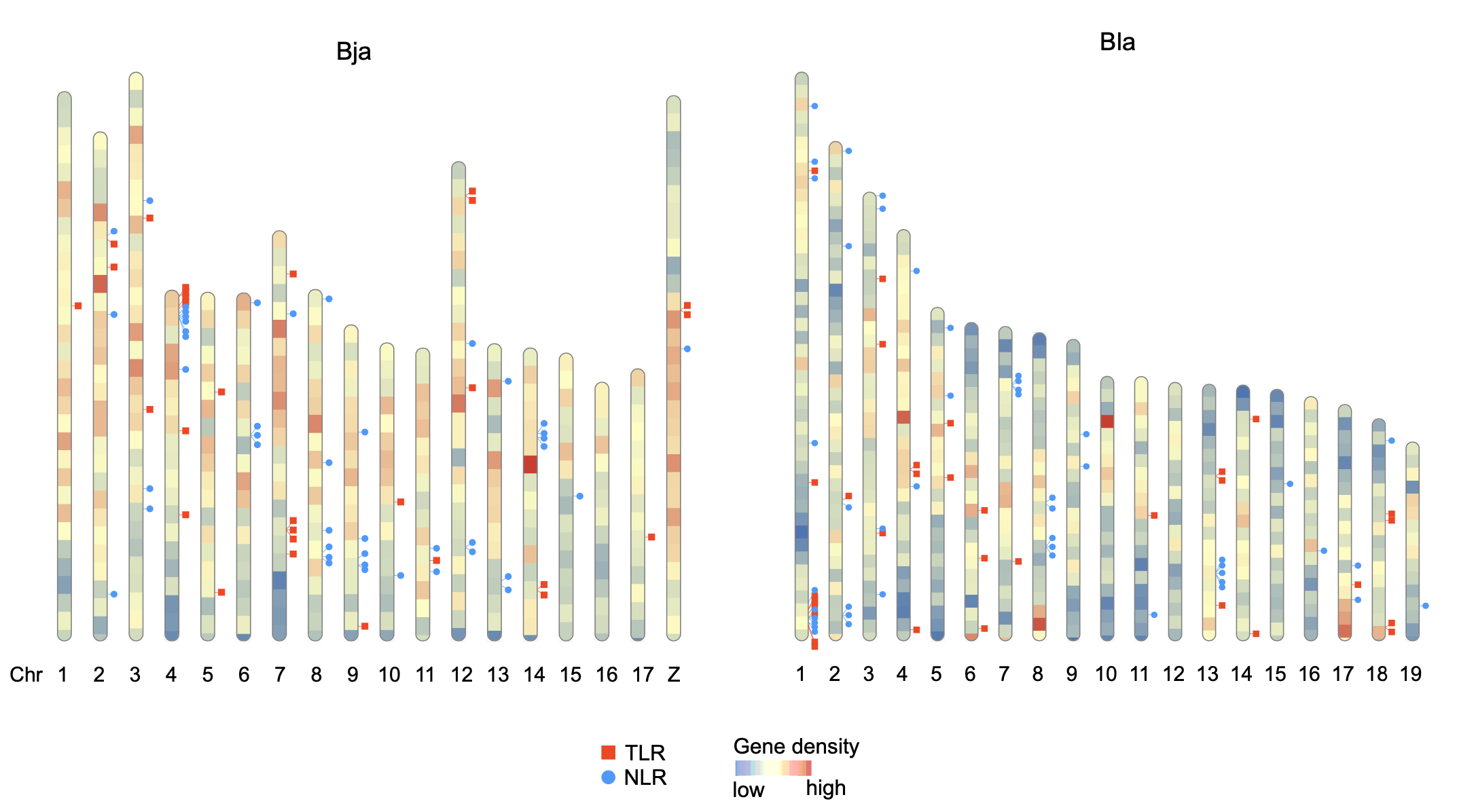

**Figure S18.** **The chromosomal distribution of *TLR* and *NLR* genes in Bja and Bla.**

The *TLR* and *NLR* genes are indicated by red and blue respectively. Bja: *Branchiostoma japonicum*; Bla: *Branchiostoma lanceolatum*.

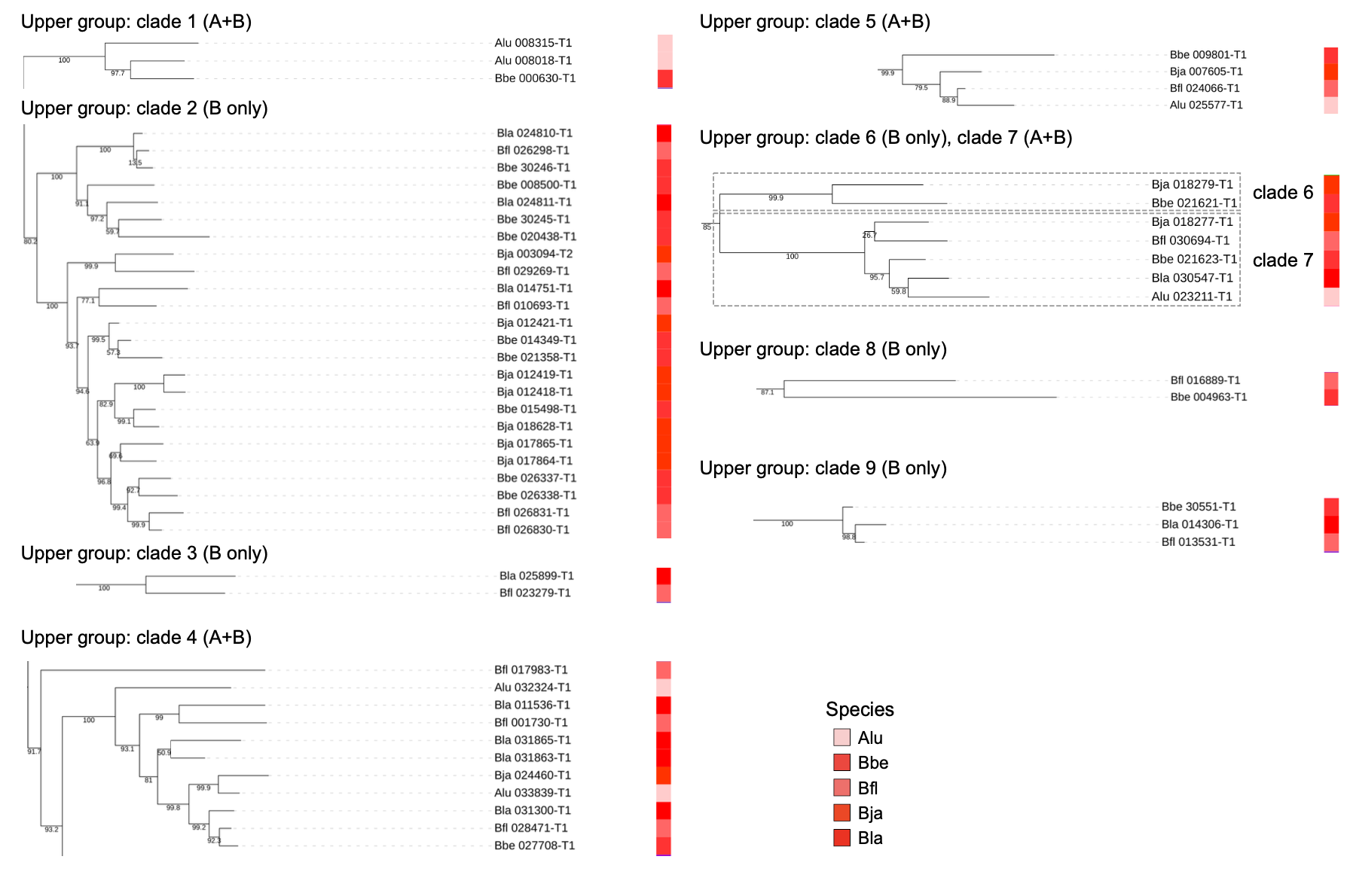

**Figure S19. Cephalochordate *TLR* clades in the upper half of the *TLR* gene tree shown in Figure 6a.**

Alu: *Asymmetron lucayanum*; Bbe: *Branchiostoma belcheri*; Bfl: *Branchiostoma floridae*; Bja: *Branchiostoma japonicum*; Bla: *Branchiostoma lanceolatum*. A+B: clade shared by both *Asymmetron* and *Branchiostoma*. B only: clade only containing *Branchiostoma* *TLRs*.

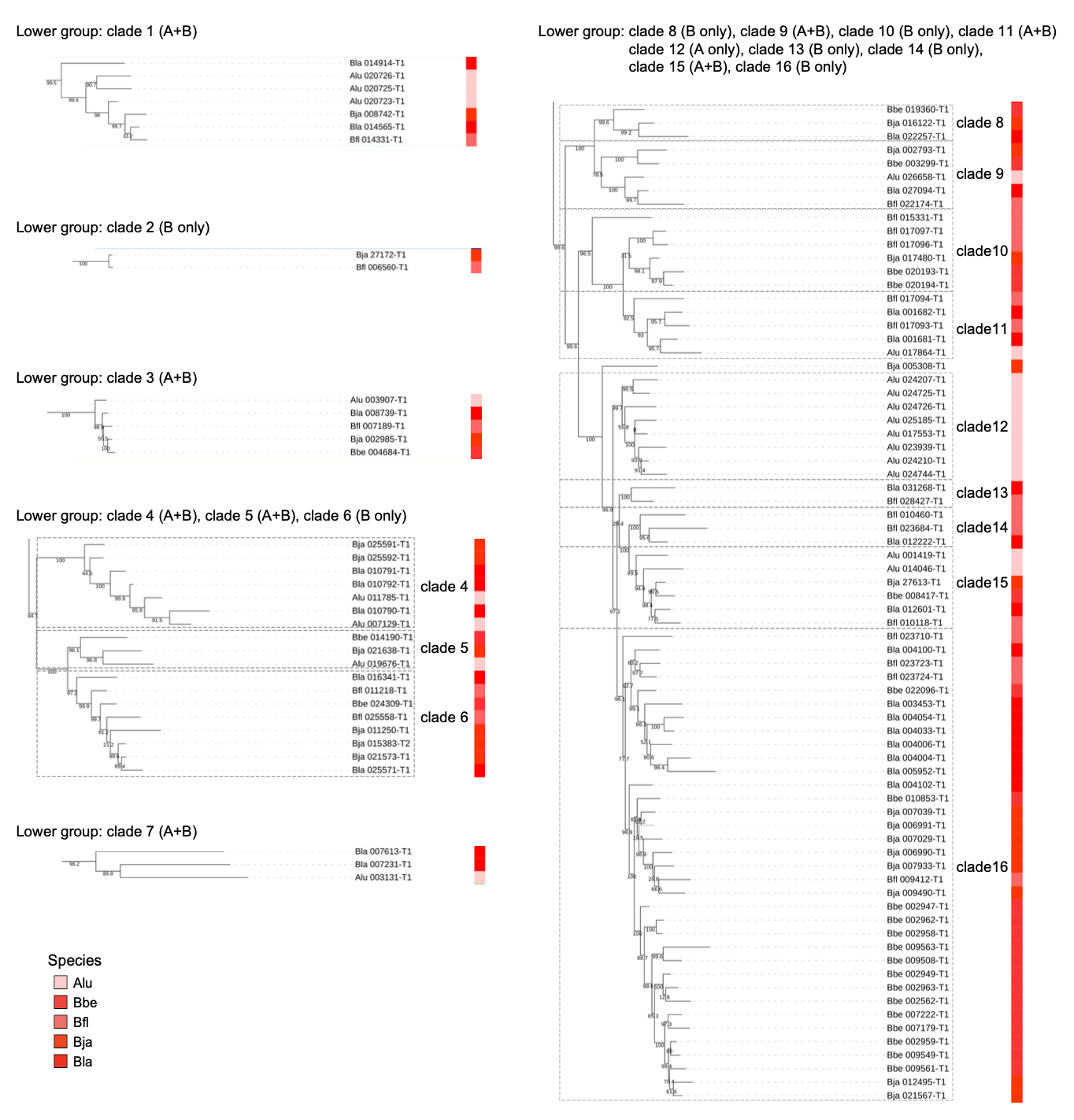

**Figure S20. Cephalochordate *TLR* clades in the lower half of the *TLR* gene tree shown in Figure 6a.**

Alu: *Asymmetron lucayanum*; Bbe: *Branchiostoma belcheri*; Bfl: *Branchiostoma floridae*; Bja: *Branchiostoma japonicum*; Bla: *Branchiostoma lanceolatum*. A+B: clade shared by both *Asymmetron* and *Branchiostoma*. B only: clade only containing *Branchiostoma* *TLRs*.

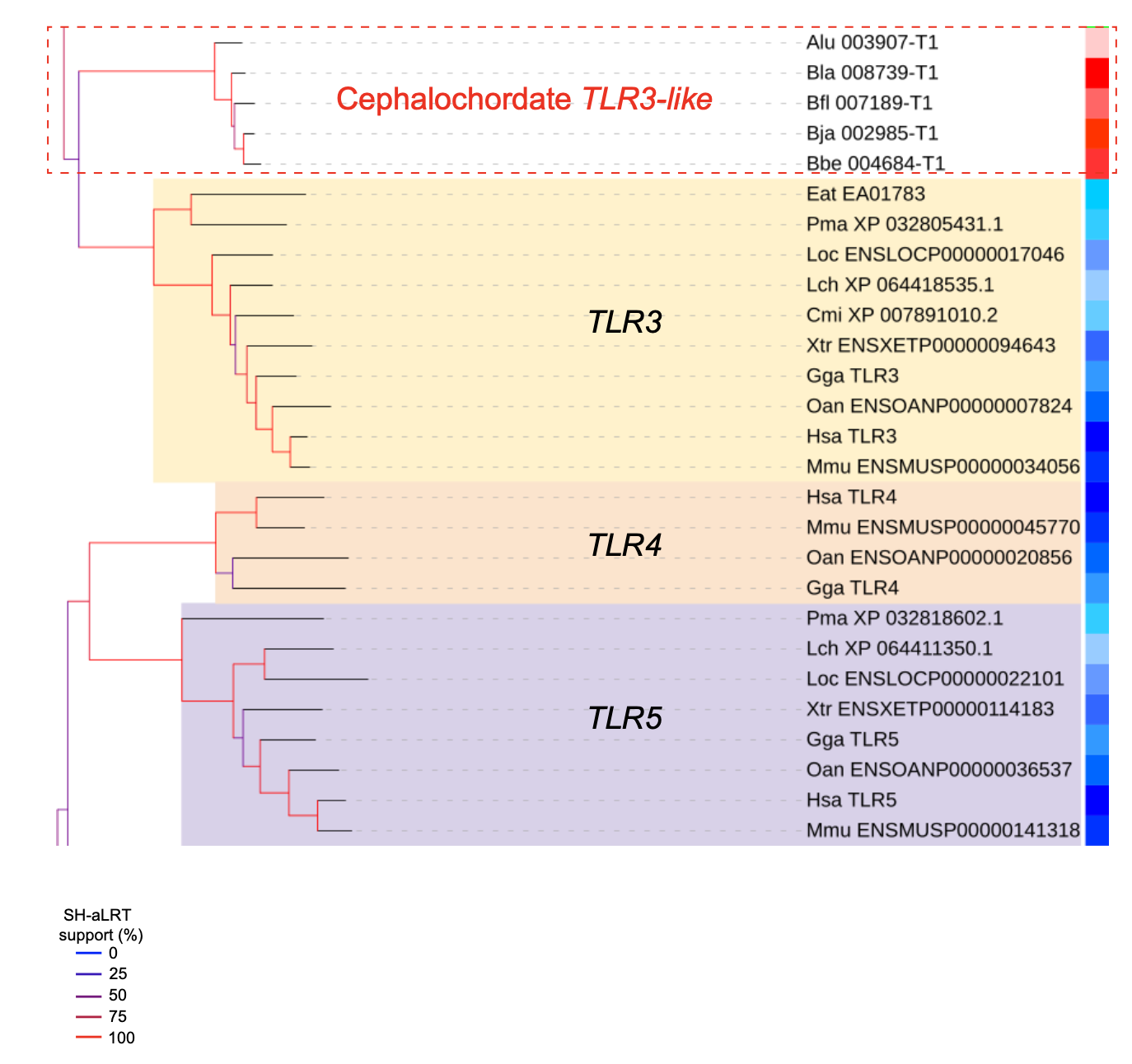

**Figure S21****. The *TLR3*, *TLR4*, and *TLR5* clades in the *TLR* phylogenetic tree.**

Alu: *Asymmetron lucayanum*; Bbe: *Branchiostoma belcheri*; Bfl: *Branchiostoma floridae*; Bja: *Branchiostoma japonicum*; Bla: *Branchiostoma lanceolatum*; Cmi: *Callorhinchus milii*; Eat: *Eptatretus atami*; Gga: *Gallus gallus*; Hsa: *Homo sapiens*; Lch: *Latimeria chalumnae*; Loc: *Lepisosteus oculatus*; Mmu: *Mus musculus*; Oan: *Ornithorhynchus anatinus*; Pma: *Petromyzon marinus*; Xtr: *Xenopus tropicalis*.

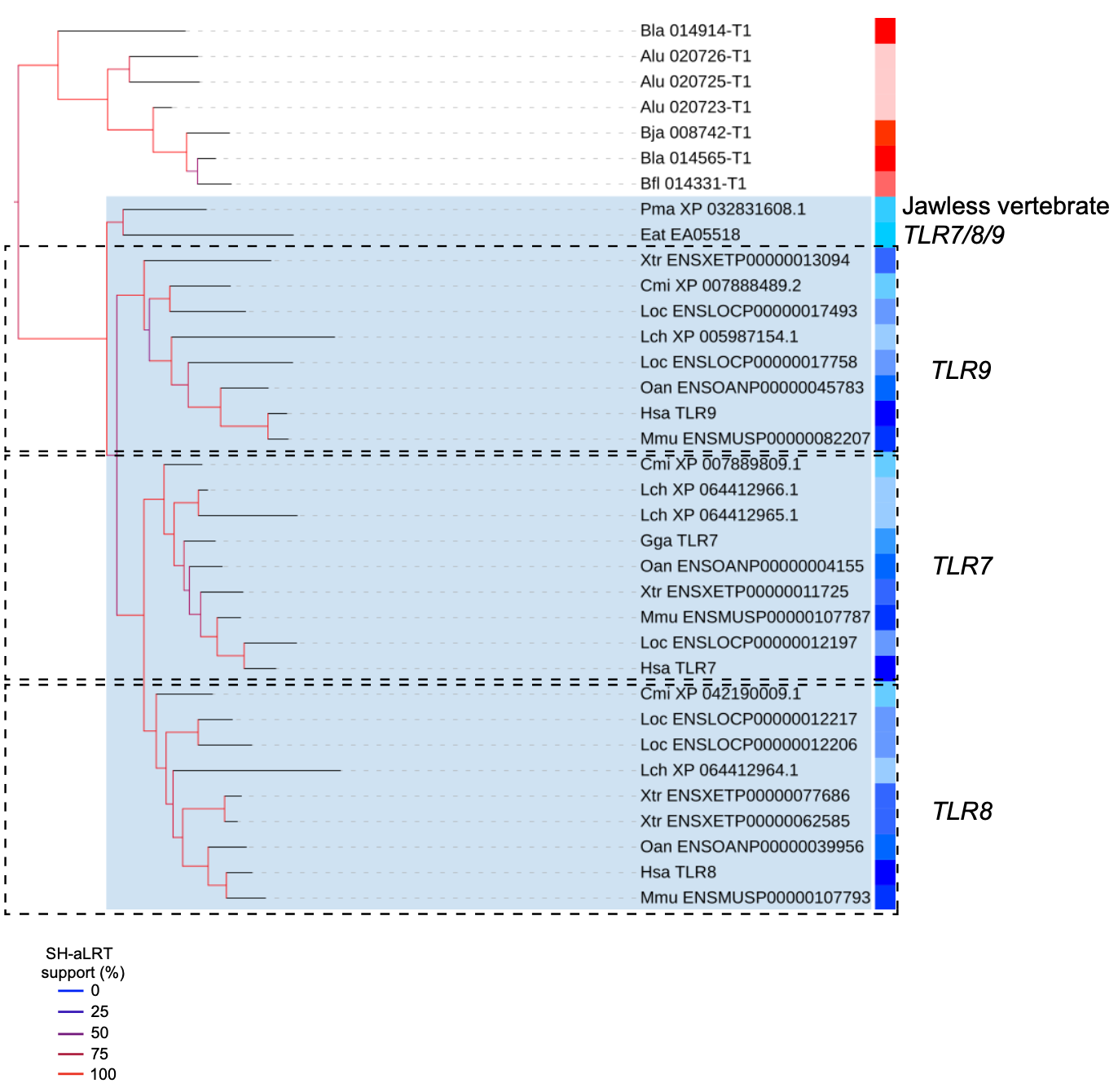

**Figure S22. The *TLR7*, *TLR8*, and *TLR9* clades in the *TLR* phylogenetic tree.**

Alu: *Asymmetron lucayanum*; Bfl: *Branchiostoma floridae*; Bja: *Branchiostoma japonicum*; Bla: *Branchiostoma lanceolatum*; Cmi: *Callorhinchus milii*; Eat: *Eptatretus atami*; Gga: *Gallus gallus*; Hsa: *Homo sapiens*; Lch: *Latimeria chalumnae*; Loc: *Lepisosteus oculatus*; Mmu: *Mus musculus*; Oan: *Ornithorhynchus anatinus*; Pma: *Petromyzon marinus*; Xtr: *Xenopus tropicalis*.

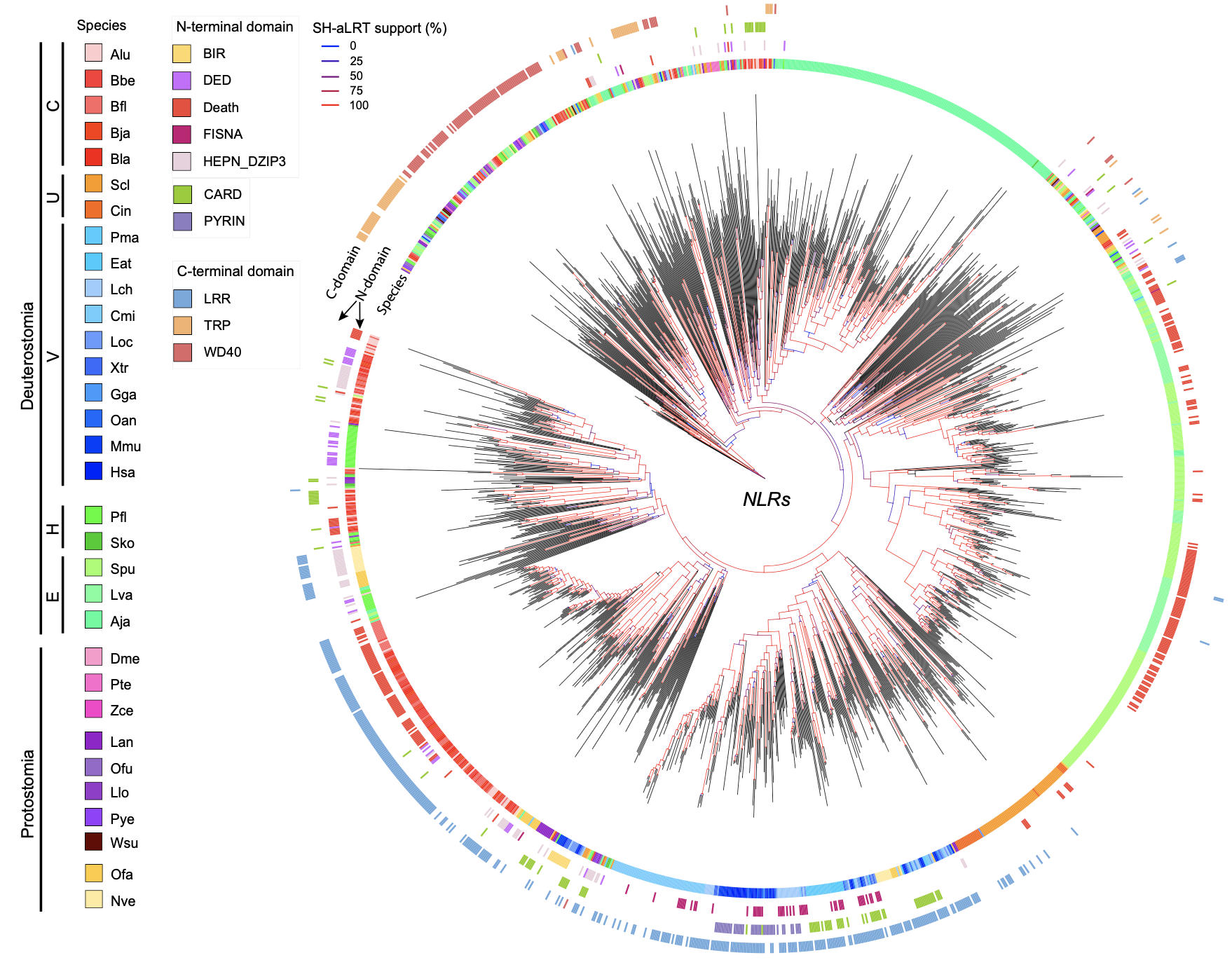

**Figure S23. The phylogenetic tree of *NLR* genes.**

The information regarding each plotted gene’s species origin, N-terminal domains, and C-terminal domains are depicted in different layer of cycles. Given that some genes processes more than one characteristic N-terminal domains, two layers of cycles were used to reflect their N-terminal organizations. C: Cephalochordata, U: Urochordata, V: Vertebrata, H: Hemichordata, E: Echinodermata.

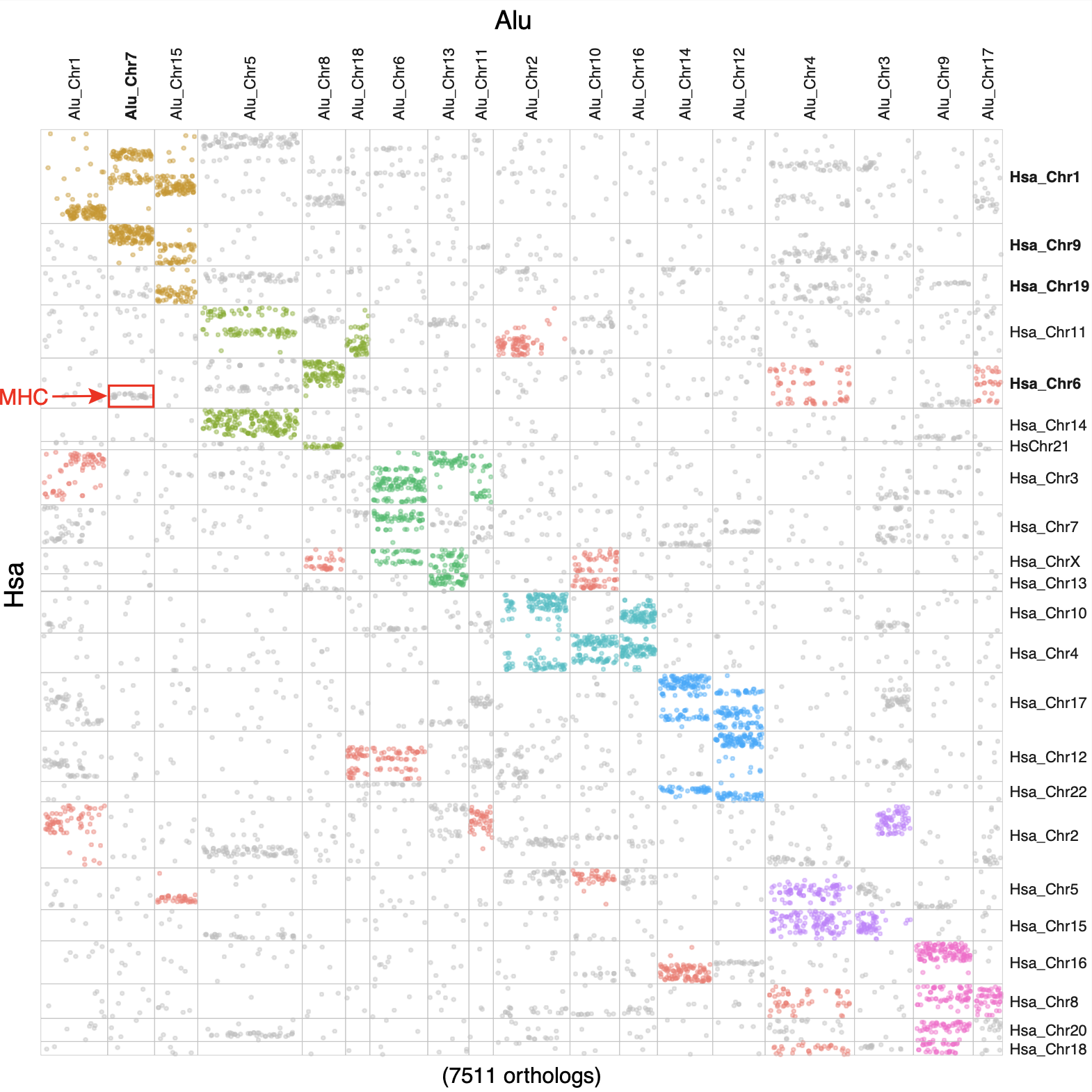

**Figure S24. The Oxford grid dotplot of *A. lucayanum* (Alu)-human (Hsa) comparison regarding the macrosynteny conservation of their shared ortholog genes.**

Orthologs inferred to the same linkage group (based on macrosyntR’s greedy clustering algorithm) was colored in the same color. Gene representing the human MHC locus are highlighted with a red box.

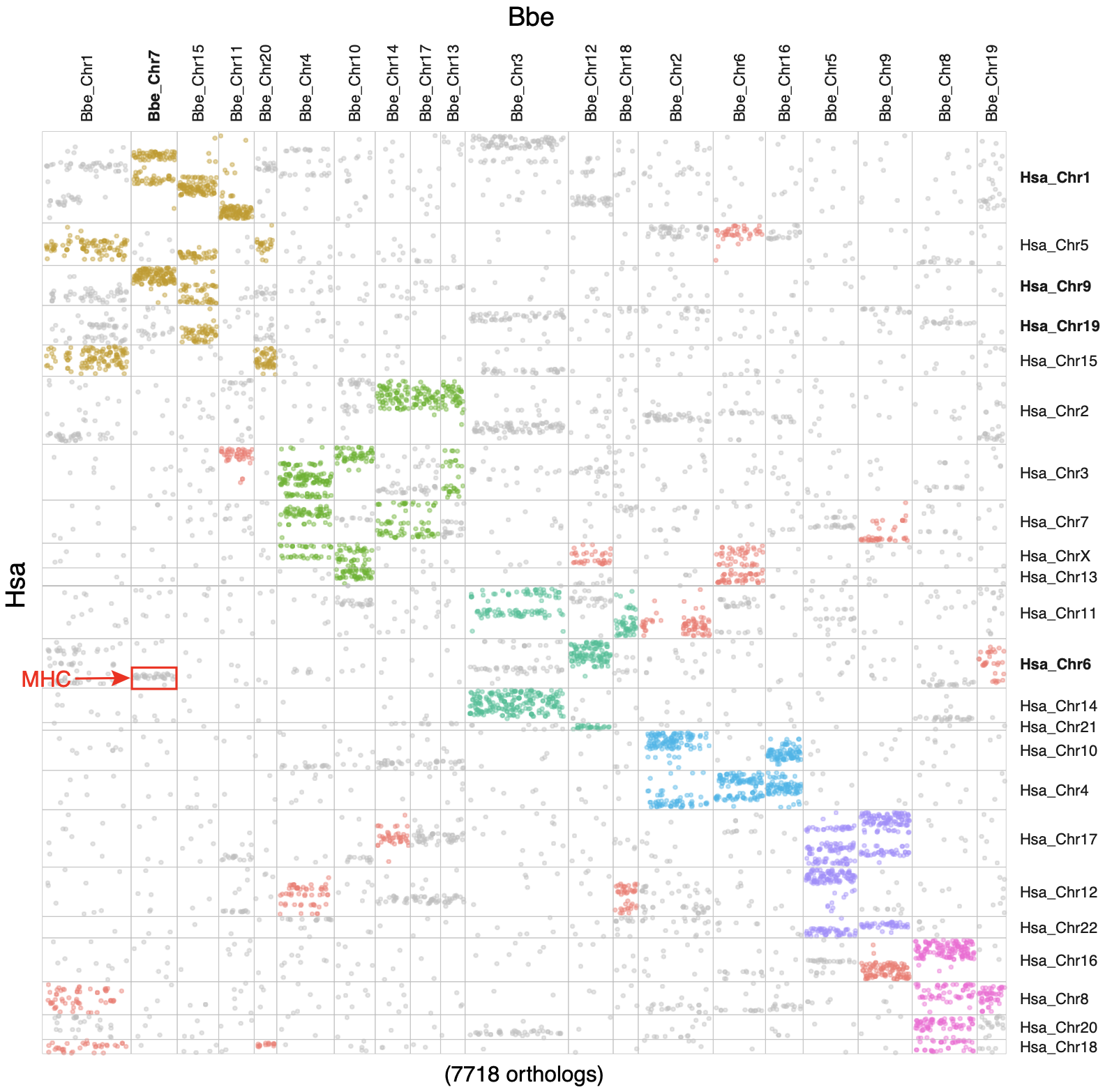

**Figure S25. The Oxford grid dotplot of *B. belcheri* (Bbe)-human (Hsa) comparison regarding the macrosynteny conservation of their shared ortholog genes.**

Orthologs inferred to the same linkage group (based on macrosyntR’s greedy clustering algorithm) was colored in the same color. Gene representing the human MHC locus are highlighted with a red box.

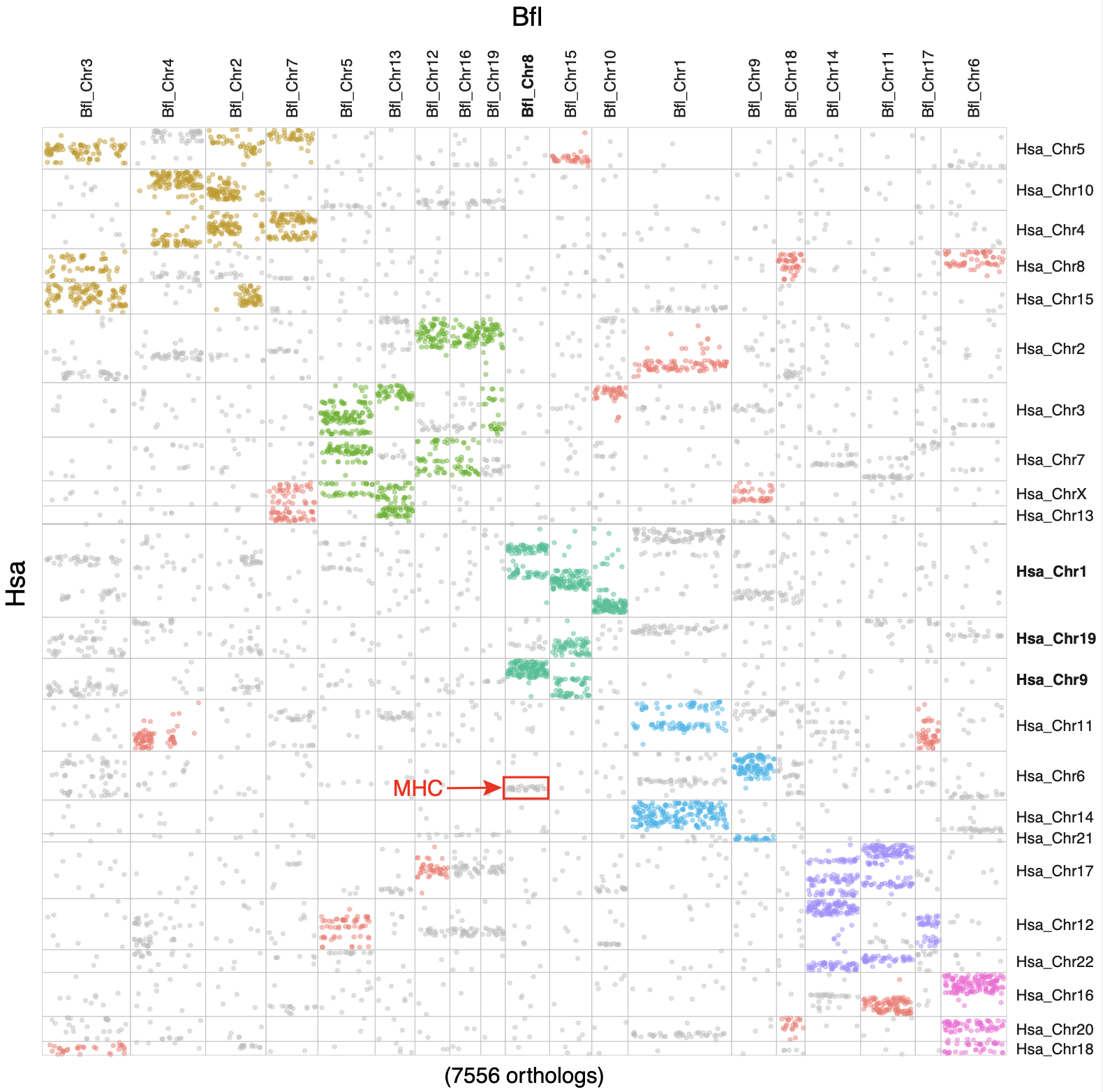

**Figure S26. The Oxford grid dotplot of *B. floridae* (Bfl)-human (Hsa) comparison regarding the macrosynteny conservation of their shared ortholog genes.**

Orthologs inferred to the same linkage group (based on macrosyntR’s greedy clustering algorithm) was colored in the same color. Gene representing the human MHC locus are highlighted with a red box.

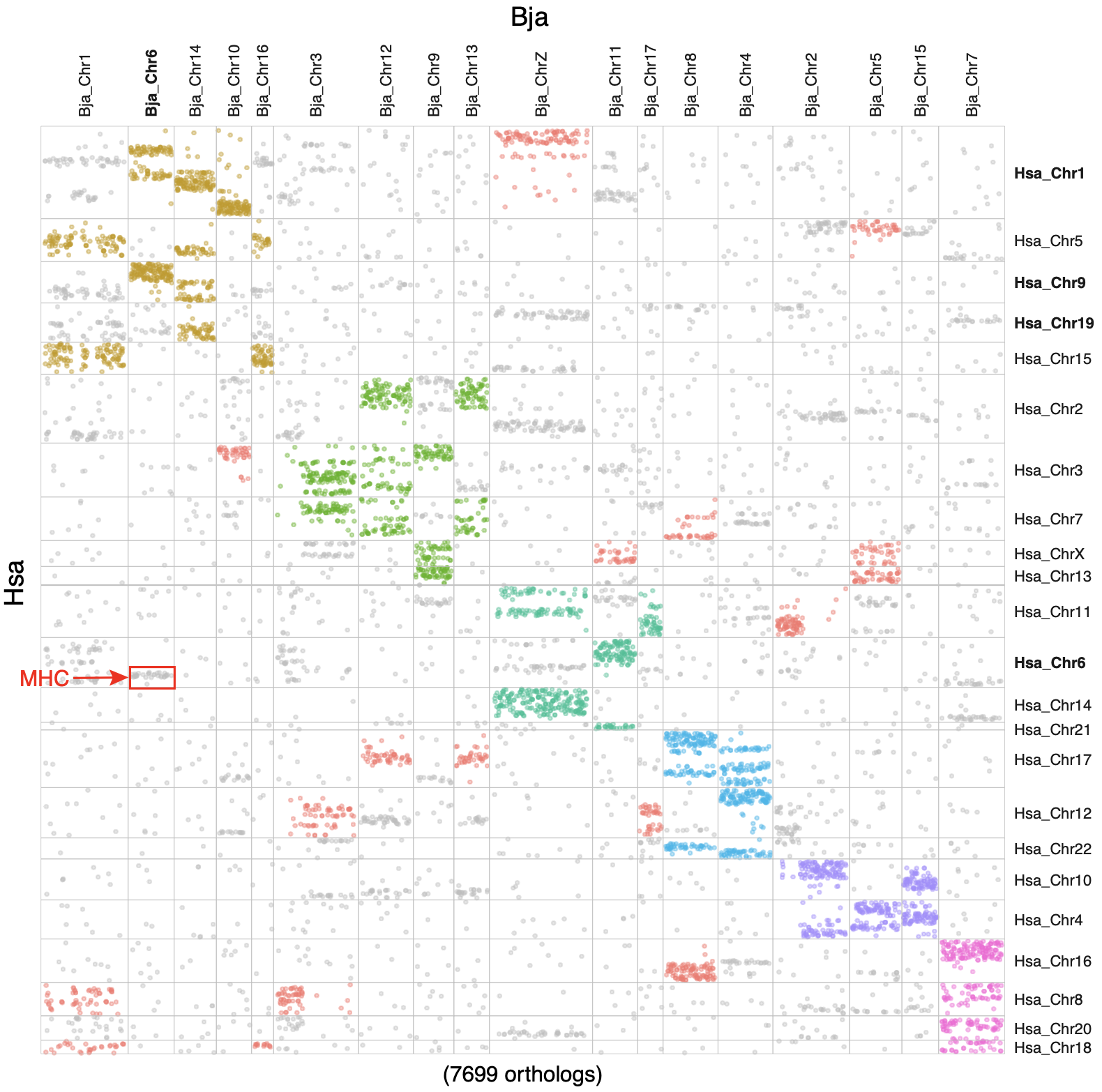

**Figure S27. The Oxford grid dotplot of *B. japonicum* (Bja)-human (Hsa) comparison regarding the macrosynteny conservation of their shared ortholog genes.**

Orthologs inferred to the same linkage group (based on macrosyntR’s greedy clustering algorithm) was colored in the same color. Gene representing the human MHC locus are highlighted with a red box.

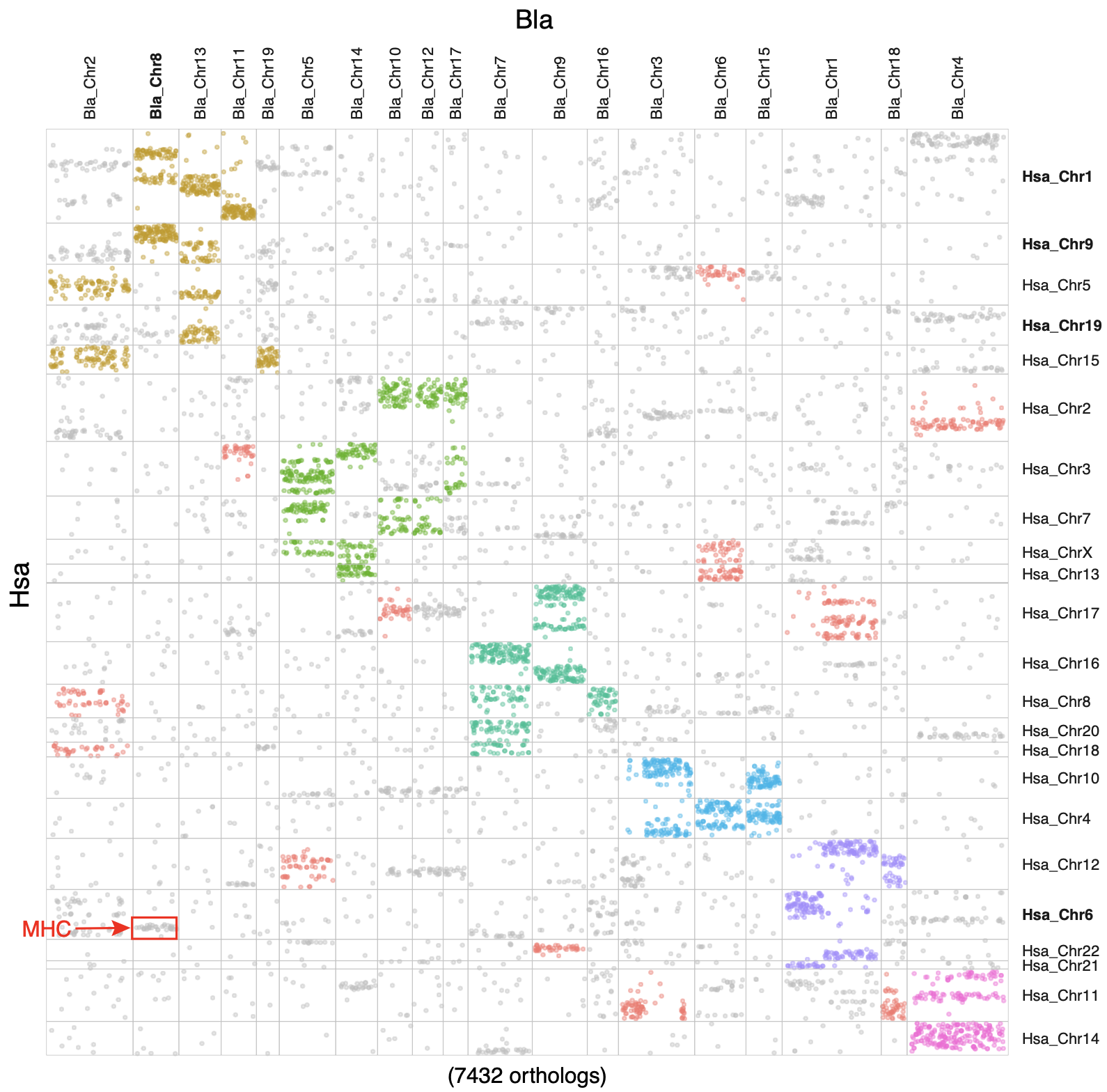

**Figure S28. The Oxford grid dotplot of *B. lanceolatum* (Bla)-human (Hsa) comparison regarding the macrosynteny conservation of their shared ortholog genes.**

Orthologs inferred to the same linkage group (based on macrosyntR’s greedy clustering algorithm) was colored in the same color. Gene representing the human MHC locus are highlighted with a red box.

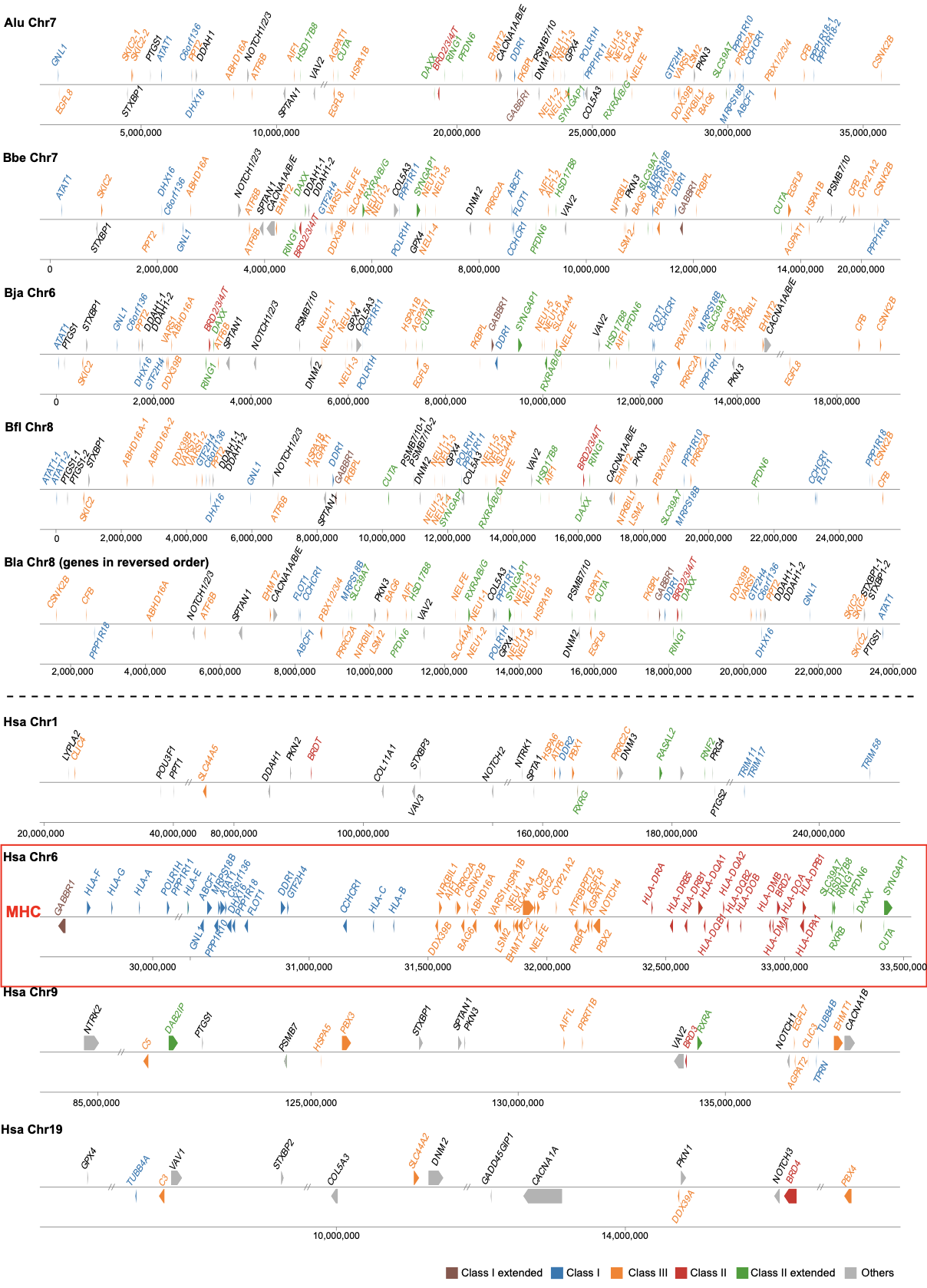

**Figure S29. The chromosomal organization of the protoMHC and MHC homolog genes shared between cephalochordates and human.**

Genes are colored based on their human MHC ortholog’s classification groups (e.g., Class I, Class II, Class III, Class I extended, Class II extended). Alu: *Asymmetron lucayanum*; Bbe: *Branchiostoma belcheri*; Bfl: *Branchiostoma floridae*; Bja: *Branchiostoma japonicum*; Bla: *Branchiostoma lanceolatum*; Hsa: *Homo sapiens*.

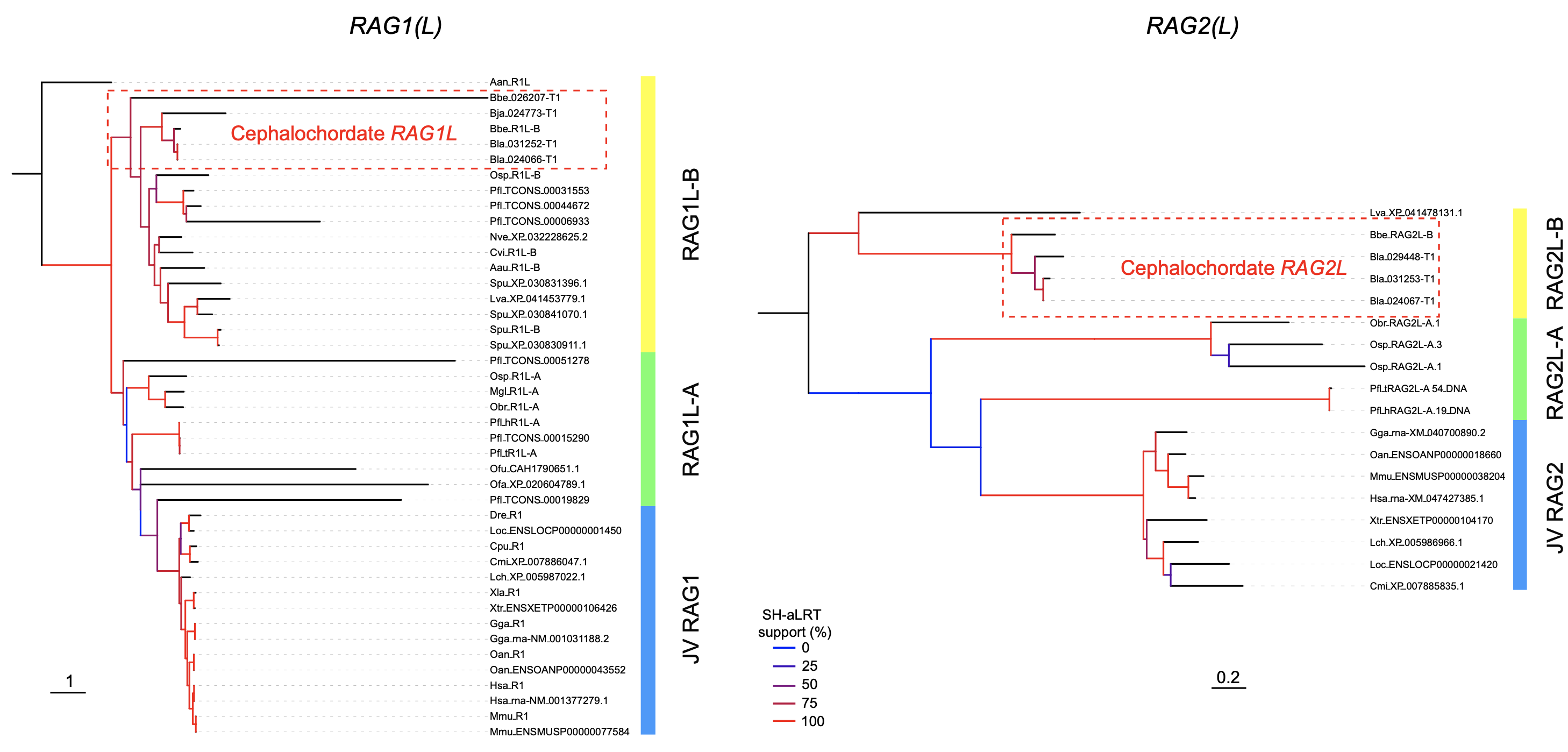

**Figure S30. The phylogenetic tree of *RAG1(L)* and *RAG2(L)* genes.**

The *RAG1(L)-A/B* and *RAG2(L)-A/B* group definition was based on the literature (Martin et al. 2023, doi: 10.1093/molbev/msad232). JV: jawed vertebrate. Aan: *Aureococcus anophagefferens*; Aau: *Aurelia aurita*; Bbe: *Branchiostoma belcheri*; Bfl: *Branchiostoma floridae*; Bja: *Branchiostoma japonicum*; Bla: *Branchiostoma lanceolatum*; Cmi: *Callorhinchus milii*; Cpu: *Chiloscyllium puntatum*; Cvi: *Crassostrea virginica*; Dre: *Danio rerio*; Gga: *Gallus gallus*; Hsa: *Homo sapiens*; Lch: *Latimeria chalumnae*; Loc: *Lepisosteus oculatus*; Lva: *Lytechinus variegatus*; Mgl: *Marthasterias glacialis*; Mmu: *Mus musculus*; Nve: *Nematostella vectensis*; Oan: *Ornithorhynchus anatinus*; Obr: *Ophioderma brevispina*; Ofa: *Orbicella faveolate*; Ofu: *Owenia fusiformis*; Osp: *Ophiothrix spiculata*; Pfl: *Ptychodera flava*; Spu: *Strongylocentrotus purpuratus*; Xla: *Xenopus laevis*; Xtr: *Xenopus tropicalis*.
